# Assigning confidence to structural annotations from mass spectra with COSMIC

**DOI:** 10.1101/2021.03.18.435634

**Authors:** Martin A. Hoffmann, Louis-Félix Nothias, Marcus Ludwig, Markus Fleischauer, Emily C. Gentry, Michael Witting, Pieter C. Dorrestein, Kai Dührkop, Sebastian Böcker

## Abstract

Untargeted metabolomics experiments rely on spectral libraries for structure annotation, but these libraries are vastly incomplete; *in silico* methods search in structure databases but cannot distinguish between correct and incorrect annotations. As biological interpretation relies on accurate structure annotations, the ability to assign confidence to such annotations is a key outstanding problem. We introduce the COSMIC workflow that combines structure database generation, *in silico* annotation, and a confidence score consisting of kernel density p-value estimation and a Support Vector Machine with enforced directionality of features. In evaluation, COSMIC annotates a substantial number of hits at small false discovery rates, and outperforms spectral library search for this purpose. To demonstrate that COSMIC can annotate structures never reported before, we annotated twelve novel bile acid conjugates; nine structures were confirmed by manual evaluation and two structures using synthetic standards. Second, we annotated and manually evaluated 315 molecular structures in human samples currently absent from the Human Metabolome Database. Third, we applied COSMIC to 17,400 experimental runs and annotated 1,715 structures with high confidence that were absent from spectral libraries.

## 1 Introduction

The discovery and elucidation of novel metabolites and natural products is cost-, time- and labor-intensive; usually, one restricts this work to a handful compounds carefully selected via intricate prior experiments, see e.g. [1, 2]. In contrast, liquid chromatography (LC) coupled to mass spectrometry (MS) allows a relatively comprehensive metabolome analysis of a biological system. LC-MS analysis can detect hundreds to thousands of metabolites from only small amounts of sample; tandem mass spectrometry (MS/MS) individually fragments the observed metabolites and records their fragment masses. Public repositories containing metabolomic LC-MS/MS data^3–5^ are growing quickly, but repurposing this data at a repository scale remains nontrivial.

Structural annotation via MS/MS is usually carried out by spectral library search, but annotations are intrinsically restricted to compounds for which a reference spectrum (usually based on commercially available chemicals) is present in the library. Despite ongoing discussions on how many detected features actually correspond to metabolites^6–8^, it is widely conjectured that a large fraction of compounds remain uncharacterized^9–11^. Beyond establishing a ranking of candidates, the score of the best-scoring candidate in the library (the hit) is used to evaluate the confidence of an annotation: A low hit score indicates that a wrong candidate has been selected, potentially because the correct answer is absent from the library. Evaluation can be carried out using ad hoc score thresholds, or by statistical methods such as false discovery rate (FDR) estimation^12^.

Recently, *in silico* methods were developed that allow to search in substantially more comprehensive molecular structure databases^13–18^, see Online Methods for details. In principle, *in silico* methods can annotate structures not present in all current structure databases, overcoming the boundaries of known (bio-)chemistry: Databases of hypothetical compound structures can be generated combinatorially^19–21^, by modifying existing metabolite structures^22, 23^, or through machine learning^24–26^. Two requirements have to be met by an *in silico* method to be useful for automated annotation of compounds at a repository scale: Firstly, it must not rely on “meta-scores” that integrate information such as citation frequencies or production volumes into the annotation process^27^; this information is clearly not available for hypothetical, novel compounds. Fortunately, CSI:FingerID^15^, best-of-class among *in silico* methods^18^, does not rely on such information. Second, we have to separate correct and incorrect annotations, as it is the case for library search or peptide annotation in shotgun proteomics; this allows us to concentrate downstream analysis on novel compounds most likely to be correctly annotated. Naturally, one may want to use the hit score of an in silico method to differentiate between correct and incorrect hits, as it is done for spectral library search; but unfortunately, separation via hit scores of current *in silico* methods turns out to be impossible.

## Results

### Method overview

Here, we present the COSMIC (Confidence Of Small Molecule IdentifiCations) workflow that combines selection or generation of a structure database, searching in the structure database with CSI:FingerID, and a machine learning-based confidence score to differentiate between correct and incorrect annotations. COSMIC can annotate a substantial fraction of metabolites with high confidence and at low FDR; our evaluations indicate that COSMIC outperforms spectral library search for this purpose, simultaneously expanding the considered compound space. COSMIC can process data at a repository scale, allowing us to repurpose the quickly-growing public metabolomics data. We demonstrate this by processing 20,080 LC-MS/MS datasets using the COSMIC workflow, annotating thousands of features with structures for which at present, no reference MS/MS data are available. Doing so, COSMIC may allow us to flip the metabolomics workflow (Fig. 1): We may concentrate on metabolites annotated with high confidence, without the need for intricate prior experiments, and try to develop a biological hypothesis from these annotations. Annotated fragmentation spectra can subsequently be searched in other datasets via “classical” spectral library search at the repository scale^28^, allowing a more comprehensive annotation of public metabolomics datasets. COSMIC does not require the user to retrain it for individual datasets.

**Fig. 1:**
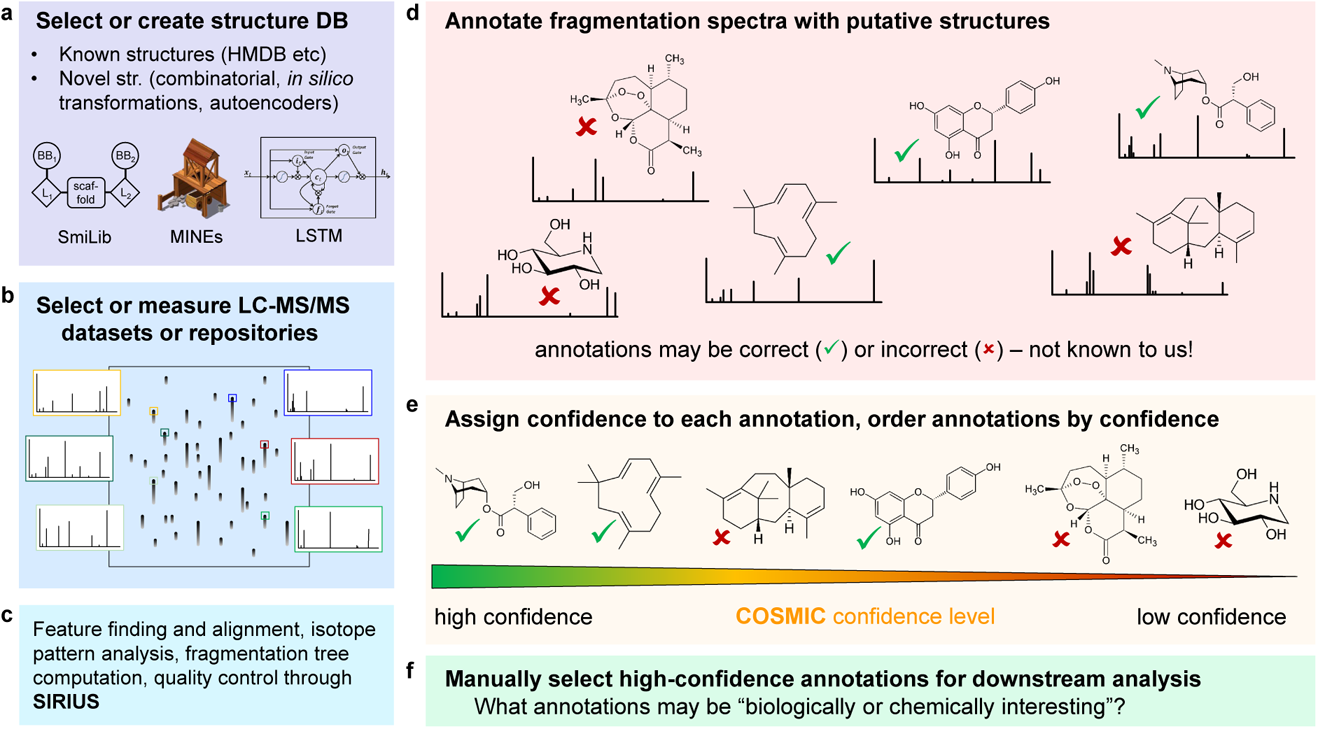
COSMIC workflow. (a) Select or create a structure database; this can be an existing structure database such as the Human Metabolome Database, or generated explicitly for this purpose. (b) Select or measure an LC-MS/MS dataset or select a complete data repository (data repurposing). (c) Data processing through SIRIUS. (d) Structure annotation of fragmentation spectra through CSI:FingerID. (e) Each structure annotation is assigned a confidence score; annotations are then sorted by confidence, allowing users to concentrate on high-confidence annotations. (f) High-confidence annotations can be used to develop or test a biological hypothesis. Notably, COSMIC can annotate metabolites at an early stage of a biological analysis.

### Method evaluation

The highest-ranked candidate for some query fragmentation spectrum is called a hit; it can be either the correct candidate (*correct hit*) or an incorrect candidate (*incorrect hit*). We first demonstrate that one cannot use hit scores of current *in silico* tools to differentiate correct and incorrect hits. We show this for four leading *in silico* tools that participated in the Critical Assessment of Small Molecule Identification (CASMI) 2016 contest^18^, MetFrag^13^, MAGMa+^16^, CFM-ID^14^, and CSI:FingerID^15^. For any reasonable *in silico* method, score distributions of correct and incorrect candidates differ substantially. Yet, the score of the correct candidate competes with all incorrect candidates for this query; by design, there are orders of magnitude more incorrect than correct candidates (Supplementary Fig. 1). Incorrect hits result from either an incorrect candidate receiving a higher score than the correct candidate, or from queries where the correct candidate is missing. Unfortunately, incorrect hits can have high scores, whereas correct hits may exhibit low scores.

See Fig. 2 for CASMI 2016 results. Receiver Operating Characteristic (ROC) curves allow a direct comparison of the separation power of different methods (Fig. 2 d). Area Under Curve (AUC) of ROC curves was between 0.40 and 0.55 for MetFrag, MAGMa+, CFM-ID, and CSI:FingerID, which is not substantially better or even worse than random (AUC 0.5). In comparison, COSMIC reached AUC 0.82. ROC curves ignore the total number of correct annotations of individual methods, so AUCs can be misleading. Here, we introduce hop plots (Methods and Supplementary Fig. 2) allowing us to assess the number of correct hits a method reaches for any given FDR (Fig. 2 ef). Clearly, we are particularly interested in small FDR values. Using COSMIC, we correctly annotated 57 hits with FDR below 10 % searching the biomolecule structure database (124 queries), and 16 hits with FDR 0 % searching ChemSpider^29^ (127 queries). In comparison, MetFrag, MAGMa+ and CFM-ID did not annotate a single compound at FDR 70 % when searching ChemSpider, whereas using the CSI:FingerID score resulted in zero annotations at FDR 40 %. Note that all FDR levels are exact, not estimated (Methods). See Supplementary Fig. 3 for CSI:FingerID results without structure-disjoint evaluation.

**Fig. 2:**
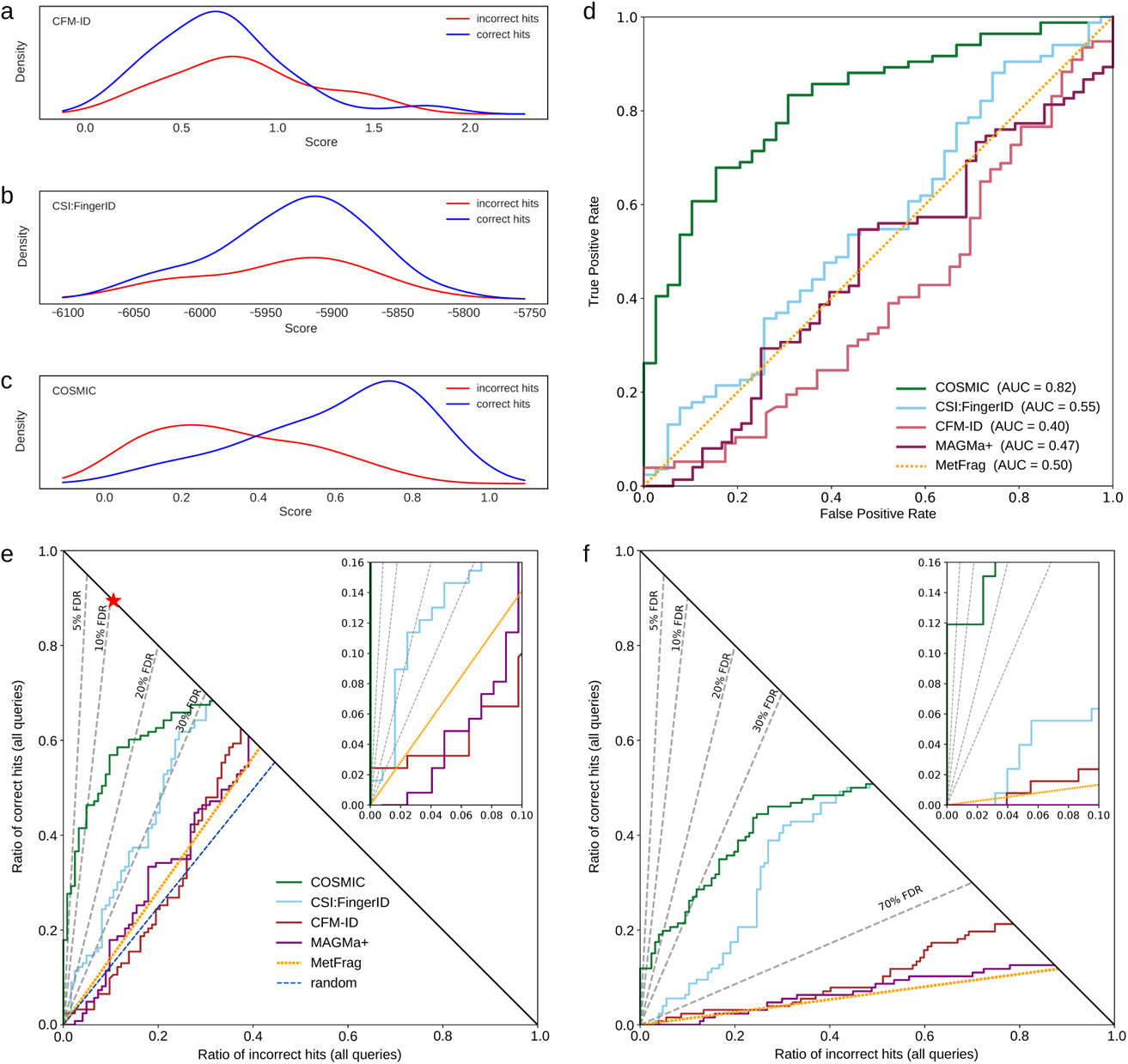
Separation by hit score for different *in silico* tools, using the CASMI 2016 contest submissions. Positive ion mode, candidates retrieved by molecular formula. (a–e) Searching the biomolecule structure database, *N* 124 queries; (f) searching in ChemSpider, *N* 127 queries. (a–c) Kernel density estimates of the score mixture distribution (correct and incorrect hits) for (a) CFM-ID, (b) CSI:FingerID, ensuring structure-disjoint training data through cross-validation, and (c) COSMIC. Kernel density estimates do not allow for a direct comparison of different tools. (d) Receiver Operating Characteristic curves for MetFrag, MAGMa+, CFM-ID, CSI:FingerID (ensuring structure-disjoint training data), and COSMIC. MetFrag normalizes scores, so the ordering of hits is exactly random. (e,f) Hop plots for the same tools, searching (e) the biomolecule structure database or (f) ChemSpider. FDR levels shown as dashed lines; FDR levels are exact, not estimated (Methods). The blue dashed line in (e) indicates random scores, resulting in random ordering of candidates and hits; the red star in (e) is the best possible search result. (a–f) CSI:FingerID and COSMIC computed here; all other scores from [18].

The CASMI 2016 dataset is comparatively small and prone to stochastic fluctuations; hence, we thoroughly evaluated COSMIC using two large datasets: We used tenfold cross-validation on the training data, and an independent reference dataset with 3,291 compounds (forensics/toxicology library, Agilent). We must use reference datasets for evaluation, as the correct answer is usually unknown for biological datasets. Unless indicated otherwise, all evaluations were again carried out structure-disjoint. In biological experiments, fragmentation spectra are often recorded at exactly one collision energy; to this end, we compiled three spectral libraries for individual collision energies, plus one with merged spectra from three collision energies. Furthermore, fragmentation spectra from biological samples seldom reach the same quality as reference spectra; to this end, we added (medium or high) noise to the reference spectra before evaluation. We search in a biomolecule structure database combined from several public databases. For 22.70 % of the queries in cross-validation and 17.29 % from independent data, the correct answer was missing from the searched structure database. Hence, COSMIC cannot correctly annotate these compounds; we did not discard these queries as in application, the correct structure may indeed be unavailable. Added noise, single collision energy spectra and unsolvable instances result in CSI:FingerID annotation rates (34.91 % to 49.15 % correct hits in cross-validation, 41.80 % to 59.98 % on independent data) substantially smaller than those previously reported^15, 30^.

COSMIC calibrates CSI:FingerID scores using E-value estimates. We used candidates in PubChem^31^ as a proxy of decoys, and empirically established that CSI:FingerID scores can be modeled as a mixture distribution of log-normal distributions (Supplementary Fig. 4). COSMIC computes E-values using the kernel density estimate of the score distribution. The calibrated score showed slightly better separation than the CSI:FingerID score (Fig. 3). The COSMIC confidence score uses a Support Vector Machine (SVM) to classify whether a hit is correct. To lower chances of overfitting, we trained a linear SVM and enforced directionality of features. For instances with only a single candidate structure, we trained an additional SVM. Besides the calibrated score, which turned out to be a highly important feature in particular if the candidate list contains only one candidate, COSMIC uses features such as score differences to other candidates and the length of the molecular fingerprints (Supplementary Table 1). Classification results were mapped to posterior probabilities using Platt estimates^32^. We used tenfold cross-validation for training and evaluation of the SVM, and ensured structure-disjoint evaluation for independent data. Separation between correct and incorrect hits is much stronger than for (calibrated) scores, see Fig. 3. We found that mass had no pronounced impact on separation via the confidence score; in contrast, the number of candidates in the structure database had a strong impact (Supplementary Fig. 5 and 6).

**Fig. 3:**
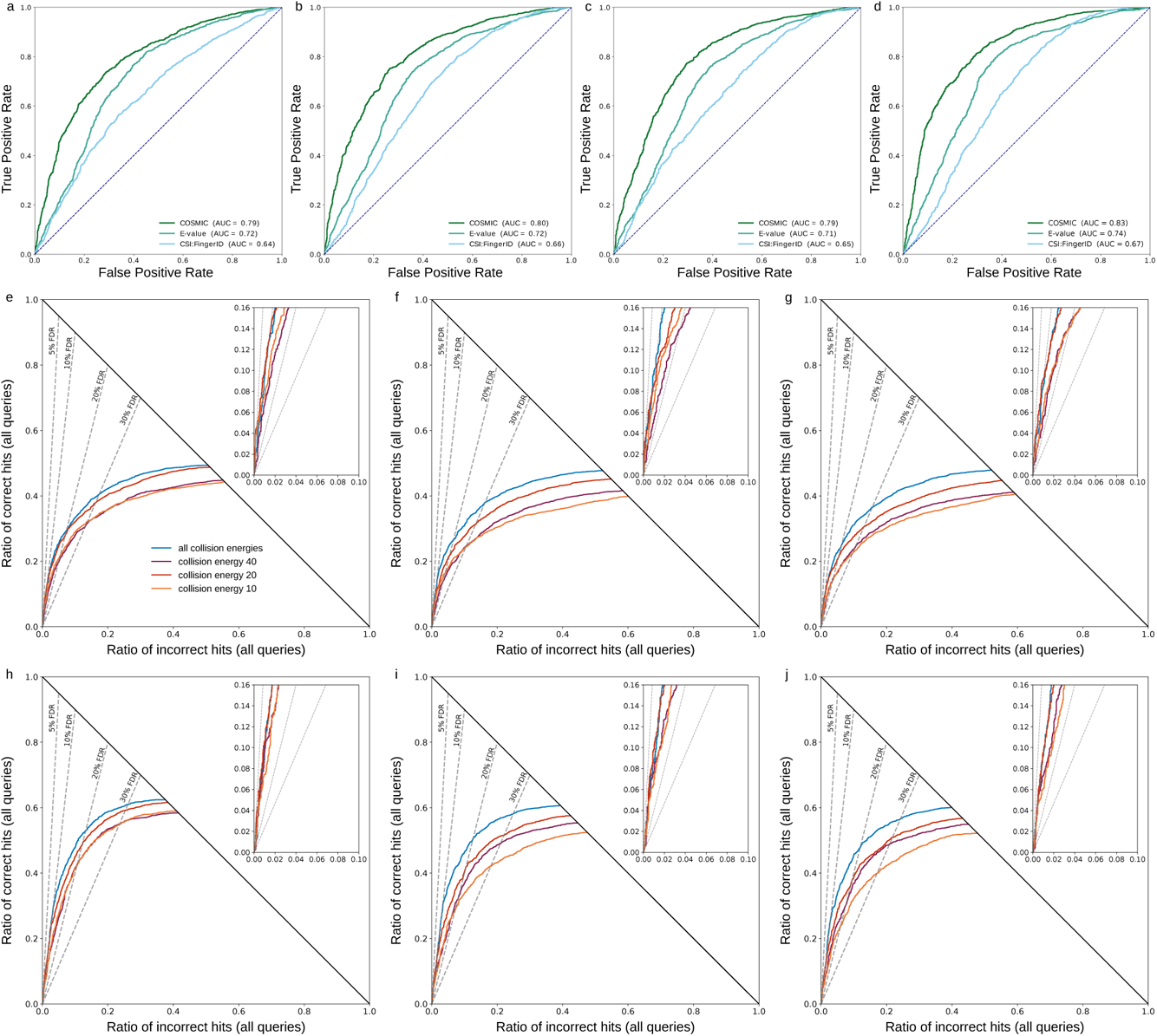
Evaluation of separation searching in the biomolecule structure database. (a–d) Comparison of CSI:FingerID score, calibrated score (E-value) and COSMIC confidence score. ROC curves, structure-disjoint evaluation, independent data, medium noise, *N* = 3, 013. (a) 10 eV, (b) 20 eV, (c) 40 eV, (d) merged spectra (“all collision energies”). In each plot, all curves end in the same number of correct hits (1,829 for (a), 1,901 for (b), 1,765 for (c), 1,948 for (d), so a hop plot would not contain additional information. (e–j) Evaluation of COSMIC confidence score: Hop plots for different collision energies. (e–g) Structure-disjoint cross-validation, *N* = 3, 721. (h–j) Independent data with structure-disjoint evaluation, *N* = 3, 013. (e,h) No added noise, (f,i) medium noise, (g,j) high noise. FDR levels shown as dashed lines; FDR levels are exact, not estimated (Methods).

Inevitably, some incorrect hits received a high confidence score and, hence, would be wrongly regarded as “probably correct”. Fig. 4 shows the nine incorrect hits with highest confidence scores when searching independent data with medium noise. In seven of nine cases, the correct structure was not contained in the biomolecule structure database. In all nine cases, the correct structure was highly similar to the corresponding hit; in contrast, the bottom nine incorrect hits generally showed little structural similarity to the corresponding correct structures (Supplementary Fig. 7). We then compared fragmentation spectra for three structure pairs from Fig. 4; for each pair, fragmentation spectra are indeed highly similar (Supplementary Fig. 8) with cosine score between 0.85 and 1.00. Hence, spectral library search may result in the same high-confidence misannotations.

**Fig. 4:**
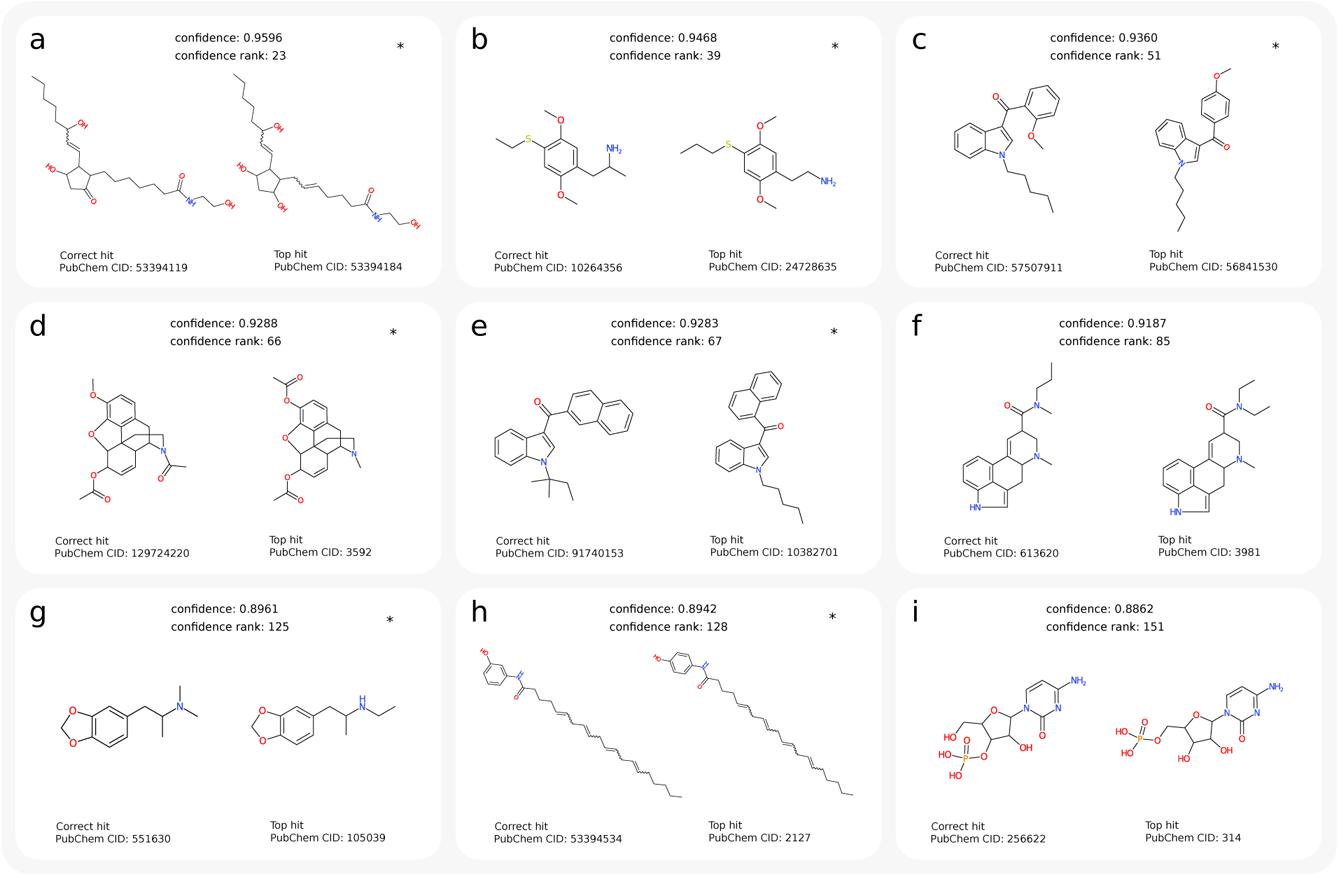
Examples of incorrect annotations with highest confidence scores. Queries are cross-validation data, merged spectra, medium noise, biomolecule structure database, structure-disjoint evaluation. (a–i) Incorrect hits (incorrect top-ranked structures) with highest confidence scores. Top-ranked structure on the left and corresponding correct structure on the right. “PubChem CID” is PubChem compound identifier number. “Confidence rank” indicates the rank of the incorrect hit in the complete list of hits: There exist 22 correct hits before the first incorrect hit, all other 142 hits with confidence score above 0.8862 were correct, and only 8 out of 150 top-ranked hits (5.33 %) are incorrect. Instances where the correct structure was not contained in the biomolecule structure database are marked by an asterisk. For these instances, a correct annotation by CSI:FingerID is impossible; at the same time, it is highly challenging for COSMIC to identify these hits as “incorrect”. In seven cases, molecular graphs of the incorrect hit and correct structure differ by the theoretical minimum of two edge deletions. Query spectra: (a) NIST 1210761/62/64, (b) NIST 1617825/29/34, (c) NIST 1320583/85/91, (d) NIST 1429464/65/71, (e) NIST 1483460/63/69, (f) NIST 1247455/57/63, (g) NIST 1480825/30/34, (h) NIST 1418771/73/80, (i) NIST 1276453/55/59.

FDR estimation for small molecule annotation is highly challenging^12, 33^; this is an intrinsic problem of small molecule annotation, as the assumption of incorrect hits being random is fundamentally violated^34^. We transferred COSMIC confidence scores to FDR estimates^12, 35^ but as expected, these estimates were of mediocre quality only (Supplementary Fig. 9). In particular, estimates for independent data were highly conservative: estimated q-values were much larger than true q-values. Consequently, confidence score values must be treated as a score, but not as the probability that the annotation is correct.

We also trained classifiers for searching in PubChem instead of the biomolecule structure database. These classifiers showed a worse performance (Supplementary Fig. 10), and we observed a substantial drop of correct annotations for small FDR values. Again, COSMIC strongly outperformed both E-values and the CSI:FingerID score.

### Evaluation against spectral library search

We also evaluated the dereplication power of COSMIC in comparison to spectral library search. One may expect that targeting novel compounds (the true purpose of the COSMIC workflow) instead of dereplication comes at a price: The biomolecule structure database is more than an order of magnitude larger than GNPS^37^ and NIST spectral libraries, and we cannot rely on direct spectral comparison. Somewhat unexpectedly, COSMIC annotated substantially more compounds for all reasonable FDR rates, see Fig. 5: At FDR 5 %, COSMIC outperformed library search 1,415 to 52 hits at 20 eV and 1701 to 1 hits using merged spectra, respectively. Notably, COSMIC correctly annotated compounds with high confidence although query spectrum and reference spectrum were (highly) dissimilar, with cosine scores between 0.06 and 0.63 (Supplementary Fig. 11).

**Fig. 5:**
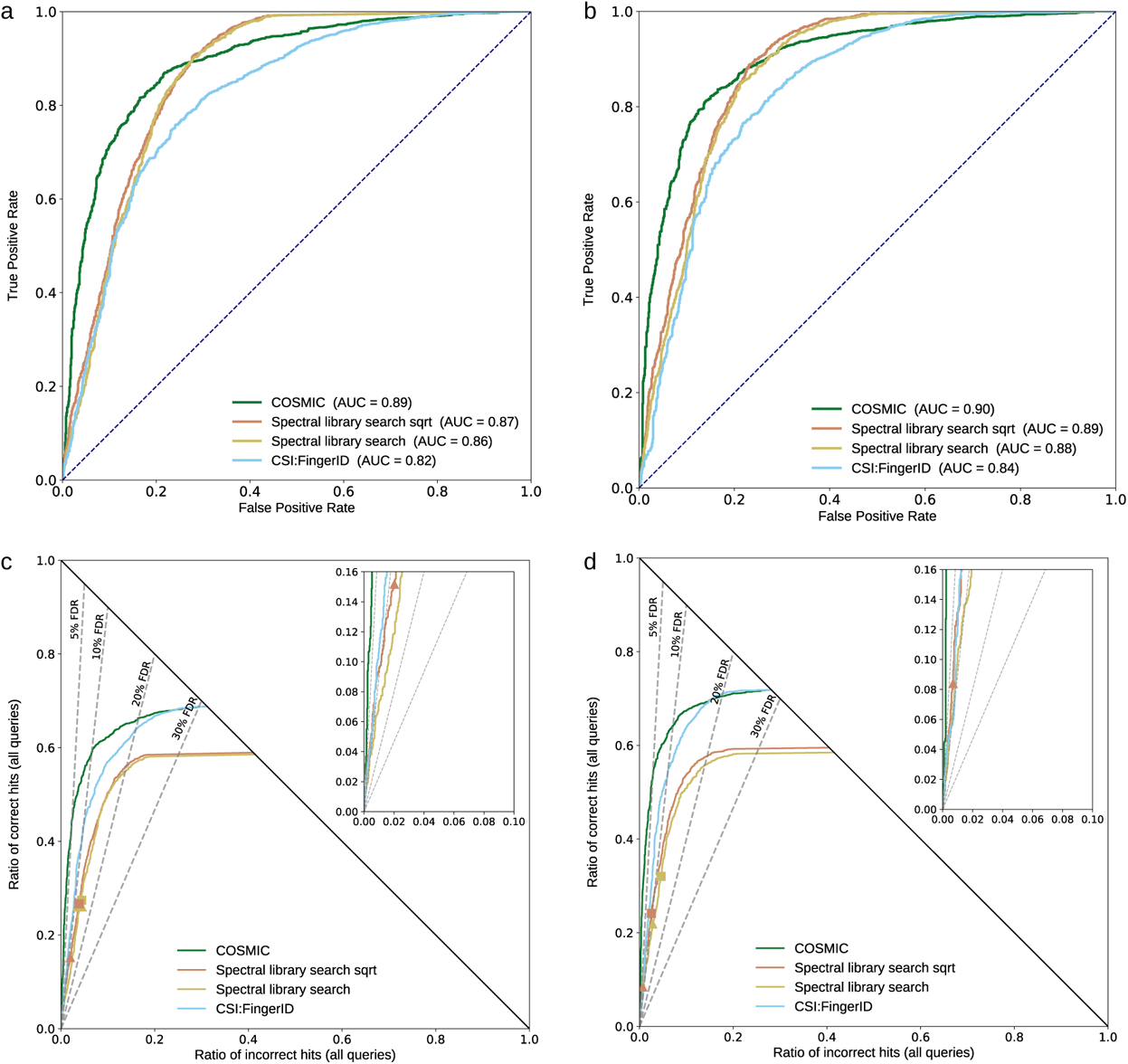
Comparison to spectral library search and separation without structure-disjoint evaluation. Query spectra (independent dataset) distorted with medium noise; COSMIC is searching the biomolecule structure database. ROC curves (a+b) and hop plots (c+d) for collision energy 20 eV (a+c) and merged spectra (b+d). There is no overlap in fragmentation spectra between training data and independent data, but we do not remove training data for which we find the same structure in the independent dataset. To this end, 2,192 of the *N* = 3, 013 structures from the independent dataset (72.75 %) are also present in the spectral library. We compare search performance and separation of COSMIC, the CSI:FingerID score and spectral library search. All three methods utilize basically the same MS/MS data. For spectral library search, we compute the normalized dot product using either regular peak intensities, or the square root of peak intensities (“Spectral library search sqrt”)^36^. Spectral library search candidates were restricted to those with the correct molecular formula for each query. The spectral library is 16-fold smaller than the biomolecule structure database, giving library search a large competitive edge in evaluation. Notably, COSMIC results in substantially more correct annotations than library searching for all reasonable FDR levels; FDR levels are exact, not estimated (Methods). For spectral library search, markers show commonly used cosine score thresholds 0.9 (triangle) and 0.8 (square), respectively.

### Searching for novel bile acids conjugates

COSMIC allows us to expand structure annotation beyond the space of known molecules, making it possible to explore novel (bio)chemical processes: To demonstrate this, we used COSMIC to search for novel bile acid conjugates. Recently, a fifth mechanism of the bile-acid metabolism by the microbiome was discovered in mice/humans^38^. In that study, three novel bile acid conjugates with phenylalanine, tyrosine, and leucine were found. This finding supports the possibility that other bile acids conjugated with different amino acids could exist (taurocholic and glycocholic acids are the two other known historically).

We explored this hypothesis applying COSMIC to a public mice fecal metabolomics dataset. Plausible bile acid conjugate structures were computed by combinatorially adding amino acids to bile acid cores, yielding 28,630 plausible bile acid conjugates. The COSMIC workflow was then applied using both PubChem and the combinatorial bile acid conjugate database. CSI:FingerID annotations were ranked by COSMIC confidence score. To establish which of the annotated structures were “truly novel” (Supplementary Table 2, Supplementary Material 3), the structures were searched in PubChem and known structures were discarded; the top 12 best-scoring “truly novel” structures proposed by COSMIC (Fig. 6) were validated by manual interpretation of the fragmentation spectra (Supplementary Fig. 12 to 23). In addition, we synthesized two annotated structures to validate annotations of phenylalanine-conjugated chenodeoxycholic acid (Phe-CDCA, **7** in Fig. 6) and tryptophan-conjugated chenodeoxycholic acid (Trp-CDCA, **12**), see Supplementary Fig. 24.

**Fig. 6:**
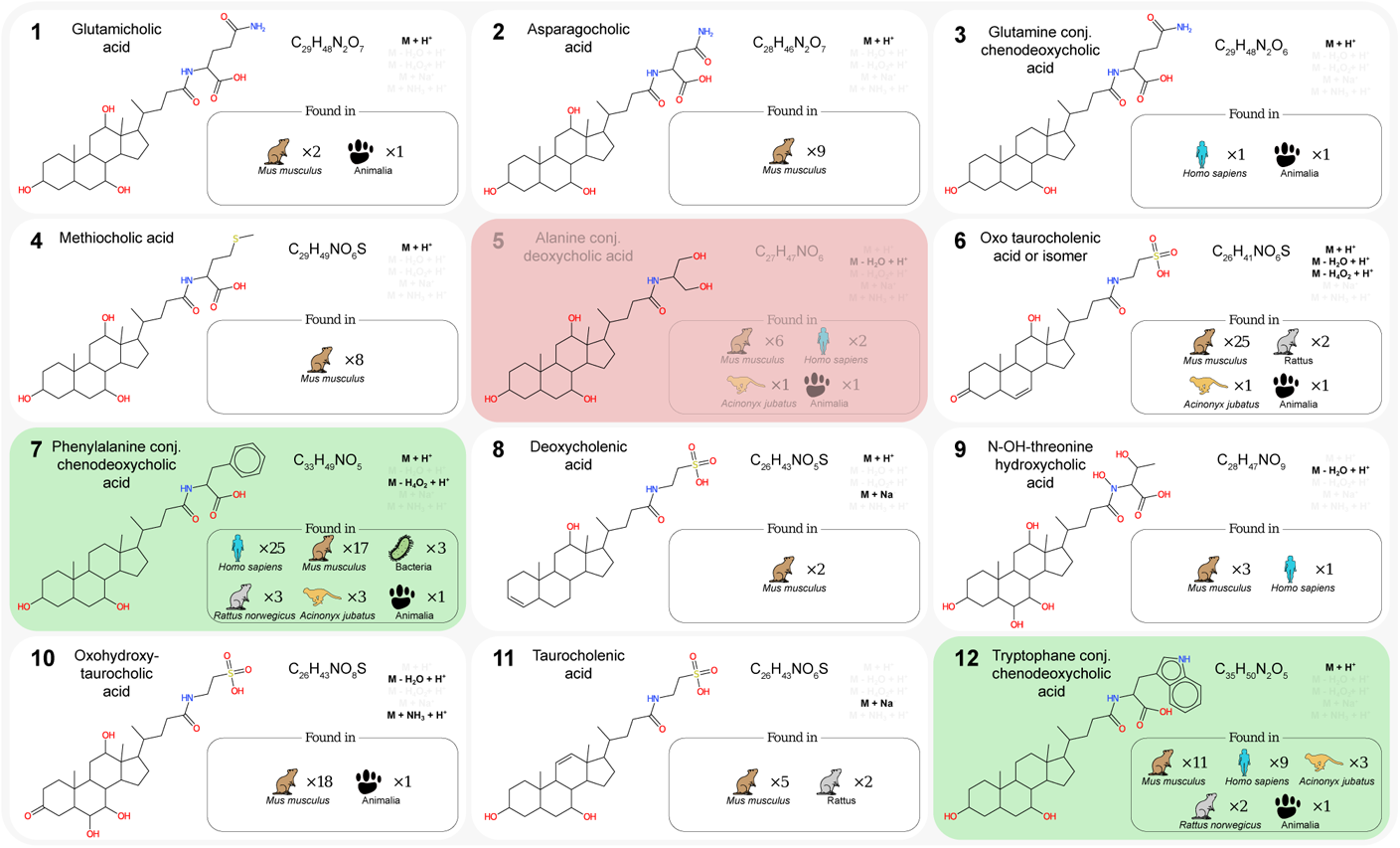
Top 12 highest-scoring COSMIC annotations of “truly novel” bile acid conjugates. Bile acid conjugates which are also present in PubChem are omitted from the list; see Supplementary Table 2 for the complete list. For each bile acid conjugate we report its chemical name, putative structure, molecular formula, and adducts. We also report species and number of datasets with spectral matches from a MASST search. Two annotations verified by authentic standards are highlighted in green, the single incorrect annotation in red.

Inspection of the fragmentation spectra showed characteristic fragment ions for the bile acid core structure: *m/z* 337.2526 and *m/z* 319.2420 for cholic acid derivatives, and *m/z* 339.2682 and *m/z* 321.2577 for a dehydroxylated bile acid core structure, presumably chenodeoxycholic acid (CDCA) given the validation for **7** and **12**. The nature of the amino acid residue was confirmed from the observation of the specific amino acid fragments. First, we observed novel bile acids derivatives conjugated for the newly discovered conjugation^38^ with phenylalanine (*m/z* 166.0863, Phe-CDCA, **7**). Most significantly, COSMIC enabled the discovery of completely novel amino acids bile acid conjugations. This includes bile acids conjugated with glutamine (fragment *m/z* 147.0764, glutamicholic acid **1**, and Glu-CDCA **3**), asparagine (fragment *m/z* 133.0608, asparagocholic acid, **2**), methionine (fragment *m/z* 150.0583, methiocholic acid, **4**), and tryptophan (fragment *m/z* 205.0972, Trp-CDCA, **12**). In addition, a bile acid conjugated with a non-canonical amino acid was annotated (N-OH threonine, fragment *m/z* 134.0440, **9**). Other annotated derivatives had modified bile acid cores, including dehydration/reduction/oxidation and were supported by the analysis of the fragmentation pattern as for putative oxotaurocholenic acid (**6**), deoxycholenic (**8**), oxohydroxytaurocholic acid (**10**), taurocholenic acid (**11**). A single COSMIC annotation was incorrect (**5**): Inspection revealed that it was wrongly interpreted as an in-source fragment. Yet, the expected fragment for a serinol-conjugated bile acid was not observed (Supplementary Fig. 16) and further interpretation of the fragmentation spectrum supported that it was likely the protonated ion of the alanine-conjugated CDCA.

Molecular networking analysis (Fig. 7) showed that the validated annotations (**7** and **12**) were part of a molecular network including leucine-conjugated CDCA as well as other CDCA conjugates annotated by COSMIC, but not among the top 12; namely methionine-conjugated CDCA and tyrosine-conjugated CDCA. Inspection of corresponding fragmentation spectra showed the COSMIC annotations were consistent (Supplementary Figs. 25 and 26). The relative abundance of these bile acid conjugates were predominantly observed in the mice group the high-fat diet. Previous research showed an increase in cholic acid and deoxycholic secretion in mice subject to a high-fat diet^39^ and could explain the higher abundance and the observation of novel bile acid conjugation.

**Fig. 7:**
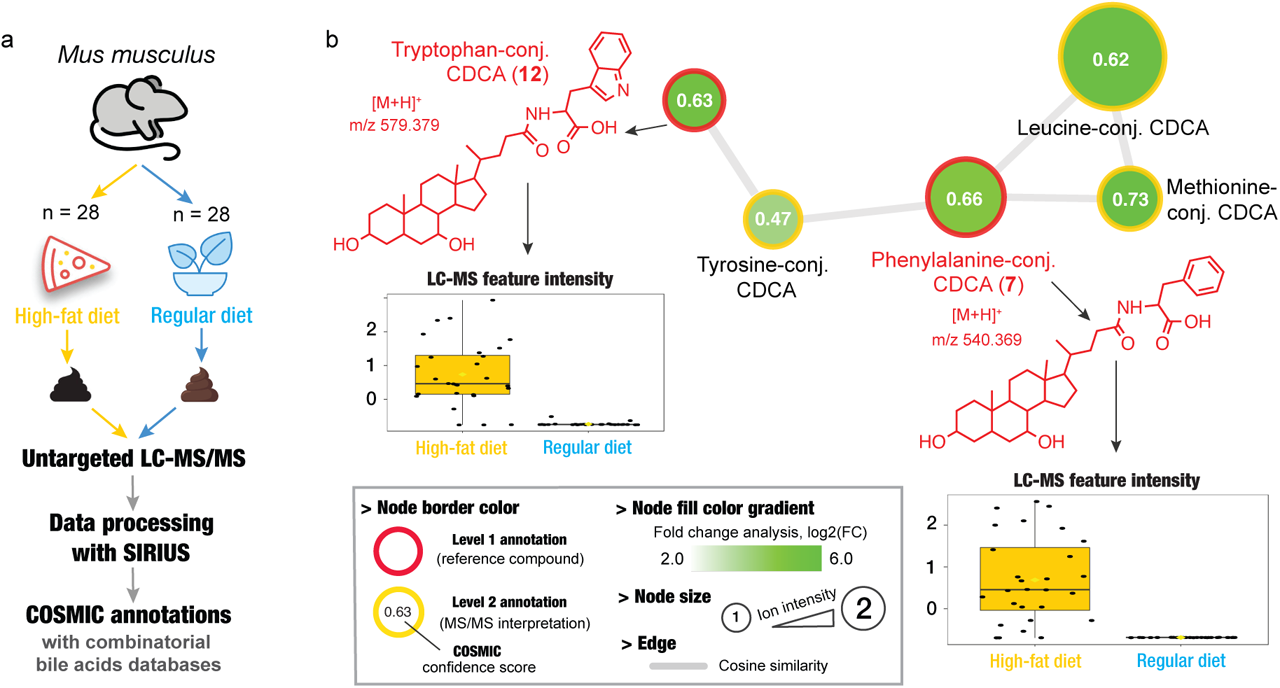
Applying COSMIC to discover novel bile acids conjugates in a mice fecal dataset. (a) Experimental design and the data processing and annotation carried out with COSMIC. (b) Mass spectrometry-based molecular network of novel bile acid conjugates annotated with COSMIC and the combinatorial bile acids structure database. Two annotations (**7** and **12**) were verified against synthetic standards and the other annotations were manually inspected. Fold change analysis showed that all these bile derivatives were predominantly observed in mice fed with a high-fat diet.

The novel bile acids were searched in all the public mass spectrometry data repositories^37^ by performing a MASST search^28^. Matching fragmentation spectra were observed in public datasets (Fig. 6), predominantly consisting of data from animal fecal samples, mostly from rodents and humans. Interestingly, a match for Phe-CDCA (**7**) was observed in a bacterial culture of two opportunistic pathogens (*Escherichia coli* and genus *Stenotrophomonas*) and resonates with previous findings on the fifth mechanism of bile acid metabolism by the microbiome^38^.

### Repository-scale annotation of novel metabolites

The Human Metabolome Database (HMDB)^40^ contains the by far most comprehensive collection of molecular structures found in or on the human body, with version 4.0 embracing 114,265 structures. Yet, certain molecular structures connected to human metabolism may currently be missing from this database. To test this hypothesis, we searched the human dataset against the biomolecule structure database; this comprises ten MassIVE datasets^37^ with 2,666 LC-MS/MS runs from different sources (serum, plasma, lips, tongue, teeth, fecal, urine). We concentrated on those hits with structures absent from HMDB. This resulted in 436 high-confidence structure annotations; 121 (27.8 %) of the structures were present in our MS/MS training data, leaving us with 315 structures for which no MS/MS reference data are available (Supplementary Fig. 27, Supplementary Table 4). The HMDB database used for excluding structures dates back to Aug 2018; since then, at least 26 of these structures were added to HMDB. This indicates that many of the novel structures are indeed present in human samples. We manually validated the 315 structures, of which 48 are proteinogenic peptides (peptides made from proteinogenic amino acids), by checking common neutral losses and fragments, and by comparison of spectra against reference spectra from similar compounds. Based on characteristic fragmentation patterns, different acyl-carnitines and N-acyl-amino acids not part of the HMDB were annotated. N-acyl amino acids are well known uncoupling agents in mitochondria^41^. From 30 spectra annotated as acyl-carnitines with high confidence, 21 were presumably correct based on manual validation. N-oleyl-leucine represents one particular example of an N-acyl amino acid annotated with high confidence; the annotation was verified using a reference spectrum that was not part of the COSMIC or CSI:FingerID training data (Supplementary Fig. 28). A MASST search^28^ in GNPS gave 84 datasets putative containing a similar spectrum, 38 being human datasets. For two additional high confidence hits, reference spectra were available and showed high similarity to the query spectra (Supplementary Fig. 28). Hits are available from https://bio.informatik.uni-jena.de/cosmic/; users can view, discuss and validate annotated structures there.

To further demonstrate COSMIC’s power to annotate metabolites at a repository scale, we then searched the Orbitrap dataset consisting of 123 MassIVE datasets^37^ and 17,414 LC-MS/MS runs (Supplementary Table 5) against the biomolecule structure database. This resulted in 3,530 metabolite structures annotated with high-confidence, of which 1,815 were present in the training data. Discarding those, we are left with 1,715 novel structure annotations (Supplementary Fig. 29, Supplementary Table 6); for comparison, the data used to train CSI:FingerID and COSMIC comprises 16,703 structures. Again, hits are available from https://bio.informatik.uni-jena.de/cosmic/.

## Discussion

Annotation scores of current *in silico* tools are not suited to separate correct from incorrect hits. Here, we have introduced the COSMIC workflow that assigns confidence scores to structure annotations. Annotation rates as well as separation were consistently better when using merged spectra, usually followed by 40 eV fragmentation spectra. COSMIC clearly outperformed spectral library search for dereplication; this is notable as COSMIC has not been designed or optimized for this task. Be reminded that in our evaluations, only the exact structure was regarded as correct; yet, small structure modifications (Fig. 4) are hard and potentially impossible to tell apart using MS/MS data alone. This is an intrinsic limitation not of COSMIC but of small molecule MS/MS in general, and requires orthogonal information to overcome; yet, these incorrect annotations often contain viable structure information. Indeed, COSMIC’s incorrect annotations with high confidence are often structurally highly similar to the correct structure; again, COSMIC has been neither designed nor optimized for finding structurally similar compounds.

We demonstrated that COSMIC can be used to search for novel metabolites and rapidly test biological hypotheses. More specifically, we found additional amino acid conjugation of bile acids beyond those previously identified by repurposing public datasets^38^, opening the gate for studying their precise structure and biological relevance. Notably, more than 90 % of the annotations for the top 12 bile acid conjugates suggested by COSMIC turned out to be correct. These annotations will help to further explore bile acid metabolism: Only the glycine and taurine bile acid conjugates were previously known in humans, despite 170 years of research in that field^42^.

We further demonstrated COSMIC’s power by repurposing data from 20,080 LC-MS/MS runs, providing high-confidence hits in a biomolecule structure database; for 49 % of these hits, no reference spectra were available in our training data. In particular, we annotated 267 metabolites in human datasets absent from HMDB with no reference MS/MS data available, compared to 108 such metabolites with reference MS/MS data. Annotations may now serve as starting points for generating biological hypotheses, or expand existing spectral libraries.

Notably, COSMIC complements compound class annotation tools such as CANOPUS^43^: COSMIC targets molecular structure annotations but annotates only a fraction of the compounds in a sample; in contrast, CANOPUS annotates practically all compounds in a sample for which fragmentation spectra have been measured, but is restricted to annotating compound classes. Hence, both methods provide viable information; which method is better suited, depends on the underlying research question.

The COSMIC score must not be mistaken as the probability that an annotation is correct; this is impossible by design of the score. We conjecture that accurate FDR estimation from fragmentation spectra of small molecules will remain highly challenging.

## Methods

### General considerations

Establishing the stereochemistry from fragmentation spectra is highly challenging and beyond the power of automated search engines; hence, only the two-dimensional structure is considered when evaluating a hit structure: we consider the identity and connectivity (with bond multiplicities) of the atoms, but ignore the stereo-configuration for asymmetric centers and double bonds.

The term “novel compound” has previously been used to describe conflicting and imprecisely defined concepts, such as when an unexpected compound is detected in a sample or organism, or whether compounds have previously been described in the literature. Throughout this paper, a structure is considered “novel” if no MS/MS data from a compound with the same structure are present in the training data; hence, the compound cannot be annotated through spectral library search. We noted above that stereoisomers (compounds with identical structure, such as L-threose, D-threose, L-erythrose and D-erythrose) show highly similar fragmentation. Hence, for L-threose to be novel, the training data must not contain MS/MS data for L-threose, D-threose, or (L- or D-)erythrose. In our evaluations, we ensure that all compounds are novel using structure-disjoint cross-validation.

Similarly, a “truly novel” compound refers to a compound structure absent from large public databases such as PubChem^31^ or ChemSpider^29^; quotation marks are in place as the (non-public) database GDB-17^44^ contains 166 billion hypothetical structures of small molecules, and “truly novel” compounds may already be in there. For CSI:FingerID and other *in silico* methods that do not rely on meta-scores, there is no difference to search in a database of “truly novel” hypothetical structures, or to search in PubChem or the biomolecule structure database. It is understood that correct annotation rates will deteriorate if the database we search in becomes too large.

COSMIC targets *biomolecules*, that is, products of nature as well as synthetic products with potential bioactivity, including drugs, toxins, food, cosmetics and other xenobiotics. This restriction of focus is due to the available MS/MS training data.

### False discovery rates

Given a list of hits, the *false discovery rate* (FDR) of this list is the number of correct hits in the list, divided by the size of the list. Hence, to compute FDR, we must know the exact number of correct and incorrect hits. Throughout this paper, evaluations were carried out using reference data so that the true structure underlying any query spectrum was *unknown to the method but known to us*. To this end, *all reported FDR rates are exact*, unless indicated otherwise. At this point, there is no need to employ methods for FDR estimation^12, 33, 45, 46^; such methods try to accurately estimate the true FDR in application, where we do not have knowledge of correct and incorrect hits. Accurate FDR estimation remains a highly non-trivial problem in general statistics as well as many fields of application, see also below.

### *In silico* methods and related work

So-called “*in silico* methods” allow us to search in a molecular structure database using MS/MS data as our query. Most methods follow one of three paradigms: (i) Combinatorial fragmenters^13, 47–49^ try to explain the query spectrum using the candidate structure, combinatorially breaking bonds in the molecular structure graph. (ii) Other methods try to predict the fragmentation spectrum of a given compound structure^14, 50, 51^; this allows us to search in the structure database by spectral matching. (iii) Alternatively, we can transform the query spectrum into information about the query structure, then use this structure information to search in the structure database^15, 52, 53^. Later publications basically present minor modifications of these ideas; an exception are the Input Output Kernel Regression variants of CSI:FingerID^17, 54^, which use molecular fingerprints but circumvent the prediction of individual molecular properties, instead predicting similarity of a candidate to the query by regression.

Some methods use so-called “metascores” that integrate information about citation frequencies or production volume^48, 55, 56^. We stress that “metascores” have nothing in common with “metadata”, except for the prefix; metadata is information about the experimental setup and the biological sample, whereas these metascores use side information unrelated to the actual experiment. These metascores usually perform well in evaluations but come with several severe restrictions; in the context discussed here, the most important restriction is that the above side information is not available, and metascores therefore not applicable, for any “truly novel” structure such as novel bile acid conjugates. Furthermore, metascores tend to prefer highly cited “blockbuster metabolite” candidates; hence, evaluation results, which are carried out using mainly such “blockbuster metabolites”, are often exaggerated. Similar limitations are associated with metascores based on taxonomy^57^ as again, this information is not available for “truly novel” structures. Thus, we ignored metascore methods in our evaluations.

Finally, some tools use networks for structure annotations; networks may be based on spectral similarity in the LC-MS/MS run, or structural similarity in the metabolite database^57–60^.

### Structure databases

Different from previous studies^15, 30^ where structures were derived from InChI (International Chemical Identifier) strings, molecular structures were standardized using the PubChem standardization procedure^31^. In particular, a canonical tautomeric form was chosen, as solvent, temperature, and pH in the sample influence the dominating tautomeric species. Standardization of compounds not in

PubChem was carried out using the web service at https://pubchem.ncbi.nlm.nih.gov/rest/pug/. Unfortunately, PubChem standardization has changed multiple times over the last years without further noticing of users; to this end, it is possible that some non-PubChem compounds were standardized slightly differently than structures from the MS/MS training data.

We searched in the following structure databases with COSMIC:

– For the CASMI 2016 evaluation^18^, we downloaded structures from the CASMI 2016 results web page (http://casmi-contest.org/2016/). Candidate structures were provided as part of the blind contest and originally retrieved from ChemSpider^29^.
– The *biomolecule structure database* is a union of several public structure database including HMDB^40^, ChEBI^61^, KEGG^62, 63^ and UNPD^64^. The resulting database contains 391,855 unique structures of biomolecules and compounds that can be expected to be present in biological samples.
– The *HMDB structure database*^40^ was downloaded Aug 8, 2018 and contains 113,983 compounds, and 95,980 unique structures with mass up to 2000 Da.
– The *PubChem structure database*^31^ was downloaded Jan 16, 2019, and contains 97,168,905 compounds, and 77,153,182 unique covalently-bonded structures with mass up to 2000 Da. We added all missing structures from the biomolecule structure database, which resulted in a total of 77,190,484 unique structures.
– A combinatorial database of 28,630 *bile acid conjugate structures* was generated with SmiLib v2.0^19, 20^, downloaded from http://melolab.org/smilib/. SmiLib generates chemical structures by combining scaffolds and building blocks provided as SMILES (Simplified Molecular Input Line Entry Specification). We curated a list of initial bile acid “scaffolds” that represent common steroid cores (i.e. cholic acid, deoxycholic acid, hyocholic acid, chenodeoxycholic acid). Initial scaffolds were modified manually with common phase II metabolism reactions (i.e. glucuronidation, acetylation, sulphation, methylation) and resulted in 322 scaffolds. To generate bile acid conjugates, scaffolds were combined with 91 building blocks, including proteinogenic and non-proteinogenic amino acids, along with their N-hydroxylated and N-methylated version, and acyls moieties. Stereochemical information was removed prior to the database generation with SmiLib.

### MS/MS reference datasets and noise addition

For evaluations, we limited ourselves to MS/MS spectra recorded in positive ion mode, as there are generally more such spectra available. This is not a restriction of COSMIC, which can also be trained on negative ion mode data. Evaluations were carried out using reference measurements, as we do not know the correct answers for biological datasets.

For the CASMI 2016 evaluation, MS/MS spectra were downloaded from the CASMI web page (http://casmi-contest.org/2016/). Spectra from different collision energies were merged prior to analysis.

To train CSI:FingerID, we used a combined dataset from MassBank^65^, GNPS^37^ and the NIST 2017 database (National Institute of Standards and Technology, v17). Reference MS/MS data were measured on different high-resolution instruments from multiple vendors. The *CSI training dataset* contains 16,703 structures with 23,965 independent MS/MS measurements. As an independent dataset, we used the commercial MassHunter Forensics/Toxicology PCDL library (Agilent Technologies, Inc.) with 3,243 structures and 3,462 independent MS/MS measurements, all measured on an Agilent Q-ToF instrument. Unlike the commercially available library, these mass spectra were not curated. When discussing reference dataset evaluations, independent MS/MS measurements will be referred to as “compounds” for the sake of brevity.

For one compound, a library usually contains fragmentation spectra measured at different collision energies. Different from the CSI training dataset as well as previous evaluations^15, 30^, we did not directly merge fragmentation spectra. Instead, we compiled fragmentation spectra sets for both training and independent data using single collision energies, namely, 10 eV, 20 eV and 40 eV. In many cases, libraries did not contain fragmentation spectra for exactly those collision energies; we allowed for a deviation of up to 4 eV, and in case fragmentation spectra for more than one collision energy were present in this interval, we used the one with collision energy closest to the desired one. Finally, *merged spectra* were generated by combining these three spectra (pseudo-ramp spectra).

Fragmentation spectra in reference libraries often have much better quality (more signal peaks, fewer noise peaks, better signal-to-noise) than fragmentation spectra from a biological LC-MS/MS run. To simulate this effect in our reference datasets, we “added noise” to each fragmentation spectrum. Distorting spectra followed comparable principles as the generation of decoy spectra^12^: We distorted spectra similar to what we expect for experimental spectra. For example, adding noise peaks with (uniform) random mass will result in spectra which are notably different from experimental ones^12^. We simulated two noise models, *medium noise* and *high noise*.

– We simulated a *global mass shift* (bias) by drawing a random number *δ*\* from 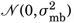, then shifting every peak mass *m* by *δ* m*. The standard deviation *σ*_mb_ was chosen as *σ*_mb_ = (10/3) *·* 10^−6^ (medium noise) or *σ*_mb_ = (15/3) *·* 10^−6^ (high noise), so that the 3*σ*_mb_ interval represents a 10 ppm shift for medium noise and a 15 ppm shift for high noise.
– We simulated individual *mass deviations* by drawing, for each peak with mass *m* individually, a random number *δ* from 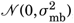 and shifting the peak by *δm*. The standard deviation *σ*_md_ was chosen so that the 3*σ*_md_ interval represents a 10 ppm shift for medium noise and a 20 ppm shift for high noise.
– We simulated intensity variations in the spectrum: Each peak intensity was multiplied by an individual random number *£* drawn from 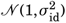. Variance was chosen as 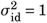 for medium noise and 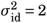 for high noise. Furthermore, 0.03 times the maximum peak intensity of the spectrum was subtracted from each peak intensity. If a peak intensity fell below the threshold of one thousands of the maximum intensity in the spectrum, the peak was *discarded*.
– Finally, we added “noise peaks” to the spectrum. As uniformly choosing the mass of a noise peak would result in peaks which are too easy to spot and sort out by our subsequent analysis^12^, we instead used peaks that appeared in other measured spectra. In preprocessing, a pool of “noise peaks” was gathered from the fragmentation spectra, using all peaks that did not have a molecular subformula decomposition of the known molecular formula of the precursor. For each spectrum, *αn* of these “noise peaks” were added to the spectrum, where *n* is the number of peaks in the spectrum and *α* = 0.2 for medium noise, *α* = 0.4 for high noise. Intensities of “noise peaks” were adjusted for maximum peak intensities in the contributing and receiving spectrum.

Parameters for medium noise and high noise were chosen in a way that the similarity between the original spectrum and the distorted spectrum reached a particular level, measured by the cosine score (dot product): For the cosine score, we allowed a mass deviation of 7 ppm when matching peaks. Precursor ion peaks were not considered for cosine score calculation, as their high intensities overshadow the lower intensity peaks. For medium noise, the cosine score between the original and the distorted spectrum had median value 0.880. For high noise, the median cosine score was 0.714. Datasets with different noise level were used for evaluations only, but not to train CSI:FingerID or individual confidence score SVMs.

Adding noise to the fragmentation spectra may result in an empty or almost empty spectrum, which would be regarded insufficient for structure annotation in applications. To this end, we removed fragmentation spectra with at most one peak. To ensure that evaluation results are comparable between collision energies and noise levels, we discarded the compound from all libraries if a fragmentation spectrum with at most one peak resulted for at least one collision energy and noise level. Doing so, 3,314 compounds were removed from the COSMIC training dataset and 171 compounds from the independent dataset. Substantially more compounds were removed from the COSMIC training dataset because many training dataset spectra have only few peaks, increasing chances that noisy spectra contain at most one peak. Here, 10 eV noisy spectra contain at most one peak for 75 % of the 3,314 removed compounds; 20 eV noisy spectra for 27 %; and 40 eV noisy spectra for 11 % (a compound can exhibit sparse spectra for more than one collision energy).

This resulted in eight libraries, four libraries with 4,046 compounds each for the *COSMIC training dataset*, and four libraries with 3,291 compounds each for the *independent dataset*. Notably, the COSMIC training dataset is a proper subset of the CSI training dataset.

### Biological datasets and data processing

– For the *mice fecal dataset*, we analyzed LC-MS/MS data of 278 samples from a public metabolomics dataset (MassIVE data repository, id no. MSV000082973). This dataset comes from a previously published study^66^; LC-MS/MS experiments were conducted on a Q Exactive Orbitrap instrument (Thermo Fisher Scientific, Bremen, Germany). In brief, the fecal mice metabolome was analyzed by untargeted metabolomics from fecal pellets aqueous-methanol (1:1) extracts from specimens of a atherosclerosis mice model (*Mus musculus atherosclerosis*-ApoE^-/-^). Specimens were either exposed or not exposed to intermittent hypoxia or hypercapnia (IHH). In addition, two groups were fed with a high-fat diet or a regular diet; each group consists of 28 specimens.
– For the *human dataset*, we analyzed ten MassIVE datasets from the MassIVE data repository (id nos. MSV000083559, MSV000079651, MSV000080167, MSV000080469, MSV000080533, MSV000080627, MSV000081351, MSV000082261, MSV000082629, MSV000082630). The dataset contains fecal, plasma, urine, lips, tongue and teeth samples from humans; all acquired on Q Exactive Orbitrap instruments (Thermo Fisher Scientific, Bremen, Germany) in positive ion mode. Runs were acquired using C_18_ RP Ultra High Performance Liquid Chromatography (UHPLC). Only files with extensions “.mzML” or “.mzXML” were considered, and LC-MS runs containing spectra in profiled mode were discarded. This resulted in 2,666 LC-MS/MS runs being processed.
– For the *Orbitrap* dataset, we followed the idea of “flipping the workflow” and reanalyzing public data at a repository scale: We restricted ourselves to MassIVE datasets measured on a Q Exactive Orbitrap instrument (Thermo Fisher Scientific, Bremen, Germany), as this instrument had the largest number of MassIVE datasets. We applied no other constraints with regards to analyzed organism, LC setup etc, resulting in 264 public MassIVE datasets (downloaded Feb 20, 2020). MassIVE datasets containing only spectra in profiled or negative ion mode were discarded, leaving us with 123 MassIVE datasets. Sample types range from environmental to natural products and include biological samples from at least 30 different species, covering diverse genera and phyla. Only files with extensions “.mzML” or “.mzXML” were considered, and LC-MS/MS runs containing spectra in profiled or negative ion mode were discarded, leading to 17,414 LC-MS/MS runs being processed. See Supplementary Table 5 for a list of all MassIVE datasets.

SIRIUS 4 was used to process LC-MS/MS runs and MassIVE datasets provided in mzML or mzXML format. Feature detection in SIRIUS 4 is similar in spirit to a targeted analysis: Instead of searching for all features in a run, SIRIUS first collects all fragmentation spectra and their precursor information, then searches for features that are associated with those fragmentation spectra (precursor ions, adduct ions, isotope peaks). Adducts and isotopes were detected using predefined lists of mass differences. Fragmentation spectra assigned to the same feature (precursor ion) are merged using an agglomerative clustering algorithm based on cosine distance. Compounds with mass beyond 700 Da were discarded to avoid high running time. MassIVE datasets that exceeded 600 LC-MS/MS runs were split to reduce memory consumption.

We use both isotope patterns and fragmentation patterns to determine the molecular formula *de novo* using SIRIUS 4 with default parameters and mass accuracy 10 ppm. CSI:FingerID with default parameters was used to rank structure candidates. We use SIRIUS default soft thresholding of molecular formulas when querying CSI:FingerID structure candidates; CSI:FingerID result lists can therefore contain structures with different molecular formulas. We used the highest-scoring structure candidate and the corresponding fragmentation tree, isotope pattern and structure candidate list features for COSMIC.

For the mice fecal dataset, SIRIUS results were imported into GNPS, and data were further annotated and explored by performing feature-based molecular networking and spectral library search on GNPS. The statistical and fold change analysis was performed using MetaboAnalyst 4.0^67^ for samples from control mice (not exposed to IHH) that were either fed on a high-fat diet or regular diet.

### Receiver operating characteristics and hop plots

We are given a list of hits, one for each query, sorted by score. Each hit can either be *positive* (correct annotation) or *negative* (incorrect annotation). Varying a score threshold, we can modify the number of hits reported to the user; our goal is to report all positives and to reject all negatives. True positives (*TP*) and false negatives (*FN*) are positives (correct hits) which pass or do not pass the threshold; similarly, false positives (*FP*) and true negatives (*TN*) are incorrect hits which pass or do not pass the threshold. For any score threshold, we plot the *true positive rate TP*/(*TP + FN*) (ratio of reported correct hits among all correct hits) against the *false positive rate FP*/(*FP + TN*) (ratio of reported incorrect hits among all incorrect hits), resulting in a Receiver Operating Characteristic (ROC) plot. The Area Under Curve (AUC) of the ROC curve is the integral of the ROC curve; the random score, corresponding to a random ordering of hits, reaches AUC 0.5. A method may reach AUC below 0.5, meaning that the hit score performs worse than random. Different from binary classification, we must not invert “predictions” to reach a better AUC: Logic dictates that the directionality of the hit score (such as, “high scores are good”) is fixed by the candidate identification task. Unfortunately, the AUC measure makes no difference between the (highly relevant) lower-left and the (mostly irrelevant) upper-right of the ROC curve.

In contrast to binary classification, two methods can differ in the number of positives (correct hits, correct annotations) they reach for the complete list of queries. This is a peculiarity of the identification task and has no equivalent in binary classifier evaluation, where the number of positives and negatives is determined by the dataset. ROC curves do not asses the number of positives; in particular, two methods can have identical ROC curves although one method reaches twice as many correct hits. We introduce *hop plots* (inspired by the hop plant *Humulus lupulus* ranking to a supporting wire) to integrate this information: We again vary the score threshold but normalize reported correct hits and incorrect hits by the *total* number of hits (queries) *N = TP + FN + TN + FP*, plotting *TP*/*N* vs. *FP*/*N*. See Supplementary Fig. 2. The resulting curve starts in the origin (0, 0) and ends in some point (*x*, *y*) *∈* [0, 1]^2^ with *x* + *y* = 1, where *y* is the ratio of correct hits for the complete list of queries. The hop curve lies in the lower-left triangle; random ordering of hits corresponds to a straight line from the origin to some point (*x*, *y*) with *x* + *y* = 1. For perfect results, the hop curve is a straight line between the origin and (0, 1); in the worst case, it is a straight line from the origin to (1, 0). Hop plots allow us to answer questions such as, “If I fix some false discovery rate, how many true discoveries will a method return?” We stress that to draw a ROC curve or a hop plot, we must have complete information about true and false positives and negatives, so we can calculate the *exact* FDR as *FP*/(*FP + TP*). A zoom-in allows us to compare methods in the particularly interesting region close to the origin. Both ROC curves and hop plots allow us to visually compare the performance of a method for different datasets in one plot; here, the total number of hits *N* is different for each curve.

We can calculate the *area under curve* of a hop plot by mirroring the curve at the line *x* + *y* = 1 before taking the integral. A method with identification rate *y* ∈ [0, 1] for the complete list of queries will have area under curve between *y*^2^ and *y*^2^ + 2(1 − *y*) *y* = 1 − (1 − *y*)^2^, with random ordering reaching area *y*^2^ + (1 − *y*) *y* = *y*. But much like the area-under-curve of a ROC curve, this number does not tell us whether a method performs well at the (highly relevant) lower-left or the (mostly irrelevant) upper-right of the curve; hence, we refrain from reporting hop plot area-under-curve.

### Training CSI:FingerID and structure-disjoint evaluation

We trained an array of Support Vector Machines (SVM) for fingerprint prediction from MS/MS data as described in [15, 30, 53]. Training of CSI:FingerID was carried out using merged spectra with all available collision energies from the training dataset. In contrast, single collision energy and merged spectra libraries as well as noisified spectra were not used when training CSI:FingerID, but only in validation of COSMIC. We used PubChem-standardized structures^68^ when computing the molecular fingerprint of a compound. In evaluations, we used the CSI:FingerID “covariance score” from [69] to rank candidates, comparing the probabilistic query fingerprint and each structure candidate fingerprint. A hit was regarded as *correct* if the PubChem-standardized structures of query and top rank were identical.

As noted above, all evaluations were carried out structure-disjoint. For the tenfold cross-validation, we partitioned the training data into ten disjoint *batches* of almost identical size, ensuring that all fragmentation spectra from compounds with identical structure (such as L-threose and D-erythrose) *end up in the same batch*: Otherwise, L-threose could be part of the training data when evaluating on D-erythrose, and vice versa. For each batch, we trained the fingerprint SVM array using the remaining nine batches; we evaluated on the tenth batch. In this way, we ensured that all compounds are novel for CSI:FingerID: For each query, MS/MS training data for the corresponding structure, including independent MS/MS measurements, were not available for CSI:FingerID. CSI:FingerID evaluations on the independent dataset were again executed structure-disjoint: We additionally trained an SVM array using the complete training dataset. Given an MS/MS query from the independent data, we checked if the structure of the query is also part of the training data: If so, we used the appropriate SVM array from cross-validation for fingerprint prediction; otherwise, we used the SVM array trained on the complete training data. Again, this ensured all structures being novel in evaluation.

### Score calibration and E-value estimation

The *p-value* of a score is the probability that a score this high or higher would be expected by chance; the *E-value* is the expected number of random hits with this score or higher. Kim *et al.*^70^ suggested to use E-values for peptide database searching; MS-GF E-value computation uses dynamic programming, based on the linear nature of peptides. Keich *et al.*^71^ calibrate peptide database search scores using decoys. Both approaches are conceptually hard to adopt for metabolite annotation: Metabolites have highly non-linear structure, and no methods have been suggested to generate reasonable decoy molecular structures for small molecules^12^.

We suggest to use the distribution of scores of PubChem^31^ candidates as a proxy for the score distribution of incorrect hits. We empirically established that scores of an individual MS/MS query roughly followed a log-normal distribution; for other queries, the score distribution was multimodal (Supplementary Fig. 4). In particular, a small fraction of candidates had a much higher score than expected from the single log-normal distribution; ignoring this would result in inflated calibrated scores.

The log-normal distribution is a reasonable proxy if there are only few samples available. To model multimodal distributions as well as distributions that deviate from the log-normal distribution, we suggest to use a kernel density estimate of the probability density function. Clearly, we do not have to “compute” the kernel density; instead, we want to know the E-value under the resulting distribution. For the ease of presentation, we do not use log-normal kernel functions but instead, model the log-transform of the scores by normal kernel functions, which is mathematically equivalent. Let *y_i_* := ln*x_i_* for *i* = 1,…, *n* be the log-scores of the PubChem “proxy decoys” *excluding the hit score*, and let *y* := ln *x* be the log-score of the hit. We first determine the bandwidth of the kernel function; we use Silverman’s rule of thumb, first determining the standard deviation *σ*^ of the sample *y*_1_,…, *y_n_*, then setting

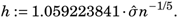

We also tested other bandwidth estimation procedures but did not find a substantial difference (data not shown). For the Gaussian kernel 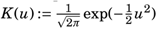 we reach

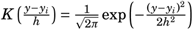

so this is just the usual probability density function of the normal distribution times *h*, which cancels out in the kernel estimator. We calculate

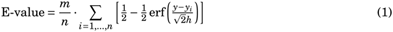

where *m* is the number of candidates in the biomolecule structure database.

### Confidence score estimation

Our method of confidence estimation is inspired by the Percolator method for peptide identification in shotgun proteomics^72, 73^. Different from there but similar to [74, 75], we do not train a classifier for an individual LC-MS run to “boost” annotations; instead, we train classifiers only once using the reference measurements, which are then applied to the biological data. As noted by Käll *et al.*^72^, this approach is highly prone to overfitting: Characteristics of correct and incorrect hits may vary between experiments, instrument types, compounds present in the sample, and others. Here, we have taken extensive measures to counter overfitting, such as “noisifying” spectra and the restriction to *linear* Support Vector Machines.

We repeated the following for each collision energy (10 eV, 20 eV, 40 eV, and merged spectra), and trained individual SVMs using spectra without added noise from that energy as training data. Features of the linear SVMs are shown in Supplementary Table 1. All features were individually standardized. Parameter *C* ∈ {10^−5^, 10^−4^,…, 10^5^} of each SVM was chosen by a nested cross-validation. We used quadratic hinge loss and *l*_2_ regularization. SVMs were trained using LIBSVM^76^.

For each collision energy, we trained three classifiers: (i) When searching PubChem, we used all appropriate features (all but Features 20–22) from Supplementary Table 1. Searching the biomolecule structure database, not all queries result in two or more candidates; but some features from Supplementary Table 1 require a candidate list of at least size two, such as the difference between score of highest-scoring vs. runner-up candidate. To this end, we trained two classifiers for the biomolecule structure database: (ii) The regular SVM assumes that there are at least two candidates; it uses all features from Supplementary Table 1 but is trained only on the appropriate subset of the training data. (iii) The single-candidate SVM uses only the appropriate sub-features (all but Features 1–4, 10, 13) but can be trained using all training data: For instances with two or more candidates, we uniformly selected one candidate.

Unfortunately, the resulting linear classifiers showed clear signs of overfitting: For example, some features received weights which were counterintuitive, such as negative weight for the quality of the SIRIUS fragmentation tree or the CSI:FingerID score. Recall that the actual hit was chosen by CSI:FingerID as the candidate with the *highest* score; hence, logic dictates that the CSI:FingerID score of the hit *must not* receive a negative weight when deciding whether a hit is correct or incorrect. The same is true for selecting the best fragmentation tree by SIRIUS. To this end, we *enforced directionality* of the features: For each feature, we decided manually whether a high value of the feature would increase or decrease our confidence in an annotation. For example, a high CSI:FingerID score should clearly increase our confidence, and so should a small E-value. See Supplementary Table 1 for enforced directions. Notably, enforcing directionality can be achieved by a regular SVM optimization without additional constraints, allowing us to use established SVM solvers: For each feature with enforced directionality, we augmented one training sample where the corresponding feature was set to a large (positive or negative) value *±β*, whereas all other features were kept at zero; the sample received a positive label (correct hit). If the absolute feature value *β* > 0 is large enough, then an optimal solution must use the feature in the desired direction; the actual value *β* is of minor importance due to the hinge loss of SVM optimization. To avoid potential numerical instabilities when finding the solution, *β* should not be chosen too large. Here, we used *β* = 10^7^; using absolute feature values 10^8^ and 10^9^ resulted in basically identical models, and differences are of no practical consequence (data not shown). Notably, some features received non-zero weights for the classifier with enforced directionality, despite the fact that these features received “counter-intuitive” weights in the unrestricted optimization: For example, feature “FP Length Hit” was repeatedly given negative weight in cross-validation but had high positive weight if we enforced directionality (unrestricted weight −0.00165, restricted weight 0.0568 in the same cross-validation fold).

When training the COSMIC SVMs, all CSI:FingerID fingerprint predictions of training spectra were carried out structure-disjoint using CSI:FingerID cross-validation models. The COSMIC training data was then partitioned for tenfold cross-validation in the same fashion as for CSI:FingerID training. Hence, cross-validation evaluation of COSMIC is again structure-disjoint, and all compounds are novel. Similar to above, we also ensured structure-disjoint evaluations on the independent dataset, by choosing the appropriate SVM from cross-validation for computing the confidence score. When applying the model to independent data, we capped feature values: For each feature from Supplementary Table 1, we record the minimum and maximum feature value in our training data. When applying the model, feature values exceeding these thresholds are set to the respective threshold value. We do so to prevent exaggerated decision values caused by unexpectedly high values of one or more features.

We map decision values to posterior probability estimates using Platt probabilities^32^. Platt^32^ proposed to use a sigmoid function as an approximation of posterior probabilities: 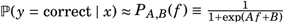, where *f* = *f* (*x*) ∈ ℝ is the decision value for hit *x*, and *y* ∈ {correct, incorrect} its label. We estimated parameters *A*, *B* ∈ ℝ using maximum likelihood^32, 77^ as implemented in LIBSVM.

Using a linear classifier enables explainable machine learning: See Supplementary Table 1 for feature weights after normalization of the three classifiers for merged spectra. We observe that certain features have weight close to zero; this might indicate that the feature is indeed uninformative, that the feature does not measure what we intended to measure, or that our training data is insufficient to learn a reasonable weight.

Recall that confidence SVMs were trained exclusively on spectra without added noise; we also trained SVMs from a combined dataset with all noise levels, but found that results were of identical quality (data not shown).

Unlike Percolator^72, 73^, we do not learn a confidence score for individual LC-MS datasets. We do so because it is non-trivial to generate reasonable decoys for small molecules and, more importantly, since incorrect hits in the target database are often not random (Fig. 4)^34^. This potentially explains why the calibrated E-value score presented here, does not allow for a satisfactory separation. Also unlike Percolator, we do not use our scores to rerank candidates^72, 73^: All of our candidates share the same molecular formula, fragmentation tree and predicted fingerprint; these features are meaningless for reranking. To this end, curves of CSI:FingerID and COSMIC in hop plots (Fig. 2, Fig. 5) always end in the same point (*x*, *y*) with *x* + *y* = 1.

### False discovery rate estimation

Recall that the false discovery rate equals *FP*/(*FP+TP*) where *TP* is the number of true positives (correct hits above some score threshold) and *FP* is the number of false positives (incorrect hits above the same score threshold). Also recall that to compute FDR, we must know the exact numbers *FP* and *TP*. Yet, in applications, we do not have this information; in this case, we need some method to *estimate* FDR values. Returning random numbers would be an admissible method for FDR estimation, albeit a useless one; to this end, a method for FDR estimation has to be validated against exact FDR values, to assess its accuracy. In application, a user selects an acceptable FDR level, and we want to return as many hits as possible so that the list of hits meets the preselected FDR. The *q-value* of a hit is the smallest FDR at which this hit is part of the output list.

We now show how to transform COSMIC confidence scores to FDR estimates. The confidence score is an estimated posterior probability of the hit to be correct; to this end, it is one minus the posterior error probability for this hit. Hence, we can use the confidence score to estimate the FDR of the top *k* hits^12, 35^: Let *p_j_* be the posterior error probability for hit *j* for *j* = 1,…, *n*, and assume that the hits are ordered by confidence score, so *p_j_* ≤ *p*_*j*+1_. Viewing the annotations as (not necessarily independent) Bernoulli trials, the expected number of incorrect annotations for the top *k* hits is 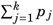, and the expected false discovery rate is

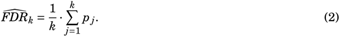

Since hits have been ordered by posterior error probability, FDR estimates 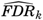 are monotonically increasing, so 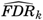 is also the q-value estimate for hit *k*.

We evaluate the accuracy of our FDR estimates by plotting exact q-values against estimated q-values in a Q-Q-plot (Supplementary Fig. 9); this has to be carried out using reference data where exact FDR values can be calculated.

### Comparing molecular structures

The *Tanimoto coefficient* measures the similarity of two molecular structures. Any Tanimoto coefficient is based on a particular set of molecular properties, constituting the *fingerprint type*. For consistency, we use the same fingerprint type (molecular properties) throughout this manuscript that we have trained SVMs for as part of CSI:FingerID. The Tanimoto coefficient is the Jaccard index of the two sets of molecular properties, that is, the cardinality of the intersection of the two sets divided by the cardinality of the union of the two sets. The advantage of the Tanimoto coefficient is that it can be quickly calculated, in particular if we have precomputed the fingerprints of all molecular structures of interest.

For highly similar molecular structures such as the pairs in Fig. 4, it is not advisable to employ the Tanimoto coefficient, as it is not apt to accurately measure such high similarity. Instead, we represent the two molecular structures as graphs, and ask for a minimum number of edges that have to be removed from the graphs such that the resulting graphs are isomorphic; naturally, hydrogen atoms are ignored in this computation. This is the Maximum Common Edge Subgraph (MCES) problem; using the number of removed edges to estimate dissimilarity is an appropriate measure for highly similar molecules, as we explicitly do not demand that the resulting subgraph is connected. Unfortunately, the MCES problem is NP-complete, as it generalizes subgraph isomorphism. See e.g. [78] for a discussion of available methods for solving MCES exactly and heuristically.

For the molecular structures from Fig. 4, it is straightforward to manually find optimal solutions: The “top hit” structure can be transformed into the “correct hit” structure via two edge deletions for examples (a–c) and (f–i), whereas examples (d) and (e) require four edge deletions. Since both graphs have the same number of edges, we require at least two edge deletions for non-isomorphic graphs.

### CASMI 2016 re-evaluation

Scores of MetFrag, MAGMa+, CFM-ID, CSI:FingerID (original) and CSI:FingerID IOKR (Input Output Kernel Regression) were downloaded from the CASMI 2016 results web page (http://casmi-contest.org/2016/, category 2, automated methods). We only consider tools that scored all candidates. CSI:FingerID (original) and CSI:FingerID IOKR were not executed structure-disjoint, as CASMI is a blind competition. We computed scores for the structure-disjoint evaluation of CSI:FingerID using the current version of CSI:FingerID.

We used hit scores (score of the top-scoring candidate for each query) to rank hits. For consistency, we restricted the set of candidate structures to those with the correct molecular formula for all tools. We performed evaluation using either all ChemSpider candidates, or restricting the search to those ChemSpider candidates that are simultaneously found in our biomolecule structure database. In 3 cases, this resulted in an empty list of candidates, and these queries were excluded from evaluation. In 13 cases, the set of candidates did no longer contain the correct structure; these queries were not excluded from evaluation. As expected^79^, MetFrag, MAGMa+ and CFM-ID profit more from restricting the set of candidates than CSI:FingerID^15^; hence, annotation rates varied less than those reported in the CASMI evaluation^18^. In fact, even randomly choosing one of the remaining candidates resulted in a decent annotation rate when searching the biomolecule structure database: In 38 cases, only a single candidate remained; and in 33 cases, the candidate list contained two or three structures. Even if there is only a single candidate, the score some *in silico* tool assigns to this candidate is important information, as we use it to rank hits.

The fact that scores of *in silico* tools, including CSI:FingerID, cannot be used to decently separate correct and incorrect hits, may be unexpected for users; but tools and scores were not developed with this application in mind. To this end, *our findings must not be misunderstood as a critique* against these tools or their developers.

COSMIC confidence scores were computed as described above, using the confidence score model for “all fragmentation energies”. We ensured structure-disjoint evaluation (all compounds novel) both for CSI:FingerID and COSMIC, as detailed above. Both for ChemSpider and the biomolecule structure database, we used the confidence score variant for searching the biomolecule structure database; this is reasonable as the number of ChemSpider candidates is often substantially smaller than the number of PubChem candidates.

For completeness, we also evaluated separation of the original submissions of CSI:FingerID and CSI:FingerID IOKR (Supplementary Fig. 3). As noted, these evaluations were not carried out structure-disjoint; hence, results mix dereplication (structures for which MS/MS data is available in the training data) and novel structure search. Unfortunately, we cannot compute confidence scores for the original CSI:FingerID submission, as features required for its computation (Supplementary Table 1) were not recorded when submitting the CASMI entry.

### In-depth method evaluation

For a query fragmentation spectrum, we again assume to know its molecular formula, and we obtained candidates from the structure databases using this molecular formula. In practice, molecular formulas can be established using SIRIUS 4^30^ or ZODIAC^80^. For 325 compounds in the training data and 278 compounds in the independent data, this resulted in an empty candidate list when querying the biomolecule structure database; these compounds were excluded from evaluation, leaving us with 3,721 queries in cross-validation and 3,013 queries for independent data. For 845 compounds in the training data and 521 compounds in the independent data, the correct structure is not present in the biomolecule structure database; *these compounds were not excluded*. We ensured structure-disjoint evaluation (all compounds novel) both for CSI:FingerID and COSMIC.

To evaluate against spectral library search, we generated two spectral libraries based on the CSI training dataset: One library with merged spectra, and one library with spectra at individual collision energies as well as merged spectra. We searched merged query spectra in the first library, and query spectra containing a single collision energy in the second library. Merged spectra are identical to those used for training CSI:FingerID, see above; this library contains 23,965 spectra. The second library contains all available fragmentation spectra at all available collision energies, plus the merged spectra, and contains 189,979 spectra. This resembles how spectral library search is usually applied in practice. To ensure a fair comparison with COSMIC, spectral library search candidates were restricted to those with the correct molecular formula for each query; in practice, this information is usually not available, and spectral library search may perform worse than reported here. In case the spectral library did not contain at least one candidate with the correct molecular formula of the query, a misannotation with score zero was assumed. We evaluated both the cosine score described above, as well as a cosine score using the square root of intensities.

The term “spectral library search” refers to searching for a query fragmentation spectrum in a database of reference fragmentation spectra measured from (usually commercial) standards, then reporting the highest-scoring candidate (hit) under some scoring. Spectral library search must not be mistaken with the task of comparing mass spectra, manually or automated, or with computing a measure of similarity between spectra such as the cosine score. Comparison of mass spectra is in use for many research questions beyond spectral library search: This includes the manual validation of annotations, MASST^28^, or CSI:FingerID (and, hence, COSMIC) that uses the cosine score as part of its machine learning framework.

### Annotation and validation of novel bile acid conjugates

For the mice fecal dataset, MS/MS measurements were taken with a collision energy of 30 eV; we used the COSMIC version trained on 40 eV spectra. The bile acid conjugates structure database was used for the annotation. No additional parameters have to be chosen in the COSMIC workflow.

The output of this workflow is a ranked list of 1,456 COSMIC structure annotations (Supplementary Material 3). In case multiple compounds were annotated with the same structure (for example, compounds being present in multiple runs, and different adducts of the same compound), entries in the COSMIC output were merged and represented by the hit with the highest confidence. This reduces the output to 626 unique structure annotations (Supplementary Table 2). Of these, 113 were present in PubChem. Here, we concentrated on the 513 “truly novel” bile acid conjugates.

The top 12 most confident bile acid conjugate annotations were manually inspected and the fragmentation interpreted to check consistency with structure proposed by COSMIC (Supplementary Fig. 12 to 23). The fragmentation of bile acid conjugates is characterized by fragment ions and neutral losses from the conjugated amino acid moiety as well as the hydroxylation pattern of the bile acid core. Annotations of two “truly novel” bile acid conjugates, phenylalanine (Phe) and tryptophan (Trp) conjugates of chenodeoxycholic acid (CDCA), were verified by comparing their fragmentation spectra and retention times with those of synthetic standards. Phe-CDCA (**7**) and Trp-CDCA (**12**) were synthesized using a procedure adapted from a previous method by Ezawa *et al.*^81^. Chenodeoxycholic acid (98.1 mg, 0.25 mmol, 1 eq.) was dissolved in THF (4.9 mL, 0.05 M and cooled to 0 °C with stirring. Ethyl chloroformate (28 μL, 1.2 eq.) was added followed by triethylamine (41 μL, 1.2 eq); then, the reaction was stirred for 2 h in an ice bath. After complete conversion of the starting material by TLC a cold, aqueous solution (4.9 mL) of amino acid (0.37 mmol, 1.5 eq.) and NaOH (14.8 mg, 0.37 mmol, 1.5 eq.) was added in one portion. The reaction was then stirred for 2 h gradually warming to room temperature. THF was removed *in vacuo* and 2 M HCl was added to acidify to pH < 2, at which point a white precipitate appears. The mixture was extracted with ethyl acetate (3 × 20 mL), and the combined organic layers were washed with brine (1 × 50 mL), dried over sodium sulfate, and concentrated. The crude material was purified over silica gel by column chromatography eluting with 3-10 % methanol in dichloromethane (plus 1 % acetic acid, v/v) to yield the desired products as confirmed by nuclear magnetic resonance (NMR) spectroscopy. NMR spectra were recorded on a BRUKER Avance (600 MHz, CryoProbe) spectrometer in CD_3_OD. Signals are reported in ppm with the internal CD_3_OD signal at ppm (^1^H) and 49.0 ppm (^13^C) as standard reference peak.

– **Phe-conjugated chenodeoxycholic acid (Phe-CDCA)**: Product was obtained as a white solid in 91 % yield. ^1^H NMR (599 MHz, MeOD): 7.29–7.18 (m, 5H), 4.67–4.61 (m, 1H), 3.81–3.78 (m, 1H), 3.42–3.33 (m, 1H), 3.22 (dd, *J* = 14.4, 4.8 Hz, 1H), 2.93 (dd, *J* = 13.8, 9.0 Hz, 1H), 2.27 (q, *J* = 12.0 Hz, 1H), 2.22–2.17 (m, 1H), 2.10–2.03 (m, 1H), 2.01–1.94 (m, 2H), 1.90–1.81 (m, 3H), 1.77–1.57 (m, 4H), 1.54–1.45 (m, 4H), 1.41–1.26 (m, 5H), 1.24–1.04 (m, 5H), 1.03–0.95 (m, 1H), 0.93–0.86 (m, 7H). ^13^C (151 MHz, MeOD): 175.22, 137.28, 128.88, 128.00, 126.32, 71.47, 67.66, 55.94, 50.13, 42.26, 41.78, 39.65, 39.37, 39.08, 37.06, 35.44, 35.17, 34.83, 34.51, 32.65, 32.45, 31.82, 29.96, 27.84, 23.23, 22.02, 20.39, 17.48, 10.81.
– **Trp-conjugated chenodeoxycholic acid (Trp-CDCA)**: Product was obtained as a white solid in 42 % yield. ^1^H NMR (599 MHz, MeOD): 7.56 (d, *J* = 7.8 Hz, 1H), 7.33 (d, *J* = 7.8 Hz, 1H), 7.10–7.06 (m, 2H), 7.00 (t, *J* = 7.8 Hz, 1H), 4.73 (dd, *J* = 8.4, 4.8 Hz, 1H), 3.81–3.77 (m,1H), 3.41–3.32 (m, 2H), 3.18–3.13 (m, 1H), 2.31–2.16 (m, 2H), 2.10–2.03 (m, 1H), 1.98–1.93 (m, 2H), 1.88–1.78 (m,3H), 1.73–1.63 (m, 3H), 1.63–1.58 (m, 1H), 1.54–1.43 (m, 5H), 1.41–1.25 (m, 5H), 1.24–0.94 (m, 7H), 0.91(s, 3H), 0.89 (d, *J* = 7.2 Hz, 3H), 0.63 (s, 3H). ^13^C (151 MHz, MeOD): 176.65, 175.39, 138.01, 128.87, 124.24, 122.37, 119.78, 112.28, 111.12, 72.84, 69.07, 57.22, 54.61, 51.49, 43.62, 43.14, 41.00, 40.72, 40.45, 36.78, 36.53, 36.20, 35.89, 34.02, 33.78, 32.99, 31.33, 29.16, 28.45, 24.60, 23.39, 21.76, 18.86, 12.15.

Reference standards were analyzed by LC-MS/MS using identical experimental conditions as used previously^82^. Retention times were 298.5 seconds for Phe-CDCA and 294.5 seconds for Trp-CDCA. Samples from the previous study were renanalyzed to ensure comparability of retention times: The putative Phe-CDCA and Trp-CDCA candidates had retention times of 300.5 and 294.5 seconds, respectively. Considering the similarity of their fragmentation spectra (Supplementary Fig. 24) and retention times, these are MSI level 1 identifications. Yet, these identifications are not unambiguous: Isomeric structures such as Phe-deoxycholic acid would show the same fragmentation spectrum and the same or very similar retention time. For a conclusive decision, a more detailed analysis method would be required, which is out of the scope of this paper.

Molecular networks were visualized in Cytoscape (version 3.7.1)^83^. The MetaboAnalyst web server^67^ was used to process the feature quantification results and perform statistical analysis. Quantile normalization and auto-scaling were used. Results of the fold change analysis were mapped onto molecular networks using Cytoscape. MASST^28^ was used to search the annotated bile acid conjugates spectra in all public mass spectrometry datasets including MassIVE-GNPS^37^, MetaboLights^4^ and MetabolomicsWorkbench^5^. Parameters and results for these jobs are part of Supplementary Table 2.

### Repository-scale annotation of novel metabolites

To estimate a reasonable COSMIC confidence score cutoff, we made use of our reference data evaluation results. In our evaluation using independent data, collision energy 20 eV and medium noise, a confidence score threshold of 0.64 corresponded to FDR 10 %. Our implicit assumption is that for the biological data, this threshold will correspond to a similar FDR. It must be understood that we cannot guarantee a similar FDR for structure annotations below, given our inability to accurately estimate FDR. Clearly, numerous hits with confidence below this threshold will nevertheless be correct.

We searched the human dataset against the biomolecule structure database; this resulted in 114,012 hits. Multiple hits can annotate the same structure; for example, these hits may originate from different LC-MS/MS runs or different adducts. Hence, we report unique structures instead, where the hit with the highest confidence is used as a representative for that structure. This resulted in 24,554 unique structures being annotated, of which 3,167 (12.9 %) were present in the CSI training dataset. We now filter the 24,554 structure annotations for high confidence (score threshold 0.64), resulting in 911 structure annotations. Of these high-confidence annotations, 475 (52.1 %) were present in the SVM training data, leaving us with 436 (47.9 %) high-confidence novel structure annotations. Finally, we excluded all hits with structures in the HMDB structure database, resulting in 21,128 unique structure annotations, 436 high-confidence structure annotations, and 315 high-confidence structure annotations without reference MS/MS data (Fig. 27). Of the 315 novel structures, 48 were proteinogenic peptides.

We searched 14 character InChI keys of all 261 novel metabolite structures in the current version of HMDB (Feb 2021) and found that at least 23 of these structures are present in the current HMDB version. The exact number may be slightly higher, as structures from the current HMDB version were not standardized using the PubChem standardization procedure. Notably, the recent inclusion of structures in HMDB does not mean that reference MS/MS data are available for these structures.

High-confidence hits were manually evaluated by checking spectra for known neutral losses and fragments that can be explained. Furthermore, spectra were compared against reference spectra from similar structures. The following paragraphs exemplary discuss high-confidence annotations where evaluation based on manual interpretation or newly generated reference spectra was possible. For none of the structures verified by spectral comparison ((2E)-octenoyl-carnitine, N-oleyl-leucine, phenazine-1,6-dicarboxylic acid), reference spectra were available in the training data of COSMIC or CSI:FingerID.

Firstly, acyl-carnitine structures were evaluated by their typical fragmentation. Characteristics fragments are found at *m/z* 85 and *m/z* 144. These are derived from an ene-type loss of the neutral fatty acid yielding *m/z* 144, undergoing a further loss of trimethylamine to yield *m/z* 85. The same loss of trimethylamine can occur from the intact molecule, yielding a fragment found at a neutral loss of 59 Da. Based on this fragmentation pattern eight high-confidence hits were ruled out and are presumably incorrect annotations. Out of these eight bogus annotations, three potentially have the wrong adduct annotation. Based on our manual validation, 21 out of 30 annotations of the acyl-carnitines are correct. Furthermore, the query spectrum annotated as (2E)-octenoyl-carnitine showed good agreement with a reference measurement (Supplementary Fig. 28).

Second, several N-acyl-amino acids were manually validated. Fragmentation of [M + H]^+^ adducts of N-acyl amino acids typically yields an ene-type of loss of a neutral fatty amide or the neutral loss of a fatty acyl ketene structure. Further fragmentation yields typical amino acid fragments allowing to potentially identify the amino acid in more detail. Within the human dataset, N-oleyl-leucine was annotated with a high-confidence score. For this structure, reference spectra are now available in MassBank^65^. A high spectral similarity (cosine score 0.85) was found between the spectrum and the reference (Supplementary Fig. 28). Since MS cannot differentiate between isomeric species, the structure might also represent N-oleyl-isoleucine: Spectra of N-oleyl-leucine and N-oleyl-isoleucine are both present in MassBank, but indistinguishable. Another example is N-palmitoyl-tryptophan: No reference spectrum is available for this substance, but the observed fragmentation pattern is in good agreement with expected fragmentation, showing *m/z* 205 which relates to tryptophan based on the loss of palmitic acid as ketene, and *m/z* 188 related to the loss of palmitic acid as neutral amide. Further fragments are typically observed in the fragmentation of tryptophan. Using MASST, twelve additional human datasets containing a similar spectrum were identified.

Third, phenazine-1,6-dicarboxylic acid was annotated in a human urine dataset. This metabolite is produced by *Streptomyces* and *Pseudomonas* species^84^, hinting at a potential urinary tract infection. Again, the query spectrum showed good agreement with a reference measurement (Supplementary Fig. 28).

Compound classes in Supplementary Fig. 27 were assigned by NPClassifier [85]. Proteinogenic amino acids and peptides were selected manually. We have installed a web interface allowing interested users to browse through structure annotations sorted by confidence, check spectra, access underlying datasets, leave comments and judge the overall quality of the annotation for the human dataset. The web interface is available from https://bio.informatik.uni-jena.de/cosmic.

To demonstrate that COSMIC can be applied at a repository scale, we searched the Orbitrap dataset with 17,414 LC-MS/MS runs against the biomolecule structure database; this resulted in 979,521 hits. Again, multiple hits can annotate the same structure; the above hits correspond to 77,932 unique annotated structures, of which 8,172 (10.5 %) were present in the CSI training dataset. We now filter the 77,932 structure annotations for high confidence (score threshold 0.64), resulting in 3,530 structure annotations. Of these high-confidence structure annotations, 1,815 (51.4 %) were present in the CSI training dataset, leaving 1,715 (48.6 %) high-confidence novel structure annotations (Supplementary Fig. 29). Again, all hits of the Orbitrap dataset can be accessed via a web interface available from https://bio.informatik.uni-jena.de/cosmic.

*Data availability.* Input mzML/mzXML files are available at MassIVE (https://massive.ucsd.edu/) with accession nos. MSV000082973 (mice fecal dataset); MSV000084630 (mass spectrometry analysis of the synthetic standards for Phe-CDCA and Trp-CDCA); MSV000083559, MSV000079651, MSV000080167, MSV000080469, MSV000080533, MSV000080627, MSV000081351, MSV000082261, MSV000082629, MSV000082630 (human dataset). See Supplementary Table 5 for accession numbers of the Orbitrap dataset. Metadata for synthetic standards of Phe-CDCA and Trp-CDCA were deposited with the dataset (id no. MSV000084630). Fragmentation spectra of Phe-CDCA and Trp-CDCA were deposited on GNPS (CCMSLIB00005467952 and CCMSLIB00005716808).

Fragmentation spectra of all other manually validated bile acid conjugates were also deposited on GNPS; see Supplementary Table 2 for individual spectra ids. Fragmentation spectra of N-oleyl-leucine (RP029701 ([M + H]^+^, 10 eV), RP029702 ([M + H]^+^, 20 eV), RP029703([M + H]^+^, 40 eV) and phenazine-1,6-dicarboxylic acid (RP018701 ([M + H]^+^, 10 eV), RP018702 ([M + H]^+^, 20 eV), RP018703 ([M + H]^+^, 40 eV) are available from MassBank. Fragmentation spectrum of (2E)-octenoyl-carnitine was deposited on GNPS.

Parameters and results of LC-MS/MS processing for the mice fecal dataset are available at https://gnps.ucsd.edu/ProteoSAFe/status.jsp?task=e78a8c8f429a46fcb24f3b34d69aff25. All used structure databases are available as Supplementary Material.

*Code availability.* COSMIC is written in Java, and will be integrated into an upcoming version of SIRIUS 4; it is open source under the GNU General Public License (version 3). It is available for Windows, macOS X, and Linux operating systems. We also provide source code, executable binaries, living documentation, training videos, sample data as well as the public part of the training data on the SIRIUS website (https://bio.informatik.uni-jena.de/sirius/); a source copy is hosted on GitHub (https://github.com/boecker-lab/sirius/).

Scripts for generating the bile acid conjugate database are available from https://bio.informatik.uni-jena.de/software/cosmic/.

## Acknowledgments

M.A.H., M.L., M.F., K.D. and S.B. supported by Deutsche Forschungsgemeinschaft (BO 1910/20 and 1910/23). L.F.N. supported by U.S. National Institutes of Health (R01 GM107550). P.C.D. supported by Gordon and Betty Moore Foundation (GBMF7622), and U.S. National Institutes of Health (R01 GM107550, P41 GM103484, R03 CA211211, U19 AG063744 01). We thank F. Kuhlmann and Agilent Technologies, Inc. for providing data used in the evaluation of COSMIC. We thank all the researchers that contributed to the public mass spectrometry data that we have analyzed with COSMIC. We also thank Michael J. Meehan for support in performing the mass spectrometry experiments.

## Author contributions

S.B. designed the research. M.A.H., M.L. and S.B. developed the computational method, with the help of K.D. M.A.H. implemented the computational method, with contributions from M.L., M.F. and K.D. M.F. integrated COSMIC into SIRIUS. L.F.N. and M.A.H. generated, searched, evaluated and validated the bile acid conjugates. E.C.G. prepared the standard bile acid conjugates, and confirmed standards via NMR. M.W. evaluated and validated search results from the human dataset. L.F.N., M.W. and P.C.D. contributed to the biochemical interpretation of results. M.A.H., L.F.N., M.L., M.F., M.W., K.D. and S.B. prepared the figures. M.A.H., L.F.N., M.W. and S.B. wrote the manuscript, in cooperation with all other authors.

## Supplementary Tables

**Supplementary Table 1:**
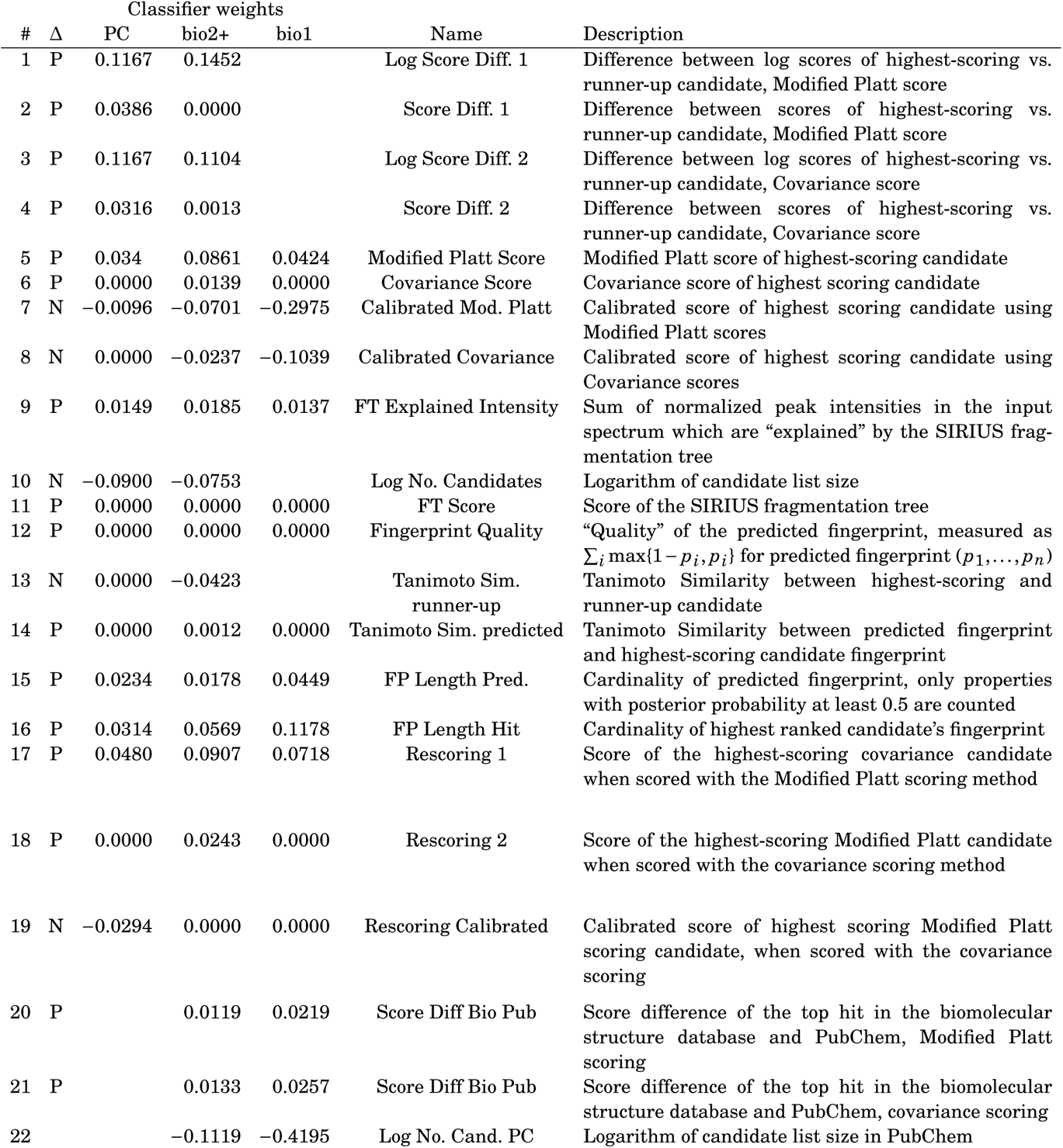
Features of the COSMIC confidence score and classifier weights for merged spectra. Features 1–19 are used for PubChem and biomolecule structure confidence scores, features 20–22 are exclusive to the biomolecule structure confidence scores. For the linear SVM trained using all fragmentation energies, we provide classifier weights for PubChem (“PC”), biomolecular structure database with one candidate (“bio1”) and two or more candidates (“biodb2+”). Clearly, features that require at least two candidates cannot be used for classifier “bio1”. *Unless explicitly stated otherwise*, we consider the candidate list from PubChem for the PubChem classifier, and the candidate list from the biomolecular structure database for the biomolecular structure classifier. CSI:FingerID scores are Modified Platt score from [15] and Covariance score from [69]. Multiple structures in the candidate list represented by the same fingerprint were treated as a single entry. Column ‘Δ’ shows if we enforced a feature to have positive (‘P’) or negative (’N’) weight in the classifier. Features are individually normalized

| # | Δ | Classifier weights |  |  | Name | Description |
| --- | --- | --- | --- | --- | --- | --- |
|  |  | PC | bio2+ | bio1 |  |  |
| 1 | P | 0.1167 | 0.1452 |  | Log Score Diff. 1 | Difference between log scores of highest-scoring vs. runner-up candidate, Modified Platt score |
| 2 | P | 0.0386 | 0.0000 |  | Score Diff. 1 | Difference between scores of highest-scoring vs. runner-up candidate, Modified Platt score |
| 3 | P | 0.1167 | 0.1104 |  | Log Score Diff. 2 | Difference between log scores of highest-scoring vs. runner-up candidate, Covariance score |
| 4 | P | 0.0316 | 0.0013 |  | Score Diff. 2 | Difference between scores of highest-scoring vs. runner-up candidate, Covariance score |
| 5 | P | 0.034 | 0.0861 | 0.0424 | Modified Platt Score | Modified Platt score of highest-scoring candidate |
| 6 | P | 0.0000 | 0.0139 | 0.0000 | Covariance Score | Covariance score of highest scoring candidate |
| 7 | N | −0.0096 | −0.0701 | −0.2975 | Calibrated Mod. Platt | Calibrated score of highest scoring candidate using Modified Platt scores |
| 8 | N | 0.0000 | −0.0237 | −0.1039 | Calibrated Covariance | Calibrated score of highest scoring candidate using Covariance scores |
| 9 | P | 0.0149 | 0.0185 | 0.0137 | FT Explained Intensity | Sum of normalized peak intensities in the input spectrum which are “explained” by the SIRIUS fragmentation tree |
| 10 | N | −0.0900 | −0.0753 |  | Log No. Candidates | Logarithm of candidate list size |
| 11 | P | 0.0000 | 0.0000 | 0.0000 | FT Score | Score of the SIRIUS fragmentation tree |
| 12 | P | 0.0000 | 0.0000 | 0.0000 | Fingerprint Quality | “Quality” of the predicted fingerprint, measured as $\sum_i \max\{1 - p_i, p_i\}$ for predicted fingerprint $(p_1, \dots, p_n)$ |
| 13 | N | 0.0000 | −0.0423 |  | Tanimoto Sim. runner-up | Tanimoto Similarity between highest-scoring and runner-up candidate |
| 14 | P | 0.0000 | 0.0012 | 0.0000 | Tanimoto Sim. predicted | Tanimoto Similarity between predicted fingerprint and highest-scoring candidate fingerprint |
| 15 | P | 0.0234 | 0.0178 | 0.0449 | FP Length Pred. | Cardinality of predicted fingerprint, only properties with posterior probability at least 0.5 are counted |
| 16 | P | 0.0314 | 0.0569 | 0.1178 | FP Length Hit | Cardinality of highest ranked candidate’s fingerprint |
| 17 | P | 0.0480 | 0.0907 | 0.0718 | Rescoring 1 | Score of the highest-scoring covariance candidate when scored with the Modified Platt scoring method |
| 18 | P | 0.0000 | 0.0243 | 0.0000 | Rescoring 2 | Score of the highest-scoring Modified Platt candidate when scored with the covariance scoring method |
| 19 | N | −0.0294 | 0.0000 | 0.0000 | Rescoring Calibrated | Calibrated score of highest scoring Modified Platt scoring candidate, when scored with the covariance scoring |
| 20 | P |  | 0.0119 | 0.0219 | Score Diff Bio Pub | Score difference of the top hit in the biomolecular structure database and PubChem, Modified Platt scoring |
| 21 | P |  | 0.0133 | 0.0257 | Score Diff Bio Pub | Score difference of the top hit in the biomolecular structure database and PubChem, covariance scoring |
| 22 |  |  | −0.1119 | −0.4195 | Log No. Cand. PC | Logarithm of candidate list size in PubChem |

**Supplementary Table 2:** The 626 unique structure annotations for bile acid conjugates after merging entries with identical annotated structure. Here, “rank ID” refers to the highest-scoring entry in the uncontracted list (Supplementary Material 3), and “COSMIC score” is the score of this entry; “occurrences” is the number of occurrences of this structure in the uncontracted list. The “PubChem CID” is the PubChem compound identifier number; it is empty for novel bile acid conjugates, in which case the row is colored green. If there exist multiple PubChem entries for some structure, we report the smallest PubChem CID. Next, “molecular formula” shows the molecular formula, “InChI key” and “SMILES” the structure of the bile acid conjugate. Column “adducts” indicates all those adduct types for which the corresponding structure was annotated, whereas the project space identifier (“PS ID”) can be used to access the specific SIRIUS project space folder containing input spectrum as well as SIRIUS, CSI:FingerID and COSMIC results (see Supplementary Material 3). COSMIC score “N/A” indicates that there were less than five structures in PubChem with this molecular formula, impeding p-value and confidence score estimation. For the twelve bile acids from Fig. 6, we additionally give the rank of the bile acid within the figure, compound name and abbreviation, MSI annotation level as well as links to Metabolomics USI spectra visualization, fragmentation spectra on GNPS, the GNPS spectral matching job, the GNPS visualizer for the fold change analysis and MASST job.

**Supplementary Table 3:** The complete SIRIUS project space for the bile acid conjugate annotation process with all SIRIUS, CSI:FingerID and COSMIC results. “Compound_identifications.tsv” contains the full list of COSMIC annotations before duplicate removal.

**Supplementary Table 4:** Details on the 315 novel molecular structures not contained in HMDB annotated in the human dataset. Reported are identification number (ID) and molecular formula; the “COSMIC score” is the maximum confidence score among all annotations with this structure, and “occurrences” is the number of such annotations. The “PubChem CID” is the PubChem compound identifier number; if there exist multiple PubChem entries for some structure, we report the smallest PubChem CID. Entries “InChI” and “SMILES” give the structure, entry “name” gives the chemical name of the structure (“null” is shown in case there is no name present in our database for that structure). Column “adduct” indicates the adduct type for which the corresponding structure was annotated.

**Supplementary Table 5:**
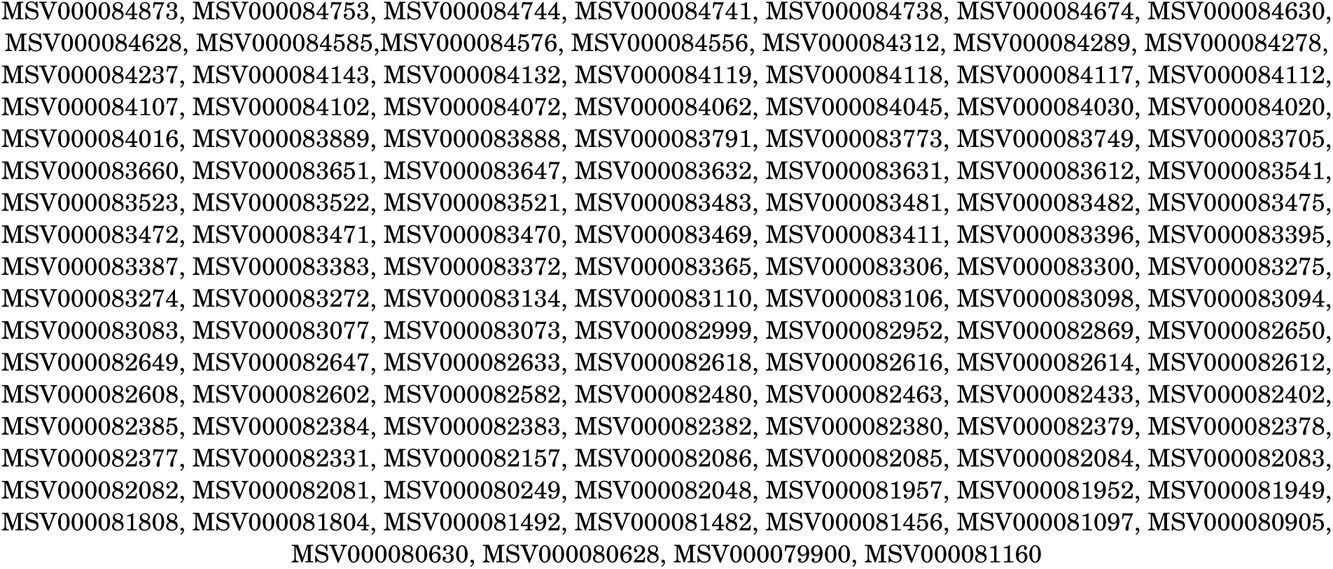
MassIVE accession numbers for the Orbitrap dataset. Corresponding mzML/mzXML files are available from MassIVE (https://massive.ucsd.edu/).

**Supplementary Table 6:** Details on the 1,715 novel molecular structures annotated in the Orbitrap dataset. Reported are identification number (ID) and molecular formula; the “COSMIC score” is the maximum confidence score among all annotations with this structure, and “occurrences” is the number of such annotations. The “PubChem CID” is the PubChem compound identifier number; if there exist multiple PubChem entries for some structure, we report the smallest PubChem CID. Entries “InChI” and “SMILES” give the structure, entry “name” gives the chemical name of the structure (“null” is shown in case there is no name present in our database for that structure). Column “adduct” indicates the adduct type for which the corresponding structure was annotated.

## Supplementary Figures

**Supplementary Fig. 1:**
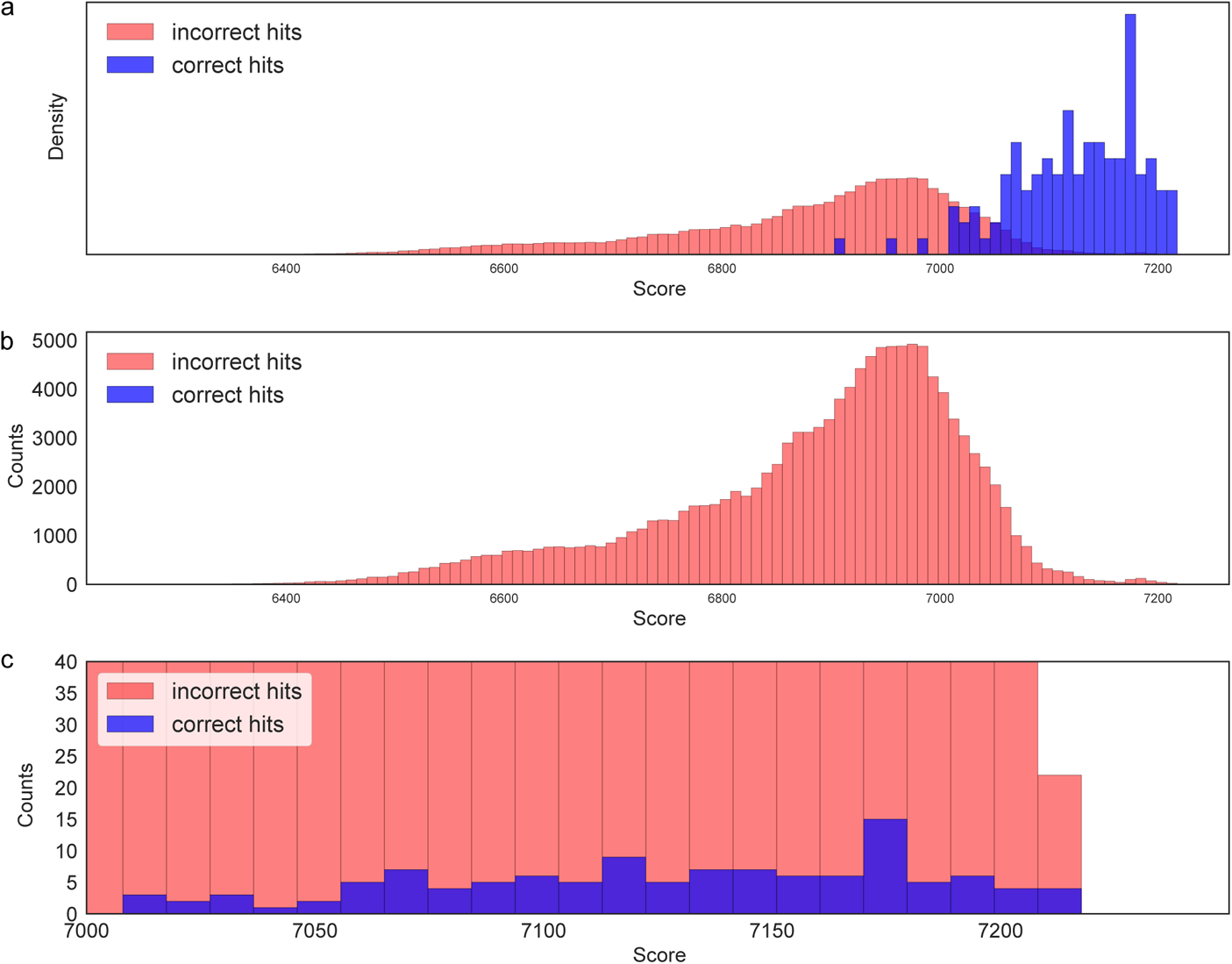
Score distribution of correct and incorrect molecular structure candidates, using CASMI 2016 contest results for CSI:FingerID. Histogram plots displaying all queries and *all candidates* of CASMI 2016 (positive ion mode) simultaneously. There are 120 correct candidates but 123 551 incorrect candidates, so incorrect candidates are three orders of magnitude more common. (a) Score distributions when both distributions have been normalized individually. Correct candidates receive a much higher score than a randomly selected incorrect candidate; if this was not the case, then CSI:FingerID would not be able to reach a reasonable annotation rate. (b) Score distributions without normalization; correct candidates are practically invisible in this plot. (c) Zoom-in into (b): We observe numerous incorrect candidates with scores as high as correct candidates.

**Supplementary Fig. 2:**
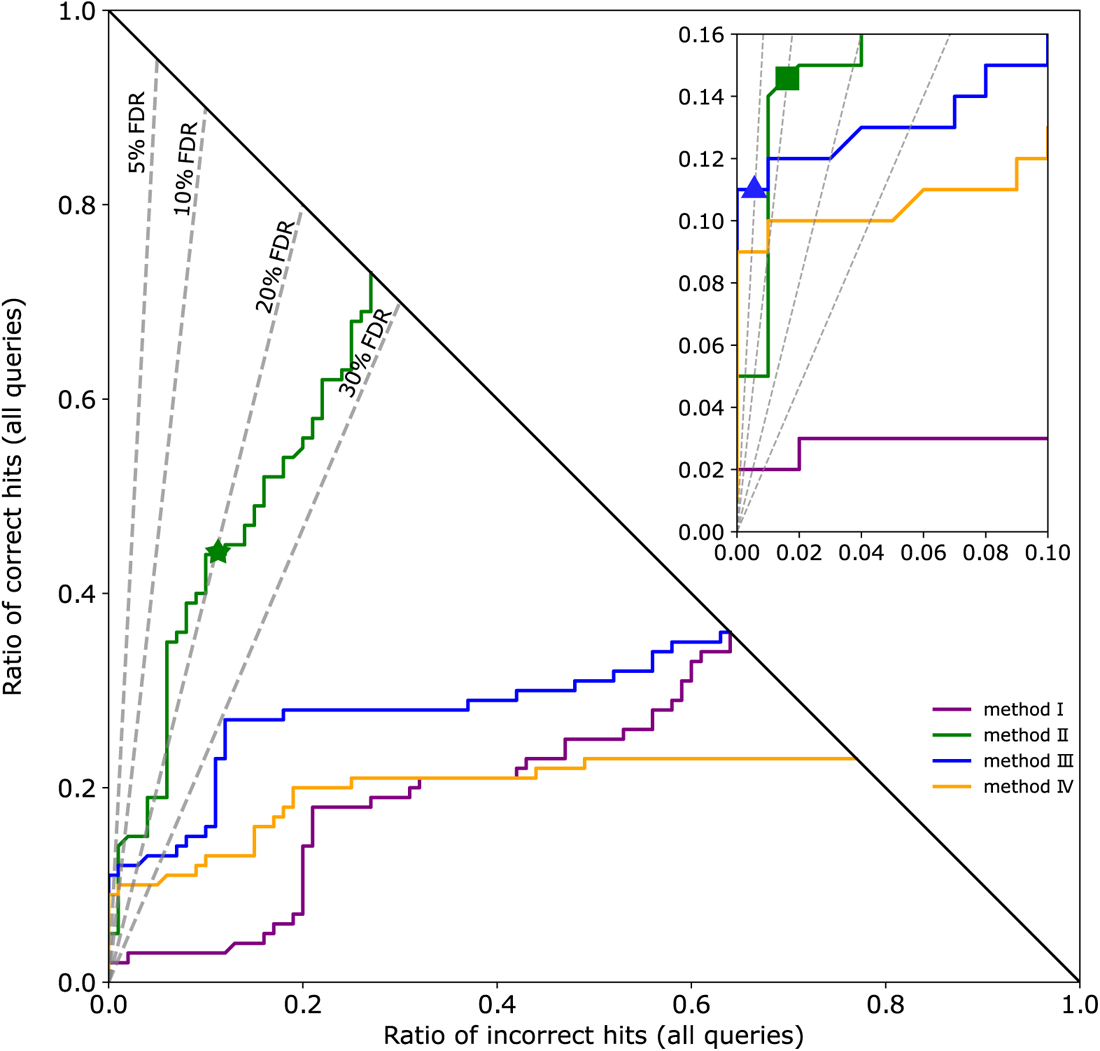
Introducing hop plots. Hop plots allow us to simultaneously assess a methods annotation rate *and* its power to separate correct and incorrect hits. Two methods with identical annotation rate will end up in the same point (*x*, *y*) with *x y* 1, see methods I and III; these methods can differ substantially in their separation power. The plot shows which method performs best for a desired number of correct annotations (horizontal lines, not shown), incorrect annotations (vertical lines, not shown), or false discovery rate (FDR, dashed lines). For example, if we are willing to accept three incorrect annotations from a total of *N* = 100 queries, then method IV clearly outperforms method III; this ordering is reversed if we consider all queries (*x y* 1). FDR levels correspond to lines through the origin; a hop curve may cross or touch some FDR line multiple times, or only in the origin. We report the maximum number of correct annotations among all crossing points. For example, method II returns 55 hits (44 correct, 11 incorrect) at FDR 20 % (star). We are usually interested in small FDR values such as FDR 10 %, so a zoom-in shows where different curves cross the corresponding FDR lines: For example, method III returns 11 hits (all correct) at FDR 5 % (triangle, zoom-in), and method II returns 15 hits (14 correct) at FDR 10 % (square, zoom-in). See Methods for further details.

**Supplementary Fig. 3:**
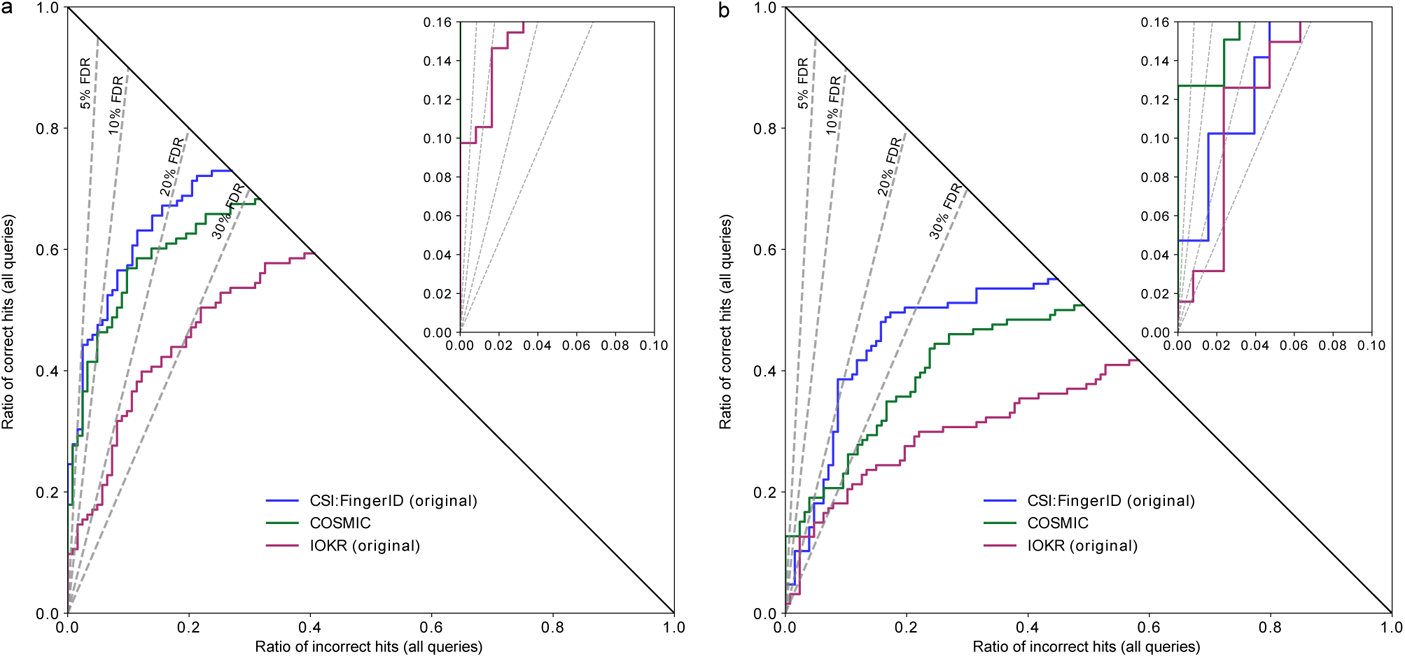
Separation for CSI:FingerID on CASMI 2016 without structure-disjoint evaluation. Hop plots for searching (a) the biomolecule structure database, *N* = 124 or (b) ChemSpider, *N* = 127. FDR levels shown as dashed lines. Positive ion mode. “CSI:FingerID (original)” refers to the original CASMI 2016 submission of CSI:FingerID, and “IOKR (original)” is the original submission for its Input Output Kernel Regression (IOKR) variant. In contrast to Fig. 2, these evaluations were not carried out structure-disjoint; CASMI is a blind competition, and the correct answers were unknown to the contestants upon submission. The COSMIC curve is given for comparison; it is copied verbatim from Fig. 2 and *ensures structure-disjoint evaluation*. In agreement with our findings for applying COSMIC without structure-disjoint evaluation (Fig. 5), separation of the original submissions is much better than for CSI:FingerID with structure-disjoint evaluation; but notably, not better than for COSMIC *with* structure-disjoint evaluation. Hence, we must attribute the separation power of the original submissions mostly to the overlap in structures between training and evaluation data: Structures for which a fragmentation spectrum is present in the training data often receive high hit scores, comparable to library searching.

**Supplementary Fig. 4:**
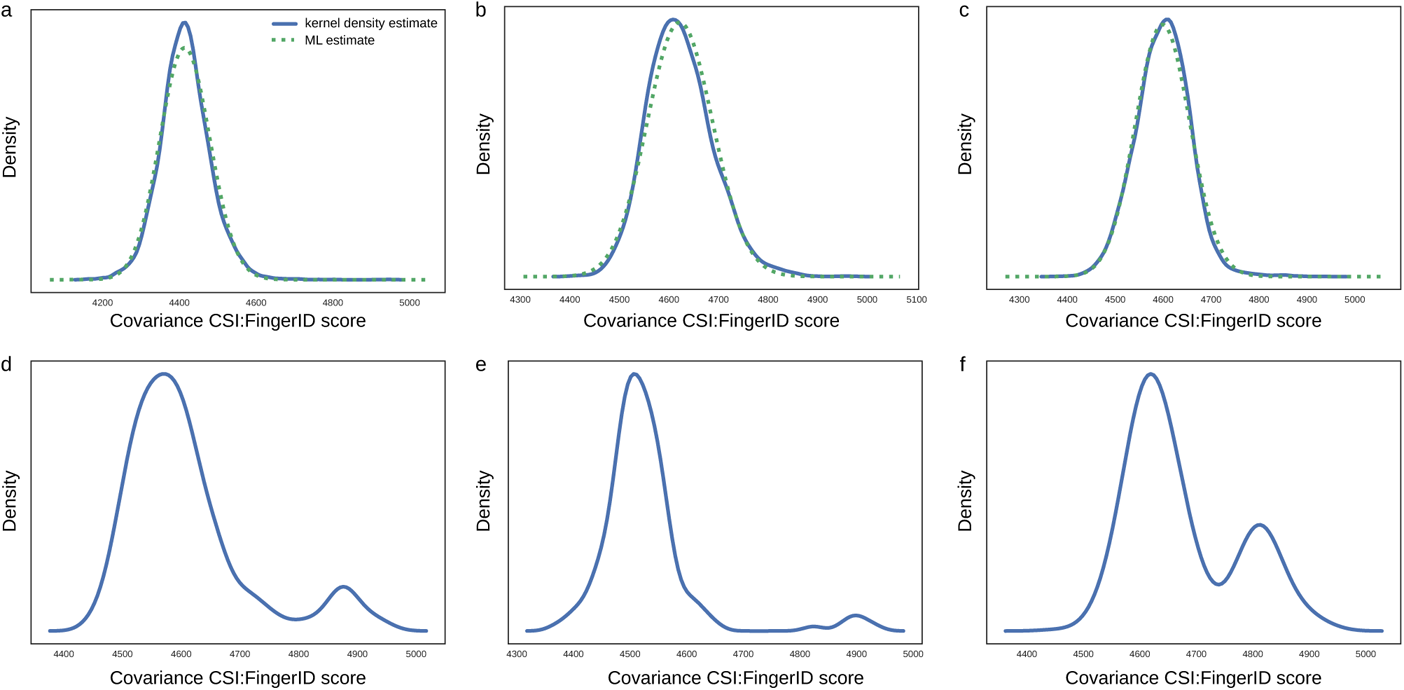
Examples of CSI:FingerID score distributions. Shown are kernel density estimates of candidate scores searching in PubChem. We find that unimodal score distributions (a–c) are often similar to a log-normal distribution (“kernel density estimate”); for comparison, we show the log-normal distribution with parameters fitted by Maximum Likelihood estimation (“ML estimate”). Other score distributions are clearly multimodal (d–f). (a) PyroGlu-Trp, C_16_H_17_N_3_O_4_, 4 862 candidates, NIST 1632483. (b) 3-Methyl-L-histidine, C_7_H_11_N_3_O_2_, 3 503 candidates, NIST 1346484. (c) 1,3-Benzodioxole-5-propanamine, C_12_H_17_NO_2_, 15 786 candidates, NIST 1306465. (d) N-(2-Hydroxyethyl)-5(6)-epoxy-8Z,11Z,14Z-eicosatrienamide, C_22_H_37_NO_3_, 471 candidates, NIST 1139175. (e) Methanone, C_24_H_22_FNO_2_, 156 candidates, NIST 1300971. (f) Benzeneethanamine, C_18_H_22_BrNO_3_, 483 candidates, NIST 1380115. Numbers of candidates from PubChem.

**Supplementary Fig. 5:**
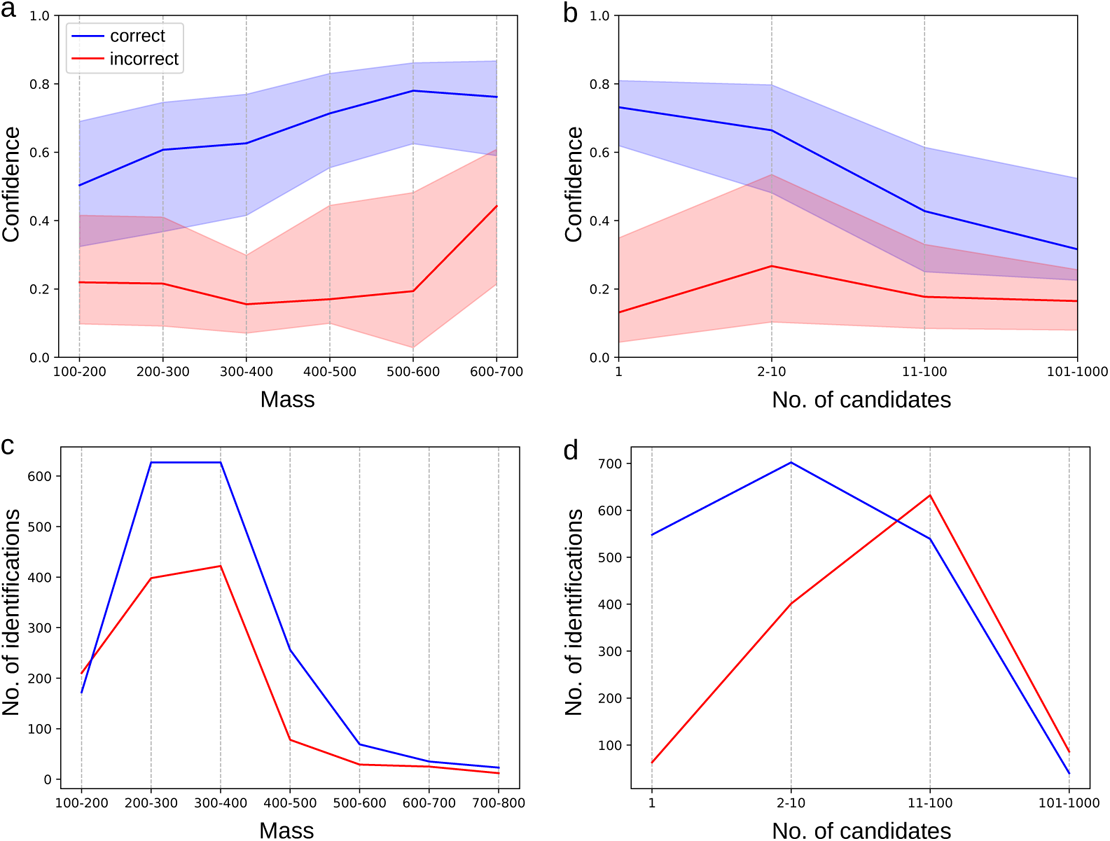
Effect of query compound mass and number of candidates on confidence scores. Independent data, merged spectra (*N* = 3013), structure-disjoint evaluation, medium noise, biomolecule structure database. (a,b) Confidence score of correct and incorrect annotations when varying query mass ranges (a) and number of candidates (b). Solid lines show median values, colored areas indicate first (25 %) and third (75 %) quartiles. (c,d) Number of correct and incorrect annotations for varying query mass ranges (c) and number of candidates (d). One compound with mass below 100 Da omitted from (a,c). Only few compounds exist above 500 Da and with more than 100 candidates, so curves (a,b) should be interpreted with care in these regions.

**Supplementary Fig. 6:**
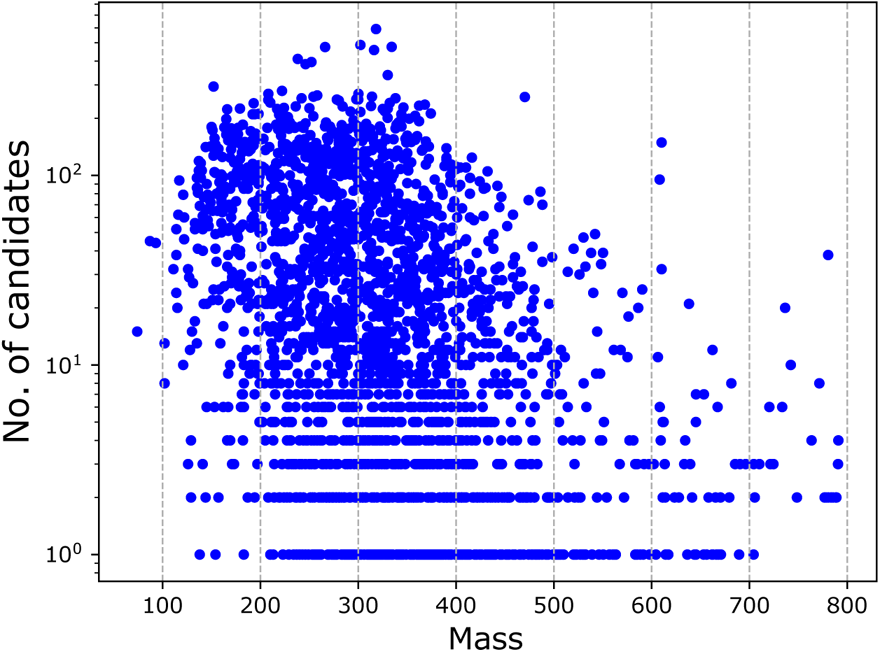
Correlation between the mass of a query compound and the number of candidates retrieved from the database. Note the logarithmic y scale. We display exactly the candidates from Supplementary Fig. 5: Independent data, merged spectra, structure-disjoint evaluation, medium noise, biomolecule structure database.

**Supplementary Fig. 7:**
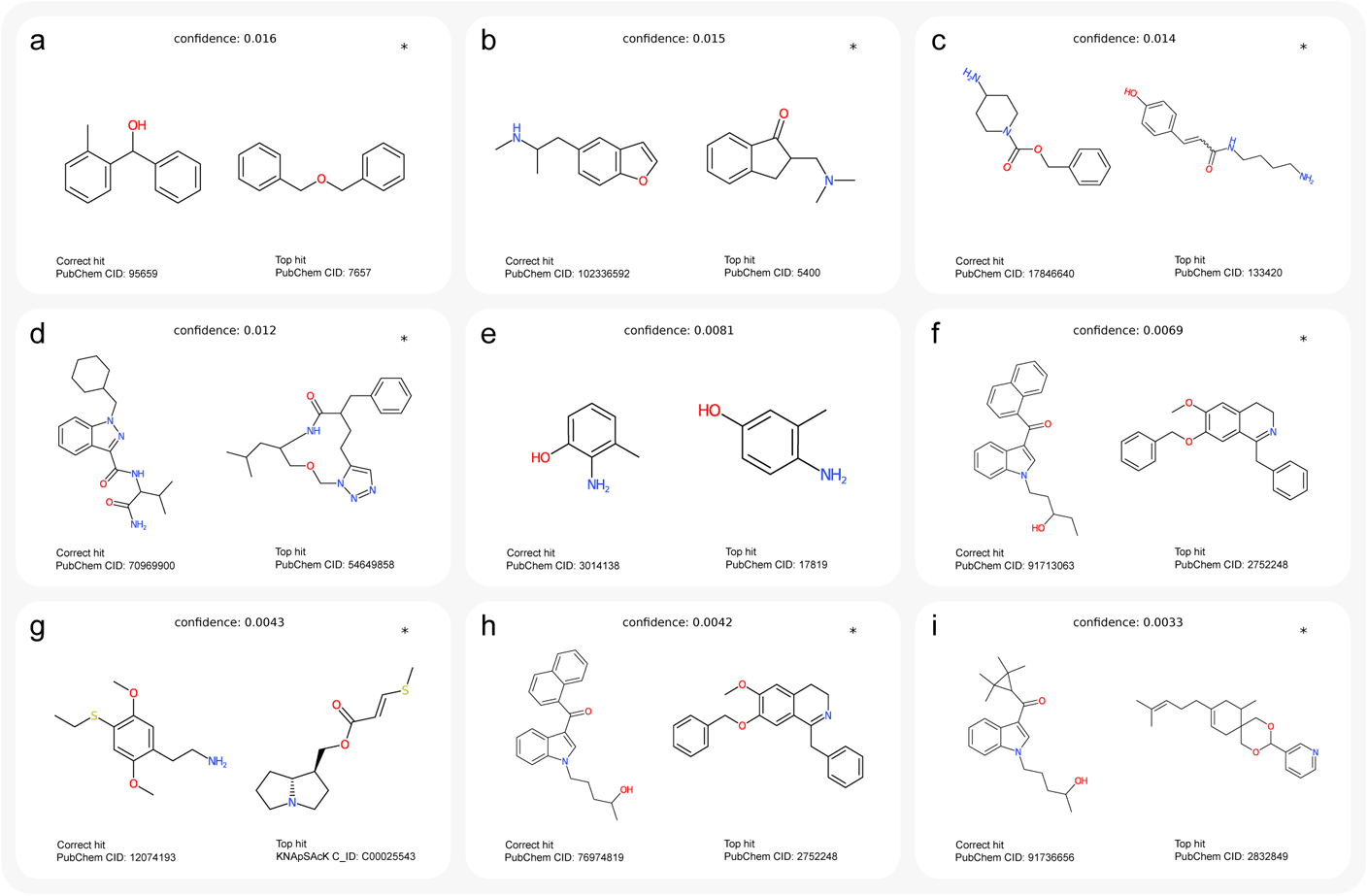
Examples of incorrect annotations with lowest confidence scores. Queries are cross-validation data, merged spectra, medium noise, biomolecule structure database, structure-disjoint evaluation. (a–i) Incorrect hits with lowest confidence scores. Top-ranked structure on the left and corresponding correct structure on the right. “PubChem CID” is PubChem compound identifier number. Instances where the correct structure was not contained in the biomolecule structure database are marked by an asterisk. For (g), the structure of the top hit is not contained in PubChem; we report the KNApSAcK compound identifier (“C_ID”) instead. For (a) and (e), molecular graphs of incorrect hit and correct structure differ by the theoretical minimum of two edge deletions. For (a), the query spectrum was heavily distorted, and only 8.6 % of peak intensities were explained by the fragmentation tree. For (e), the three top-scoring candidates — including the correct one — were structurally highly similar and received almost identical CSI:FingerID score. Hence, COSMIC rightfully showed little confidence in these (incorrect) hits. Query spectra: (a) NIST 1544714/19/23, (b) NIST 1322859/64/69, (c) NIST 1627646/51/56, (d) NIST 1462584/87/93, (e) NIST 1340388/91/96, (f) NIST 1320854/56/62, (g) NIST 1386503/07/12, (h) NIST 1305770/72/78, (i) NIST 1325235/37/43.

**Supplementary Fig. 8:**
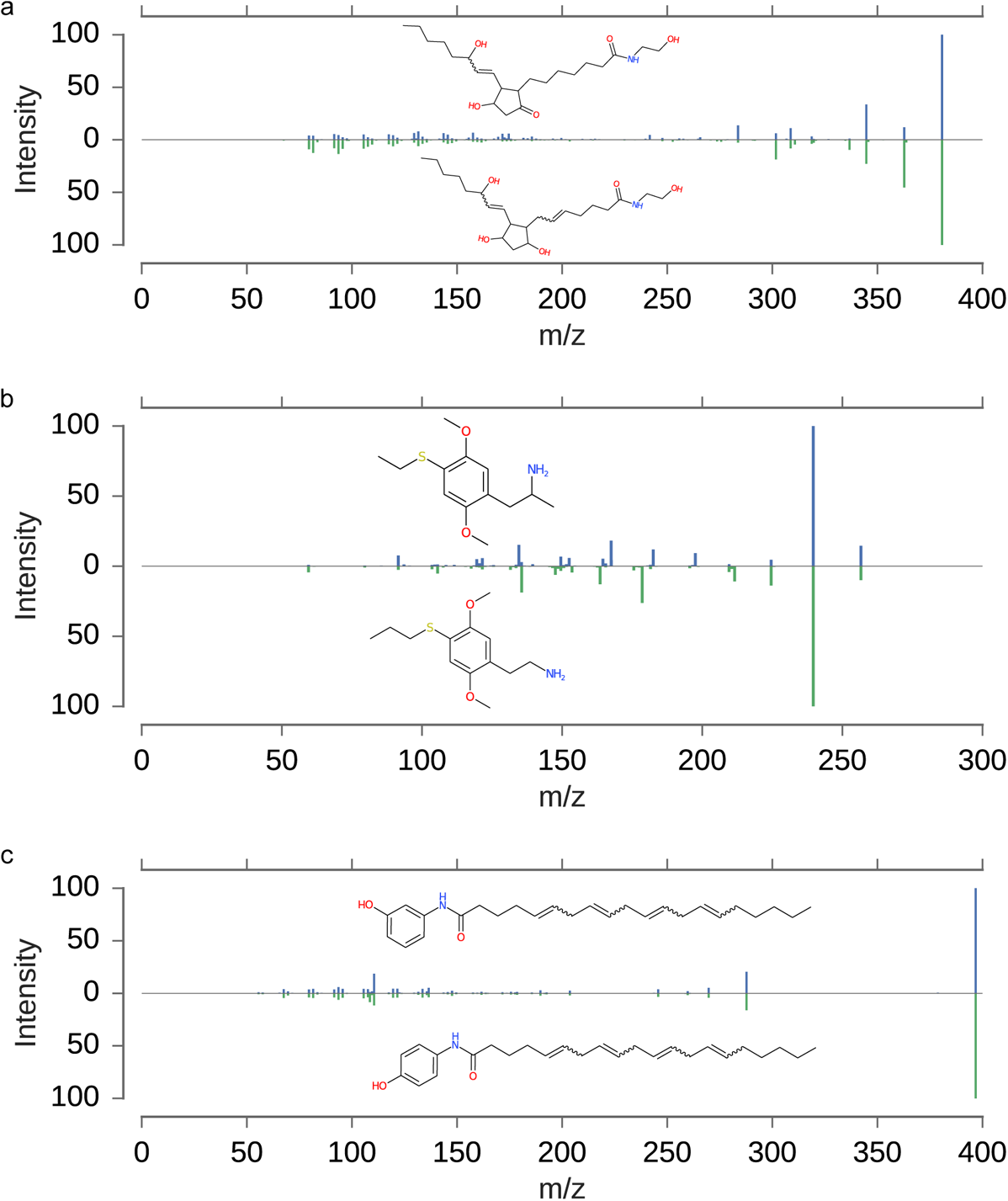
Comparison of fragmentation spectra for high-scoring incorrect hits from Fig. 4. These correspond to compound pairs where COSMIC search resulted in an incorrect hit; would spectral library search be able to avoid these incorrect annotations? For three incorrect hits from Fig. 4 (**a**,**b**,**h**) there exist merged spectra; for the remaining six incorrect hits, no such data are available. Be reminded in all three cases (**a**,**b**,**h**), the correct structure was not contained in the searched molecular structure database. Merged spectrum and structure of correct structure shown top, merged spectrum and structure of incorrect hit bottom. Merged spectra were combined from 10 eV, 20 eV and 40 eV spectra as described in the Methods section. (a) Mirror plot for Fig. 4 a, confidence 0.9596, cosine score 0.8566. Mirror plot for Fig. 4 b, confidence 0.9468, cosine score 0.9432. (c) Mirror plot for Fig. 4 h, confidence 0.8942, cosine score 0.9968. In all three cases, the cosine score is above 0.85, and would result in a high-confidence but incorrect library search annotation if one of the spectra was in the library, the other our query. For (c) we argue that no method could possibly distinguish between these structures based on the MS/MS data. Merged spectra: (a) correct NIST 1210761/62/64, incorrect hit NIST 1215622/23/27; (b) correct NIST 1617825/29/34, incorrect hit NIST 1386465/69/74; (c) correct NIST 1418771/73/80, incorrect hit NIST 1375293/295/301.

**Supplementary Fig. 9:**
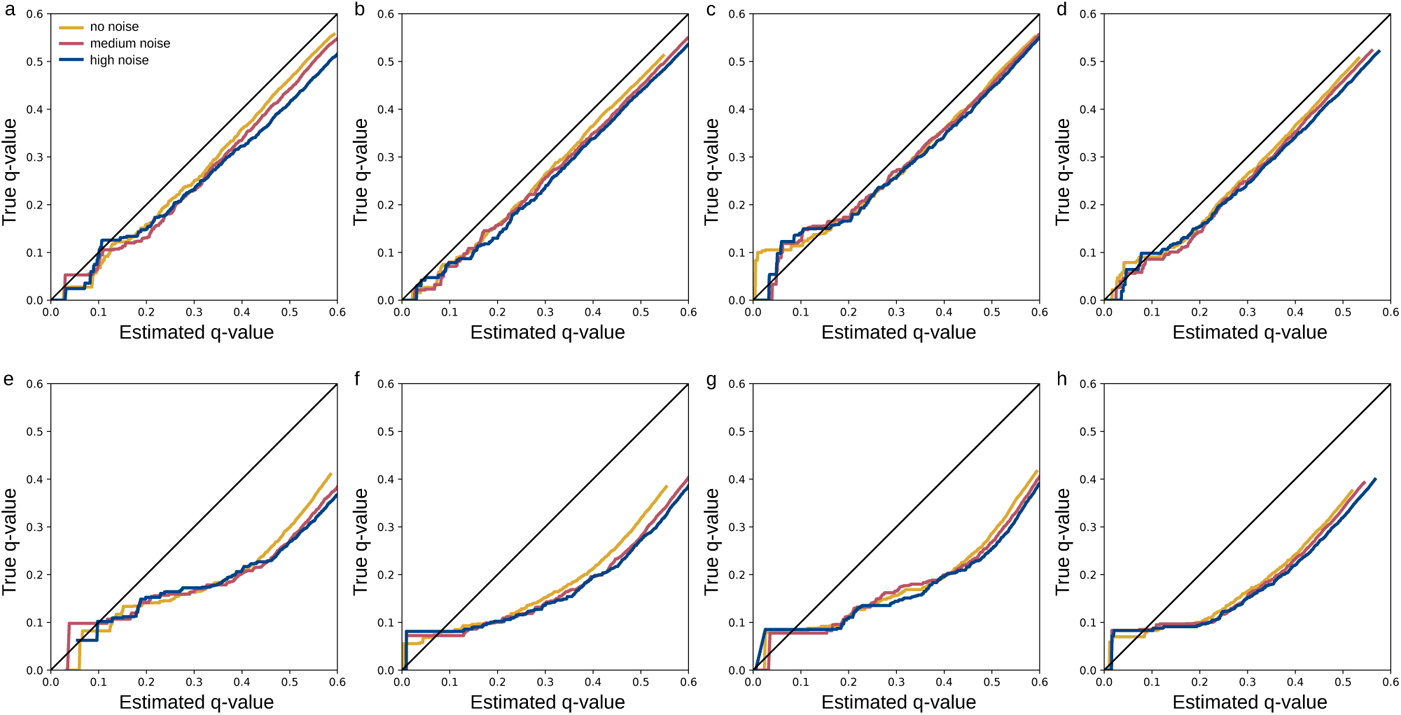
False discovery rate estimation. Q-Q plot of true vs. estimated q-values with no added noise, medium noise, and high noise. (a–d) cross-validation, *N* = 3721. (a) 10 eV, (b) 20 eV, (c) 40 eV, (d) merged spectra. (e–h) Independent data, *N* = 3013. (e) 10 eV, (f) 20 eV, (g) 40 eV, (h) merged spectra. The “step” at the beginning of most curves in (e–h) is not an issue of FDR estimation, but due to the fact that no non-zero (true) q-values below this exist in the dataset.

**Supplementary Fig. 10:**
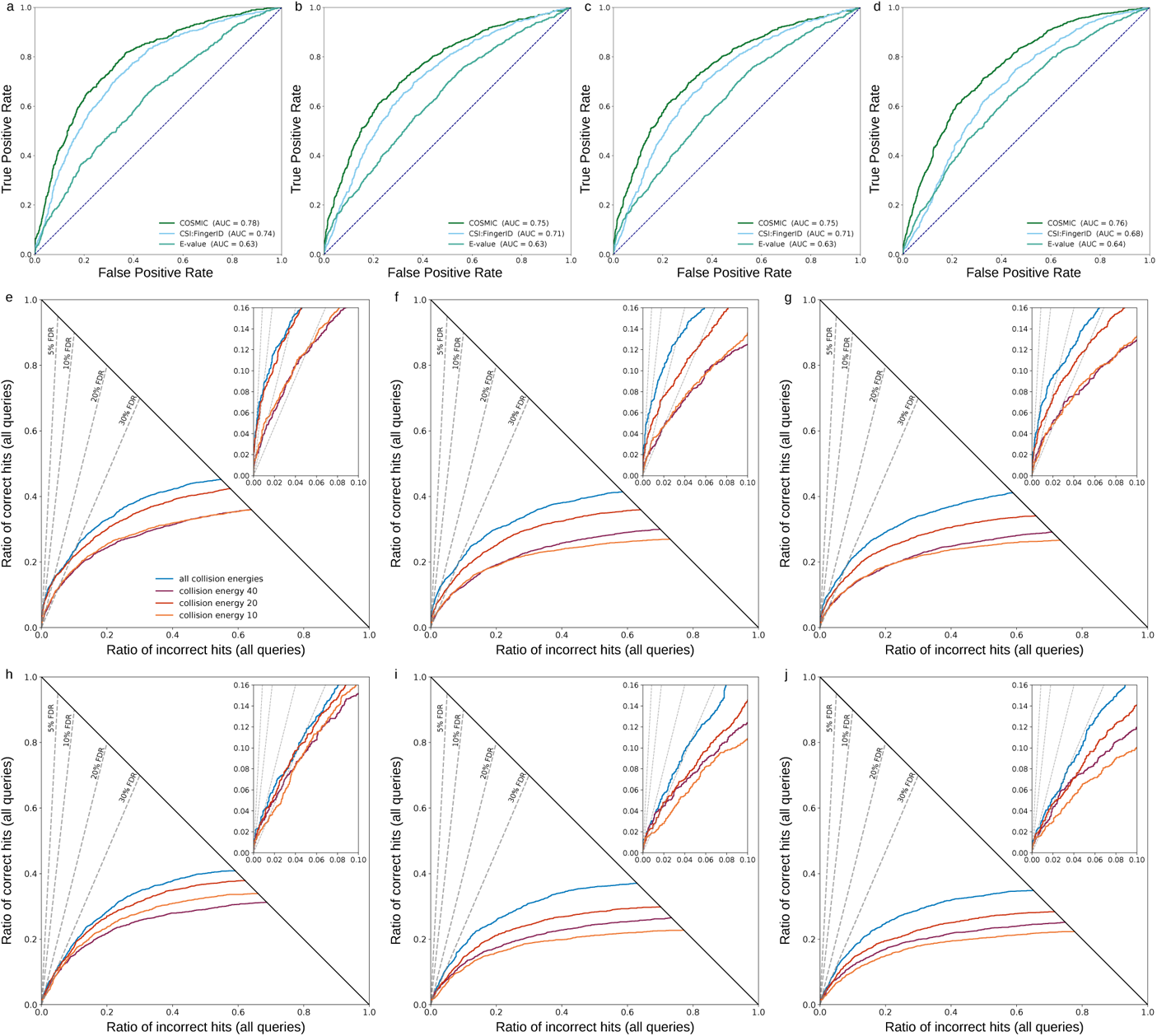
Evaluation of separation searching in PubChem. (a–d) Comparison of CSI:FingerID score, calibrated score and COSMIC confidence score. ROC curves, structure-disjoint evaluation, independent data, medium noise, *N* = 3013. (a) 10 eV, (b) 20 eV, 40 eV, (d) merged spectra (“all collision energies”). Notably, E-values sometimes result in worse separation than the CSI:FingerID score. (e–j) Evaluation of the COSMIC confidence score: Hop plots for different collision energies; notably, these result in substantially different annotation rates. (e–g) Structure-disjoint cross-validation, *N* = 3721. (h–j) Independent data with structure-disjoint evaluation, *N* = 3013. (e,h) No added noise, (f,i) medium noise, (g,j) high noise.

**Supplementary Fig. 11:**
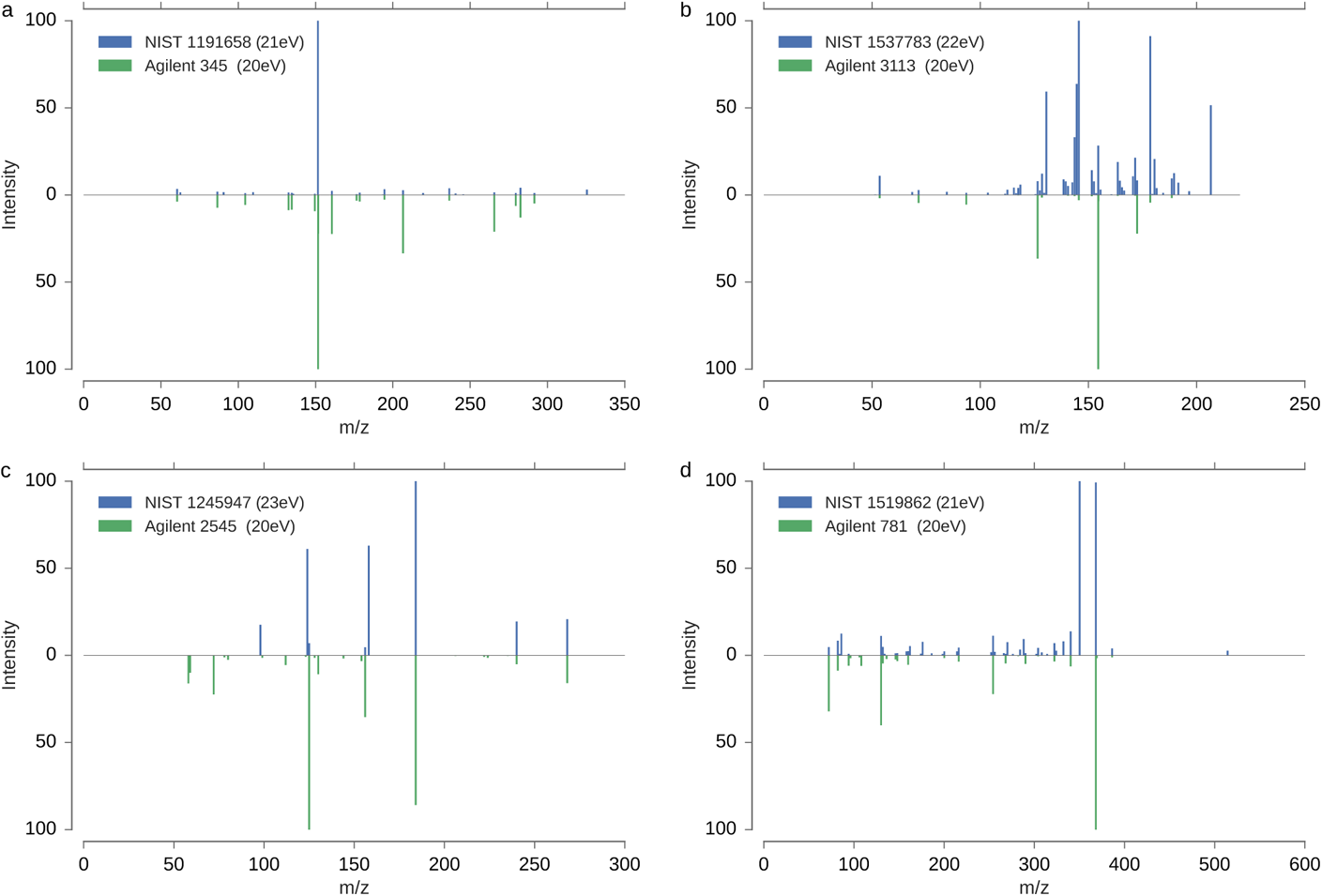
Mirror plots of low-scoring library hits that were correctly annotated with high confidence using COSMIC. Shown is the query spectrum (bottom) from the independent dataset, plus the top-scoring reference spectrum (top) from the spectral library, that is, the SVM training dataset without merging spectra. Cosine scores were calculated using regular intensities (cosine) as well as square root of intensities (cosine-sqrt). All query spectra consist of a single 20 eV collision energy measurement with medium noise added. Reference spectra consist of a single collision energy measurement with no added noise; shown is the spectrum with the *highest* cosine, among *all* spectra in the spectral library for this compound. (a) Spectra of Thiophanate, PubChem CID 3032792, molecular formula C_14_H_18_N_4_O_4_S_2_. Reference spectrum NIST 1191658, query spectrum Agilent PCDL 345. Correct COSMIC annotation with confidence 0.9092, cosine 0.0637, cosine-sqrt 0.3165. (b) Spectra of Chlorbufam, PubChem CID 16073, molecular formula C_11_H_10_ClNO_2_. Reference spectrum NIST 1537783, query spectrum Agilent PCDL 3113. Correct COSMIC annotation with confidence 0.9347, cosine 0.1949, cosine-sqrt 0.3523. (c) Spectra of Duloxetine, PubChem CID 60835, molecular formula C_18_H_19_NOS. Reference spectrum NIST 1245947, query spectrum Agilent PCDL 2545. Correct COSMIC annotation with confidence 0.9283, cosine 0.5197, cosine-sqrt 0.4767. (d) Spectra of Proscillaridin, PubChem CID 5284613, molecular formula C_30_H_42_O_8_. Reference spectrum NIST 1519862, query spectrum Agilent PCDL 781. Correct COSMIC annotation with confidence 0.9720, cosine 0.6312, cosine-sqrt 0.4852. Unlike the commercial Agilent library, the query spectra shown here are uncurated and artificial noise was added.

**Supplementary Fig. 12:**
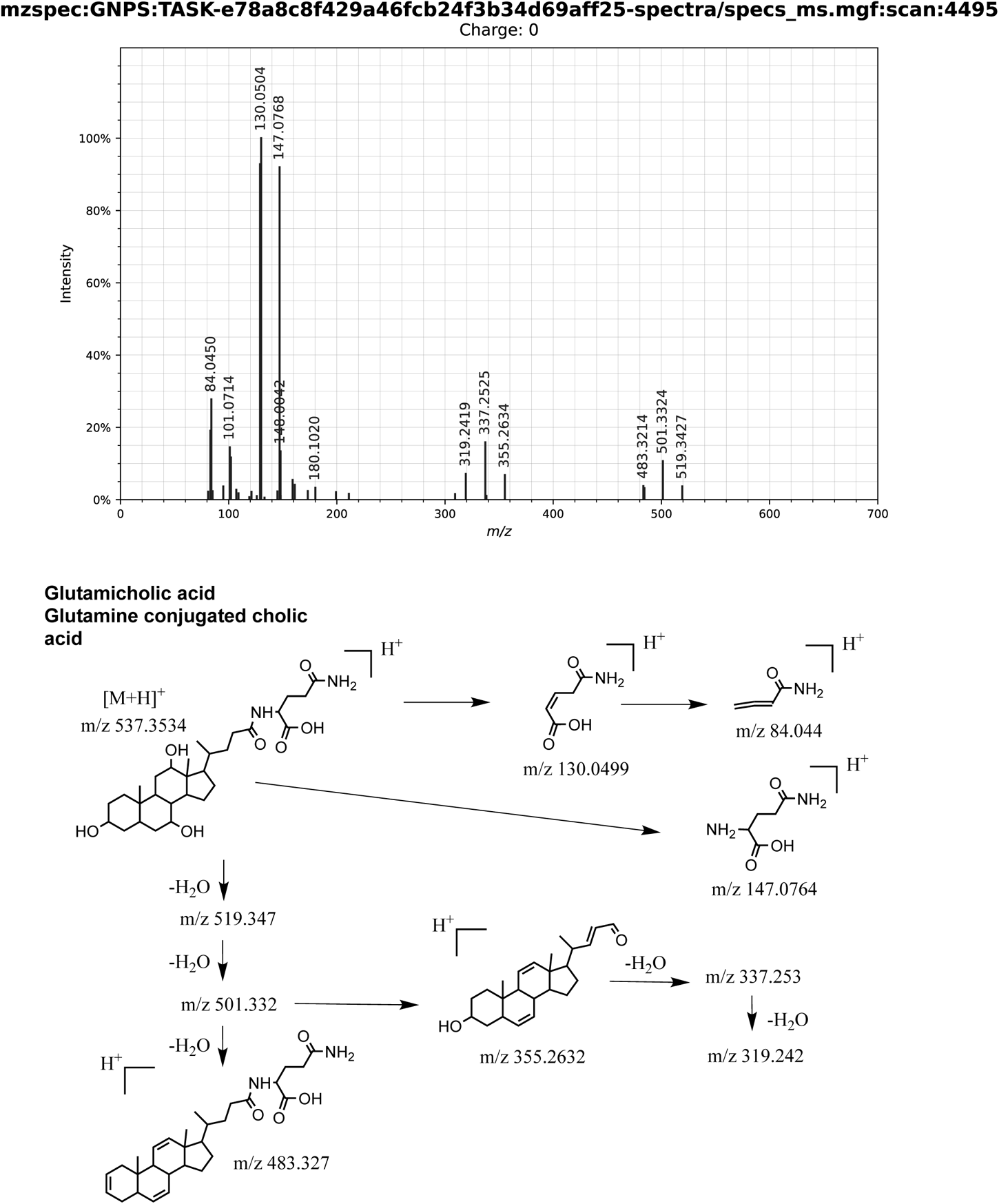
Manual fragmentation analysis of the COSMIC bile acid conjugate 1. Shown is the fragmentation spectrum of *m/z* 537.354 at 196.2 seconds. Library ID CCMSLIB00005467949, MetabolomicsUSI spectrum link.

**Supplementary Fig. 13:**
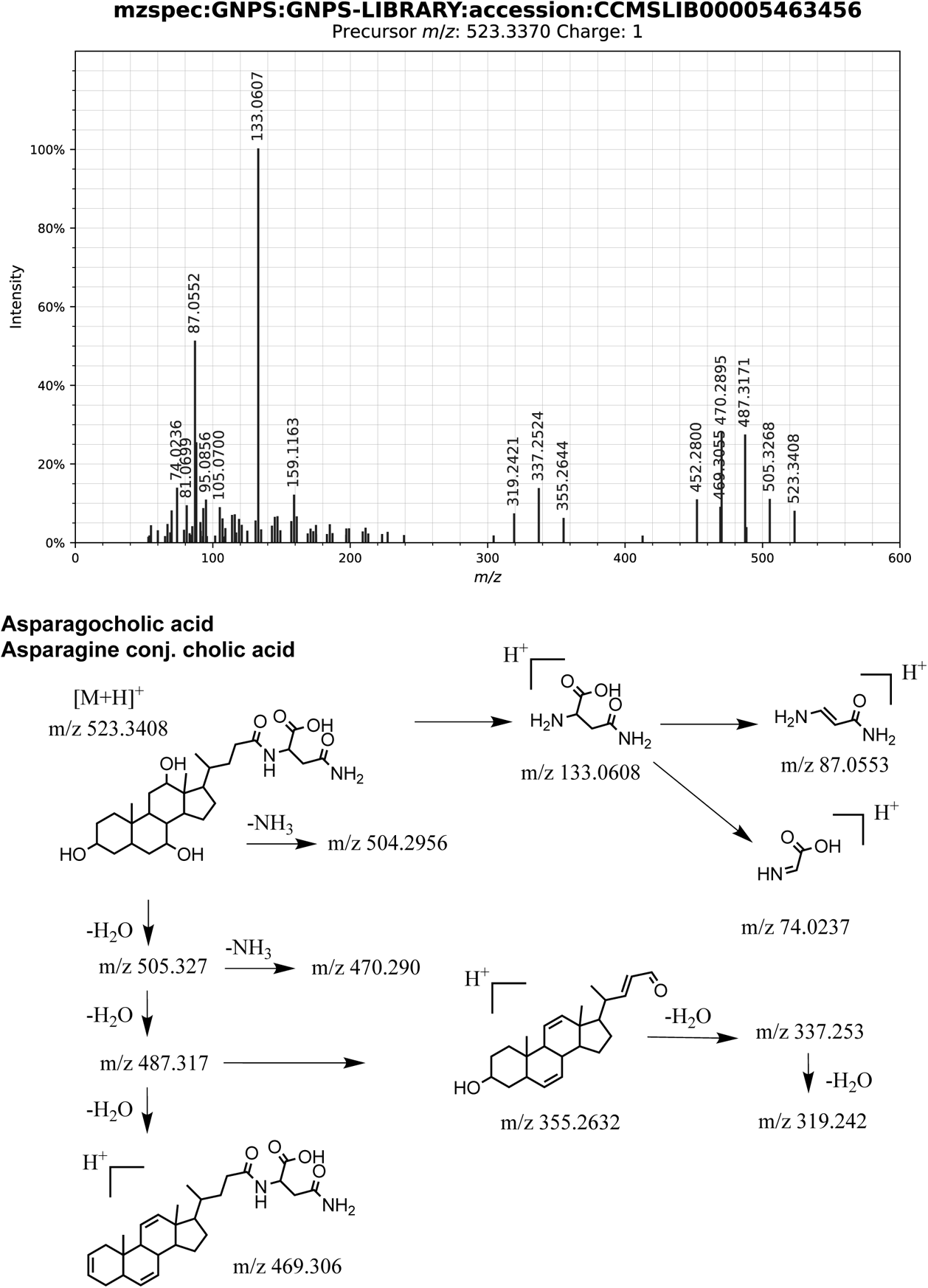
Manual fragmentation analysis of the COSMIC bile acid conjugate 2. Shown is the fragmentation spectrum of *m/z* 523.3383 at 194.6 seconds. Library ID CCMSLIB00005787996, MetabolomicsUSI spectrum link.

**Supplementary Fig. 14:**
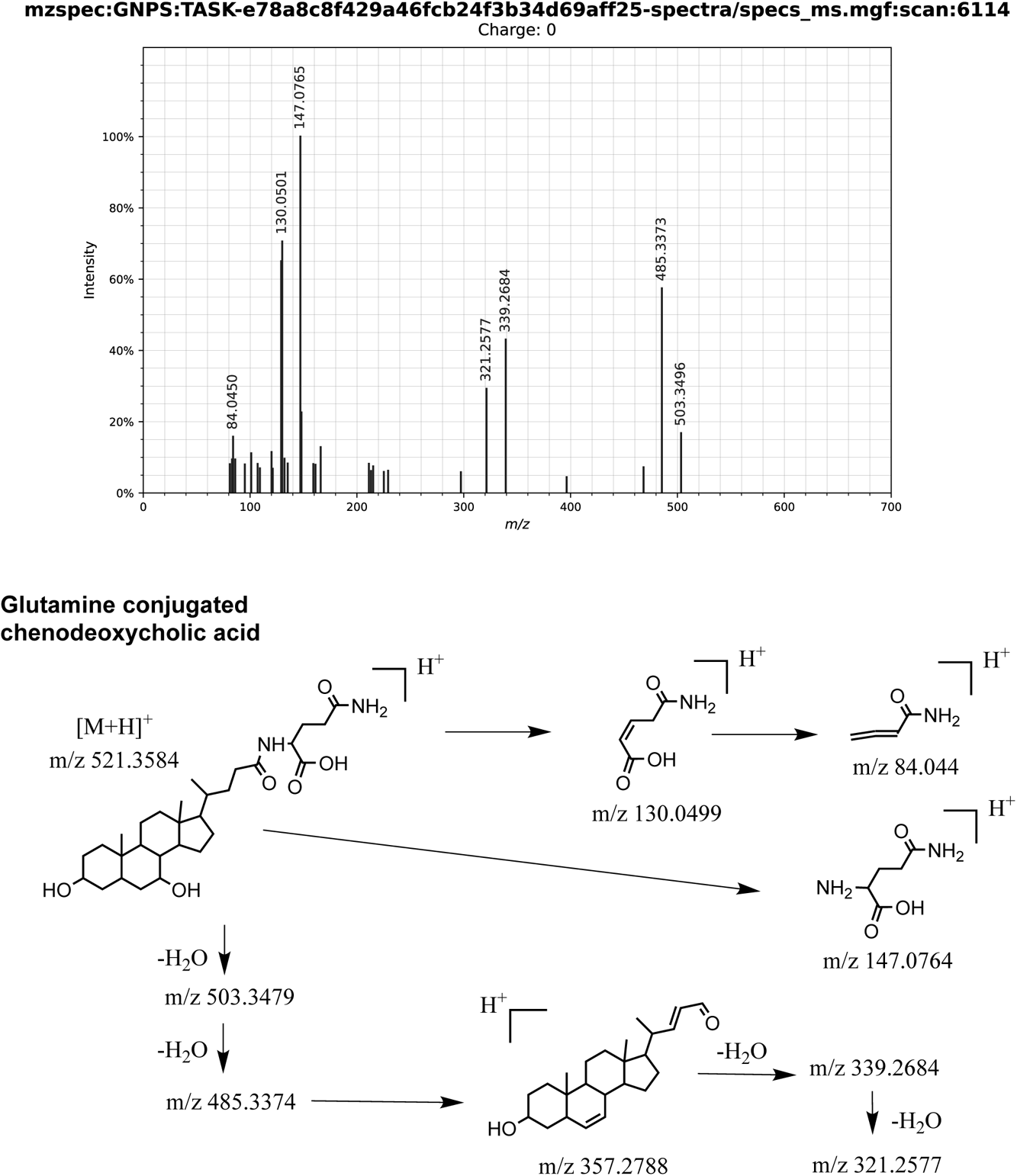
Manual fragmentation analysis of the COSMIC bile acid conjugate 3. Shown is the fragmentation spectrum of *m/z* 521.3598 at 246.3 seconds. Library ID CCMSLIB00005788119, MetabolomicsUSI spectrum link.

**Supplementary Fig. 15:**
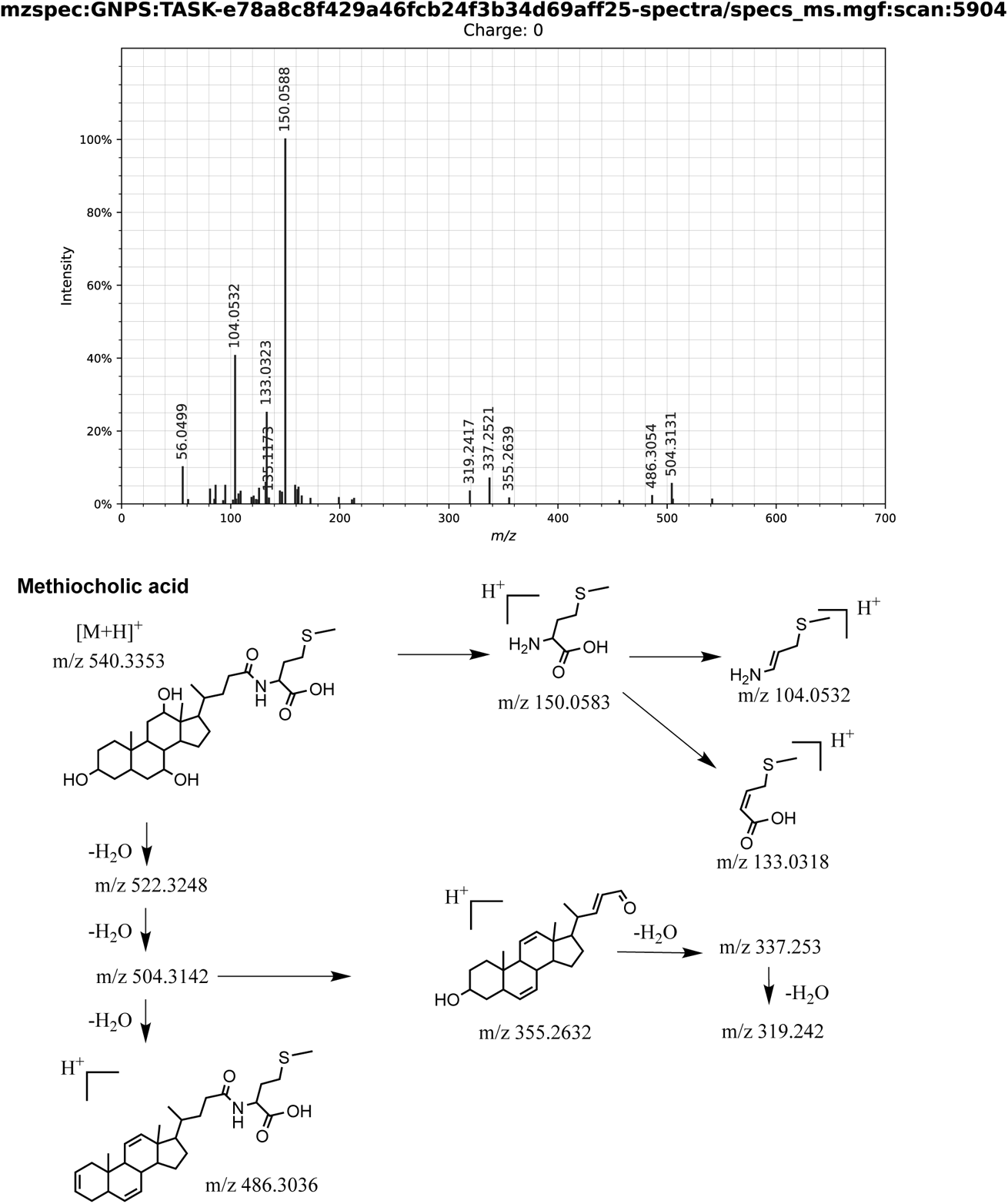
Manual fragmentation analysis of the COSMIC bile acid conjugate 4. Shown is the fragmentation spectrum of *m/z* 540.3384 at 238.71 seconds. Library ID CCMSLIB00005716809, MetabolomicsUSI spectrum link.

**Supplementary Fig. 16:**
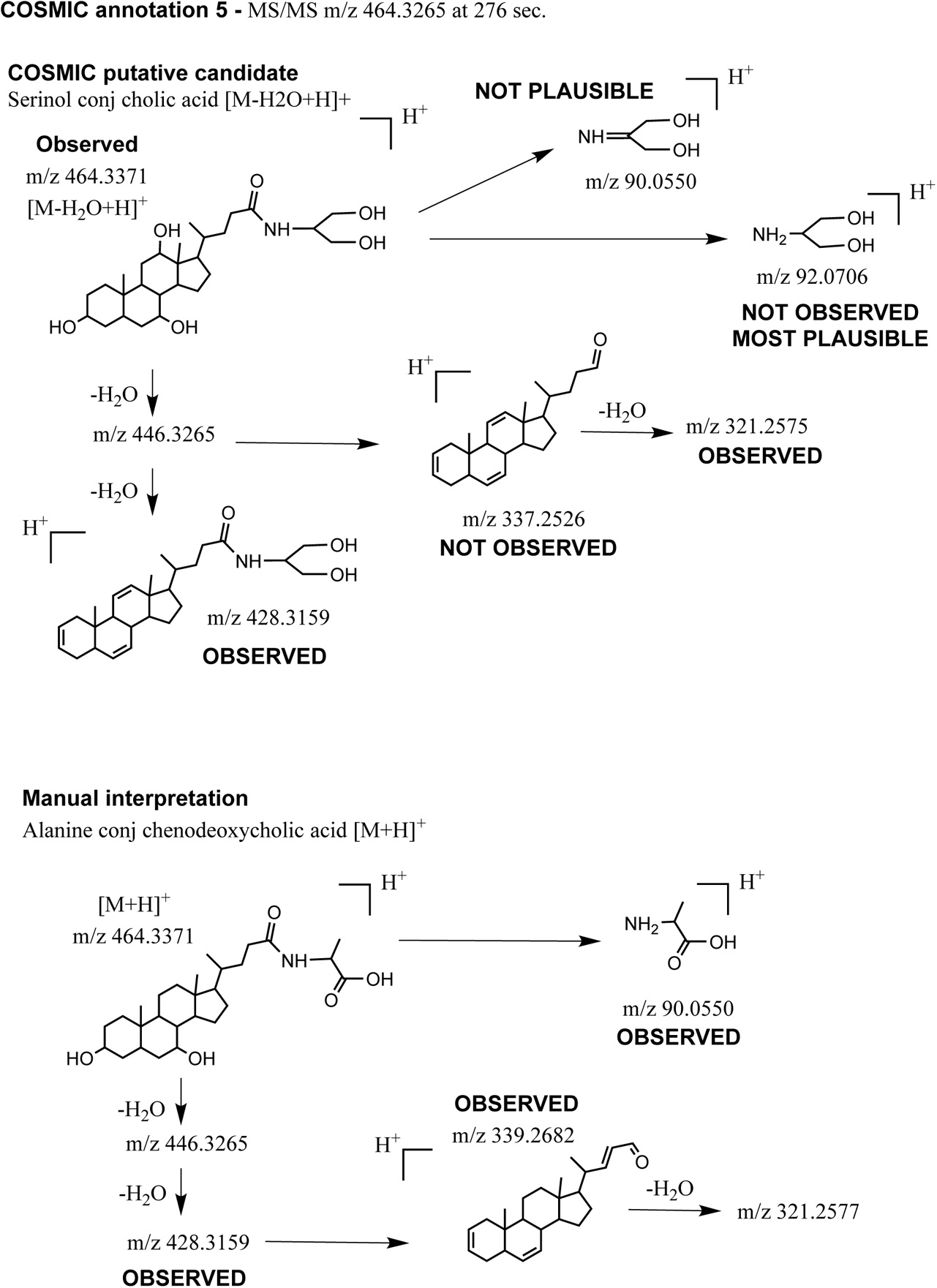
Manual fragmentation analysis of the COSMIC bile acid conjugate 5. Shown is a comparative fragmentation analysis for the fragmentation spectrum of *m/z* 464.3387 at 276.59 seconds. Fragmentation analysis for the COSMIC annotation (Serinol conjugated acid) is shown on top, interpretation of more likely structure (Alanine conjugated chenodeoxycholic acid)on bottom. Library ID CCMSLIB00005788120, MetabolomicsUSI spectrum link.

**Supplementary Fig. 17:**
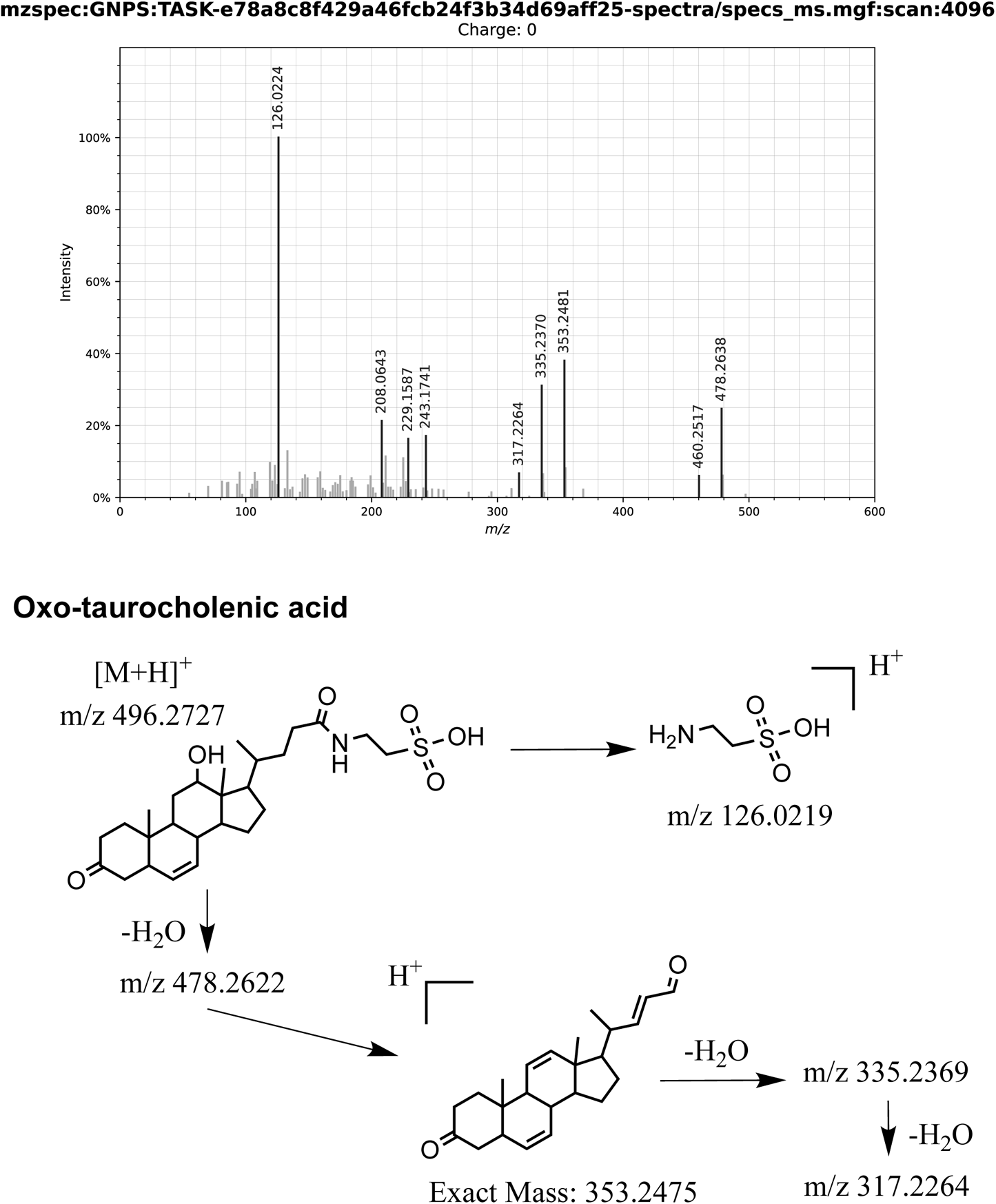
Manual fragmentation analysis of the COSMIC bile acid conjugate 6. Shown is the fragmentation spectrum of *m/z* 496.2717 at 183.2 seconds. Library ID CCMSLIB00005788121, MetabolomicsUSI spectrum link.

**Supplementary Fig. 18:**
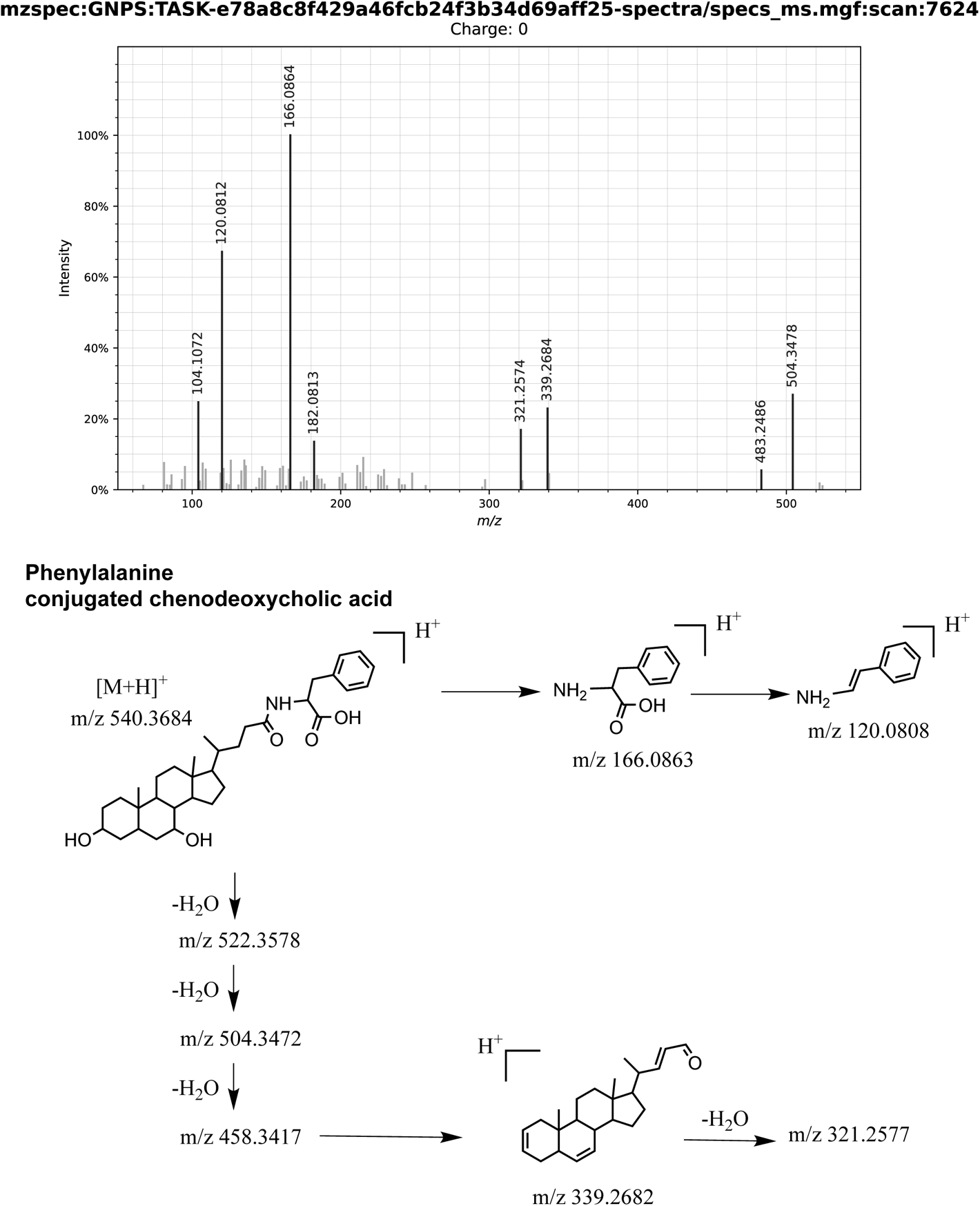
Manual fragmentation analysis of the COSMIC bile acid conjugate 7. Shown is the fragmentation spectrum of *m/z* 540.3689 at 303.0 seconds. Library ID CCMSLIB00005467952, MetabolomicsUSI spectrum link.

**Supplementary Fig. 19:**
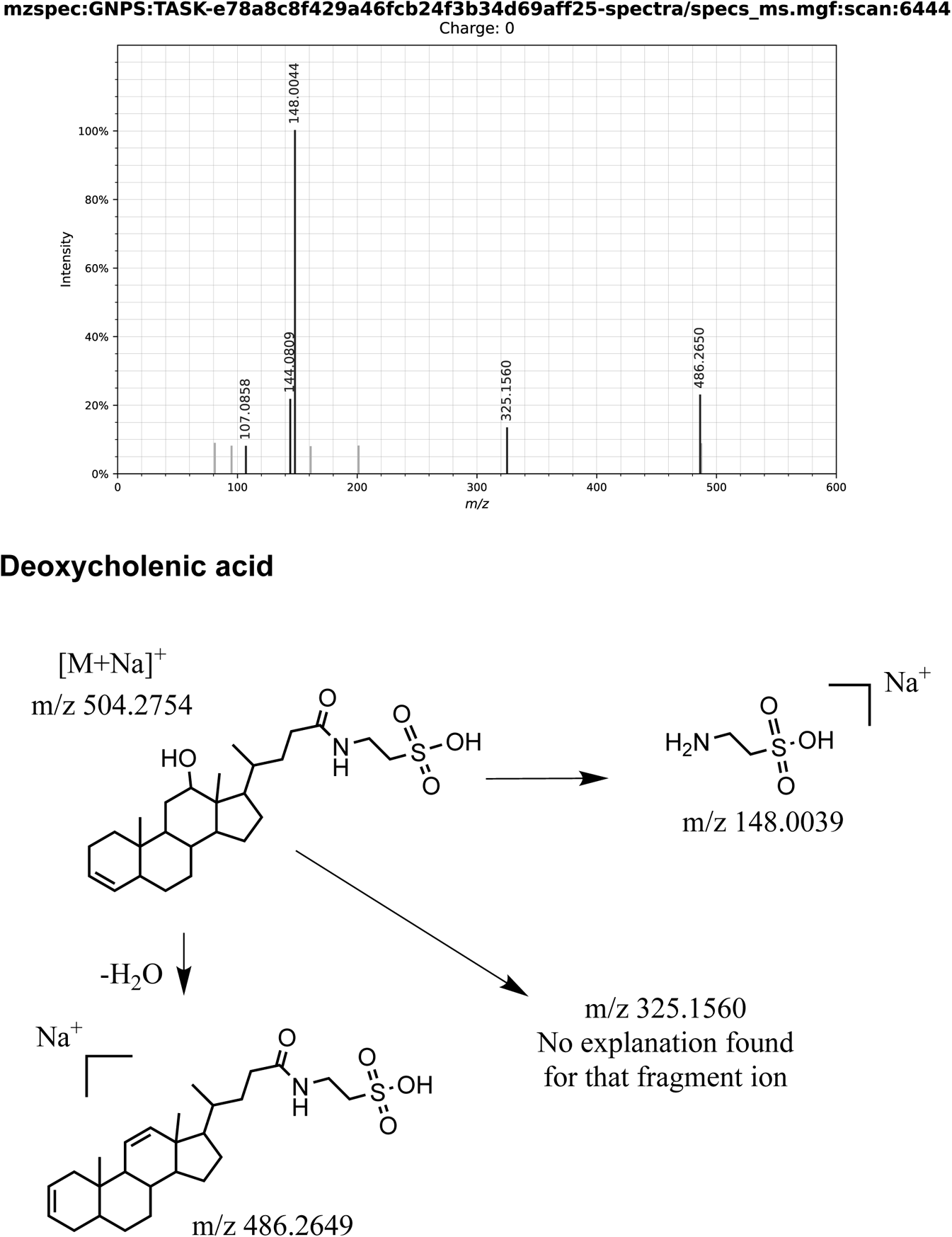
Manual fragmentation analysis of the COSMIC bile acid conjugate 8. Shown is the fragmentation spectrum of *m/z* 504.2742 at 260.1 seconds. Library ID CCMSLIB00005788122, MetabolomicsUSI spectrum link.

**Supplementary Fig. 20:**
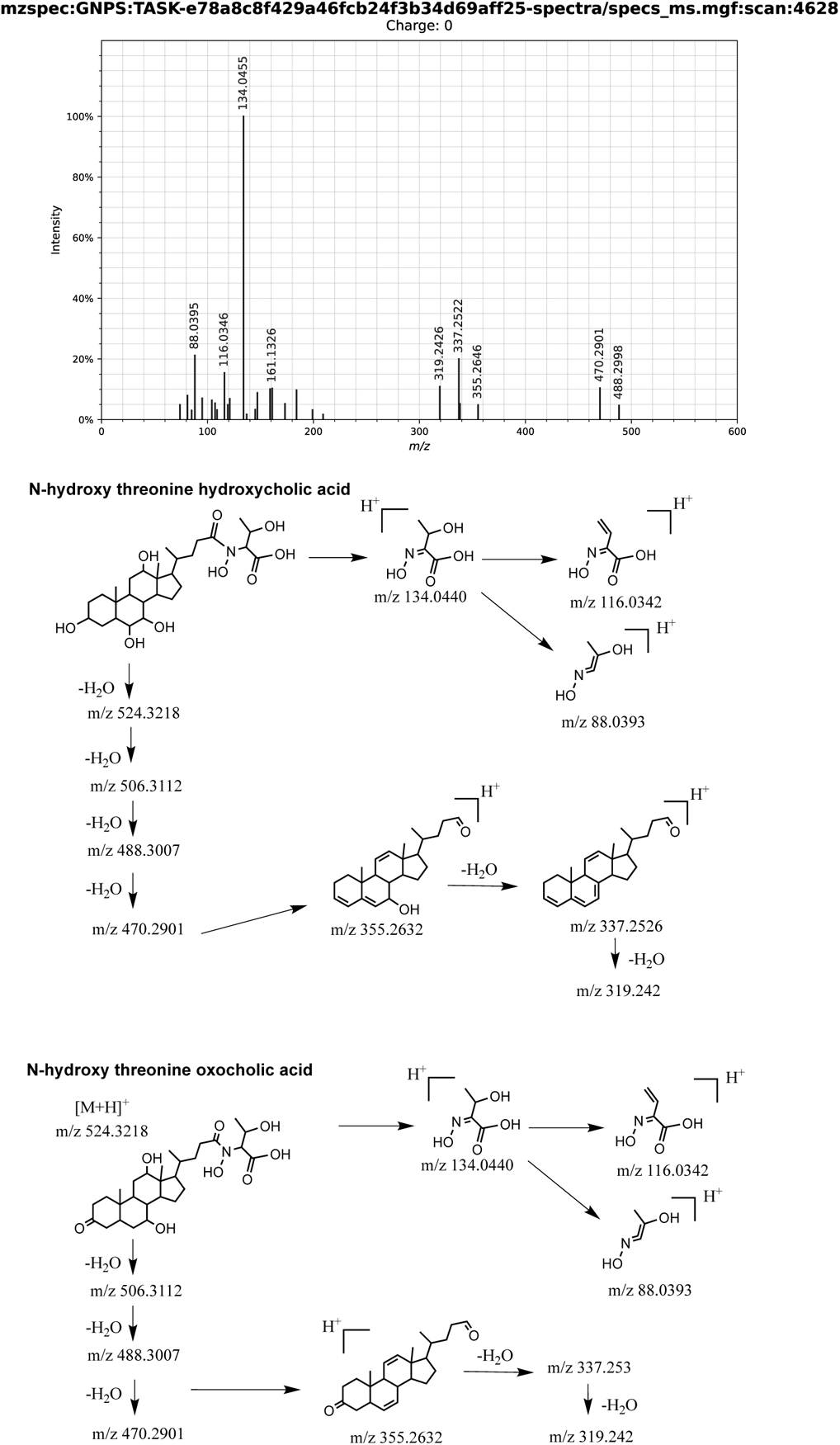
Manual fragmentation analysis of the COSMIC bile acid conjugate 9. Shown is the fragmentation spectrum of *m/z* 524.3222 at 199.57 seconds. Library ID CCMSLIB00005788123, MetabolomicsUSI spectrum link.

**Supplementary Fig. 21:**
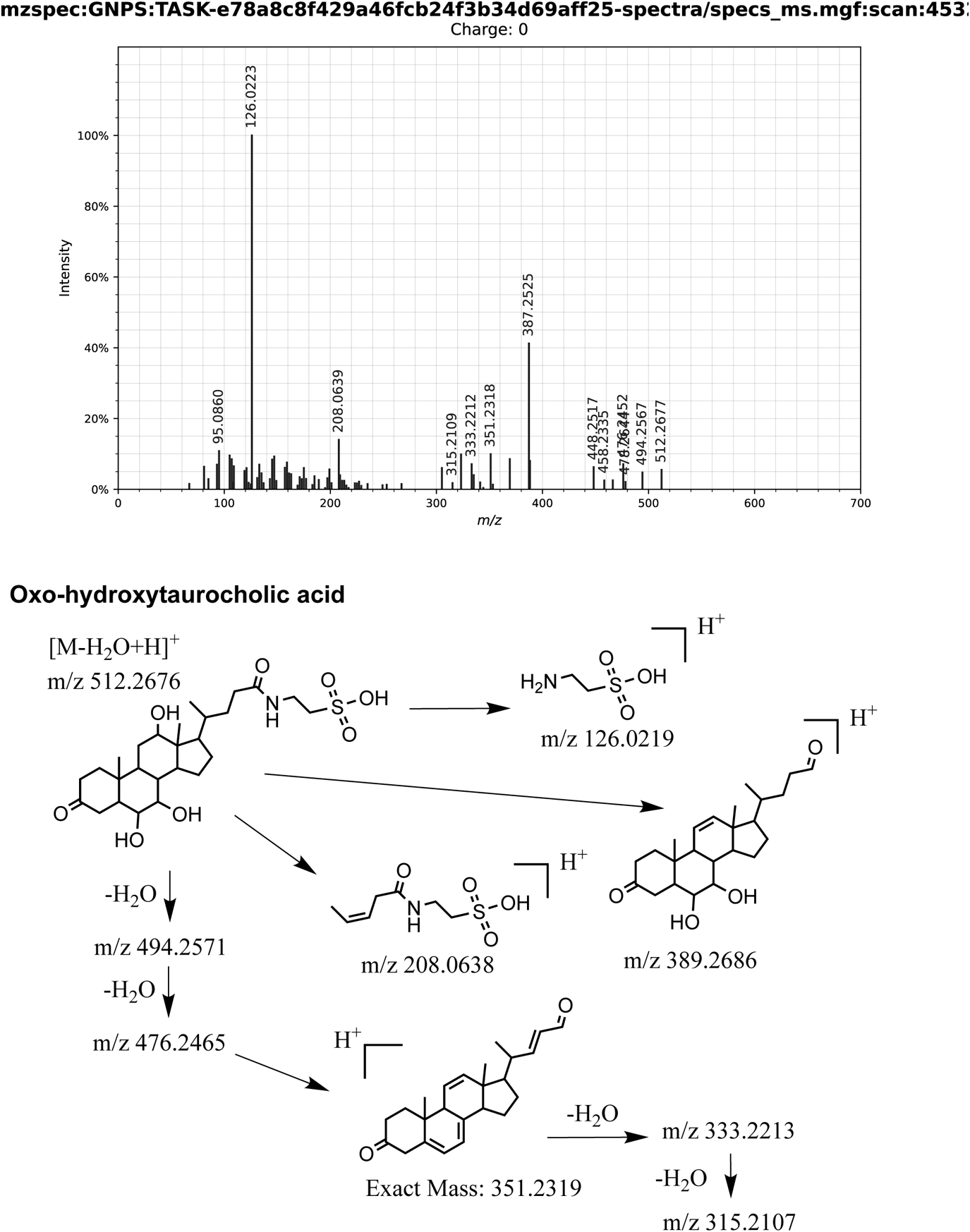
Manual fragmentation analysis of the COSMIC bile acid conjugate 10. Shown is the fragmentation spectrum of *m/z* 512.2691 at 197.38 seconds. Library ID CCMSLIB00005788124, MetabolomicsUSI spectrum link.

**Supplementary Fig. 22:**
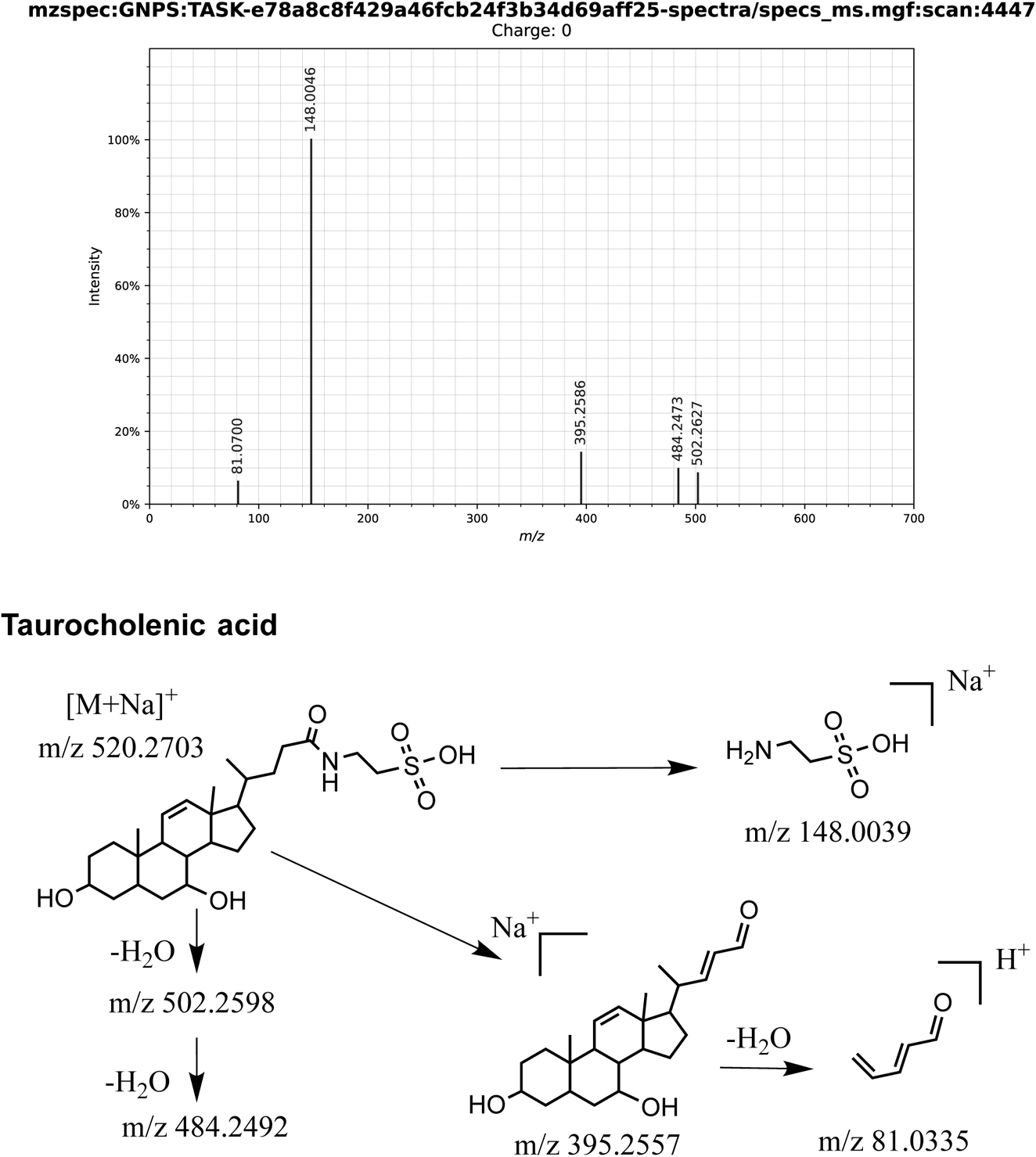
Manual fragmentation analysis of the COSMIC bile acid conjugate 11. Shown is the fragmentation spectrum of *m/z* 520.2728 at 195.1 seconds. Library ID CCMSLIB00005788125, MetabolomicsUSI spectrum link.

**Supplementary Fig. 23:**
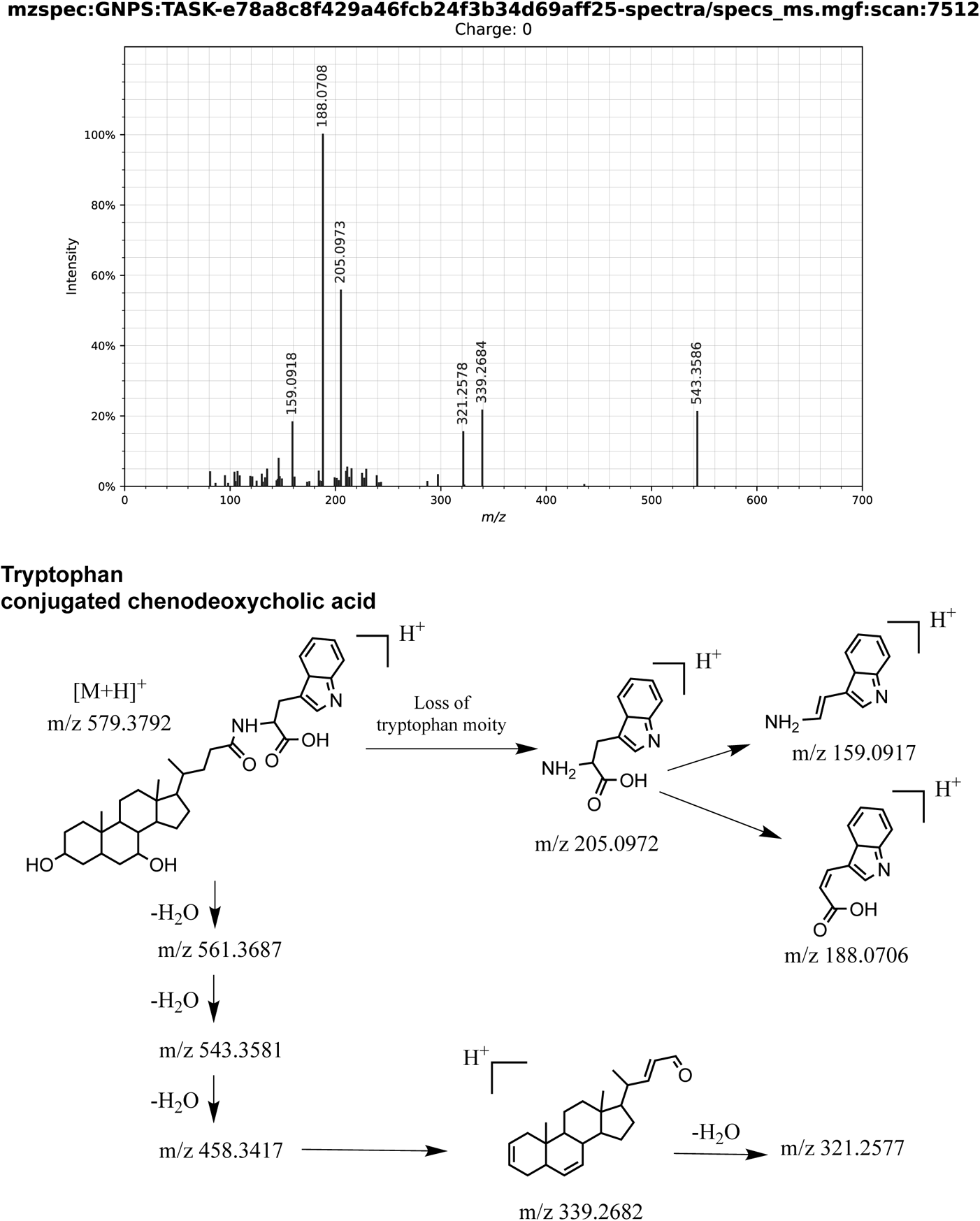
Manual fragmentation analysis of the COSMIC bile acid conjugate 12. Shown is the fragmentation spectrum of *m/z* 579.3788 at 299.1 seconds. Library ID CCMSLIB00005436493, MetabolomicsUSI spectrum link.

**Supplementary Fig. 24:**
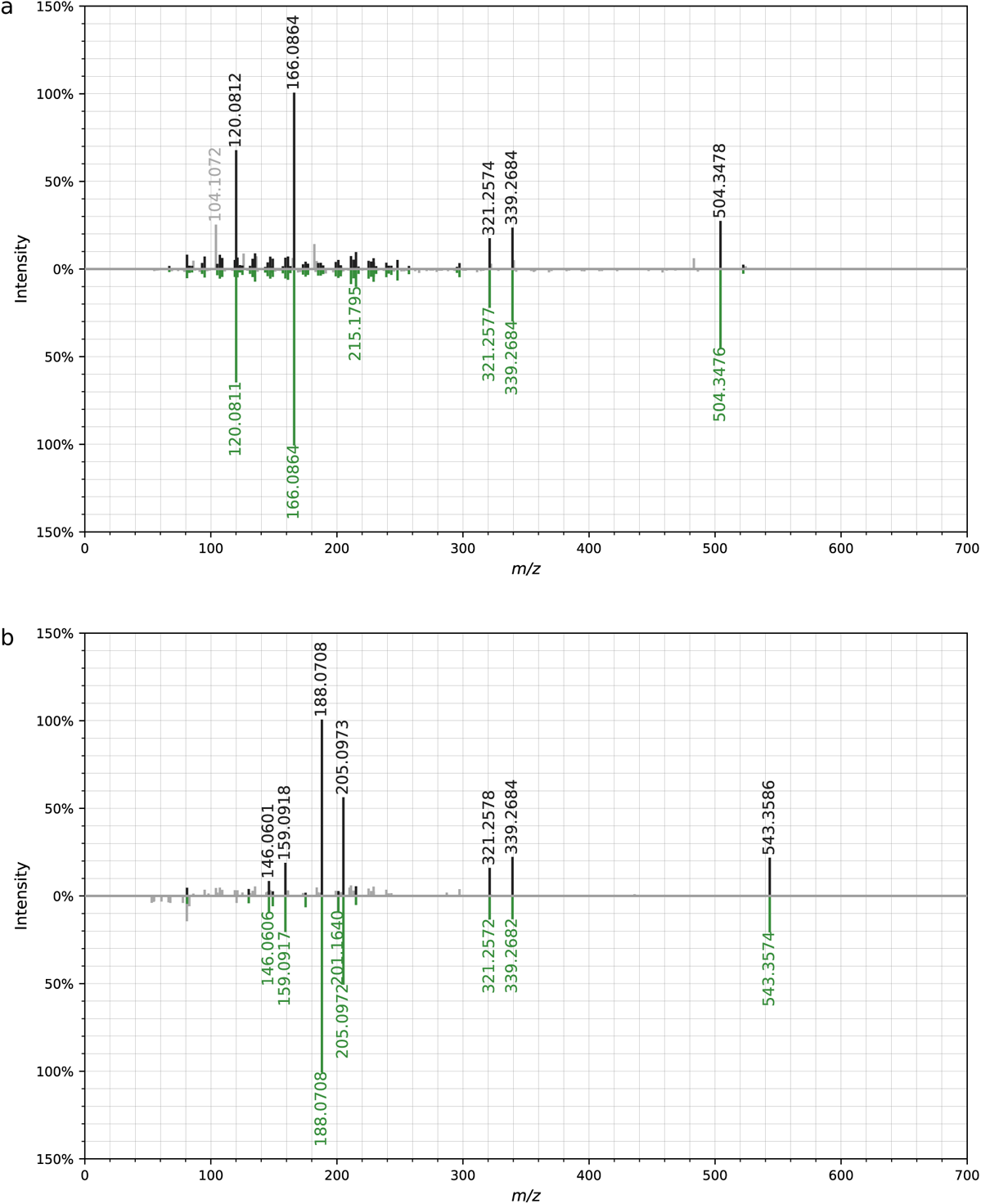
Mirror plots of fragmentation spectra for novel bile acid annotations. Query spectra above, reference spectra below the x-axis. Reference and query spectra of Phe-CDCA 7 (a) and Trp-CDCA 12 (b). Reference and query spectra were both measured on Q Exactive Orbitrap instruments.

**Supplementary Fig. 25:**
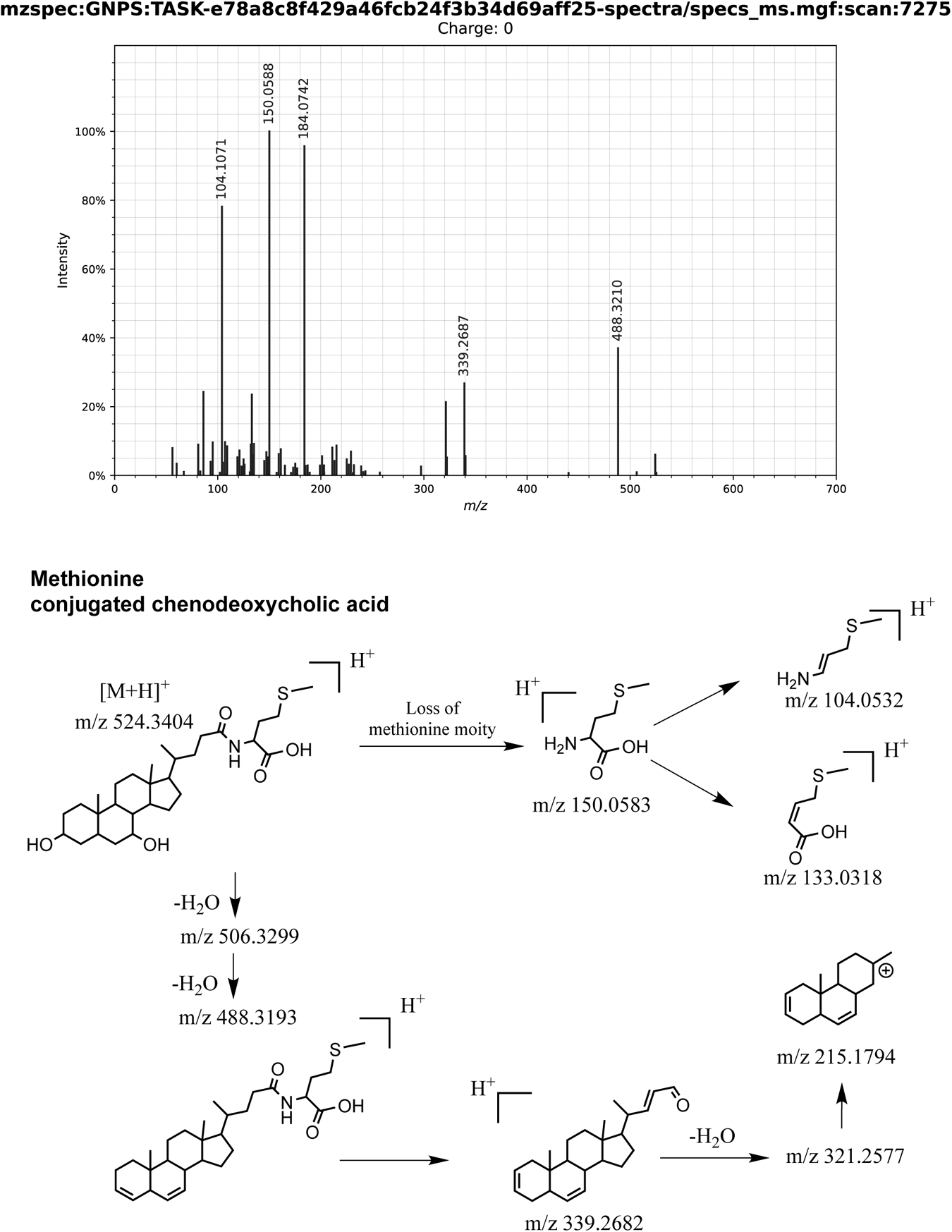
Manual fragmentation analysis of methionine-conjugated CDCA. Shown is the fragmentation spectrum of *m/z* 524.3392 at 291.99 seconds. Library ID CCMSLIB00005725539, [MetabolomicsUSI spectrum link](https://metabolomics-usi.ucsd.edu/spectrum/?usi=mzspec:GNPS:TASK-e78a8c8f429a46fcb24f3b34d69aff25-spectra/specs_ms.mgf:scan:7275#%7B%7D).

**Supplementary Fig. 26:**
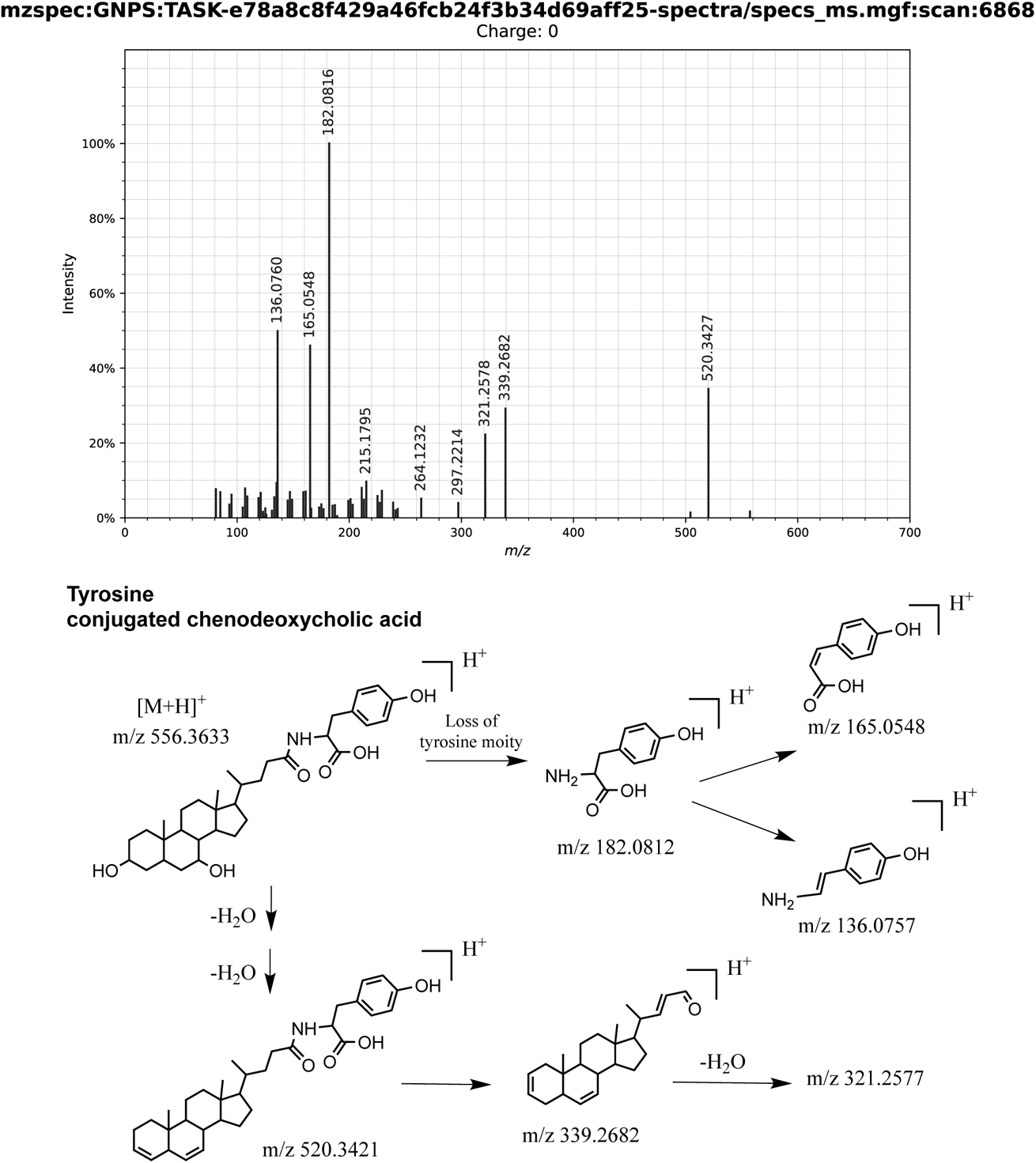
Manual fragmentation analysis of tyrosine-conjugated CDCA. Shown is the fragmentation spectrum of *m/z* 556.3633 at 278.33 seconds. Library ID CCMSLIB00005467948, MetabolomicsUSI spectrum link.

**Supplementary Fig. 27:**
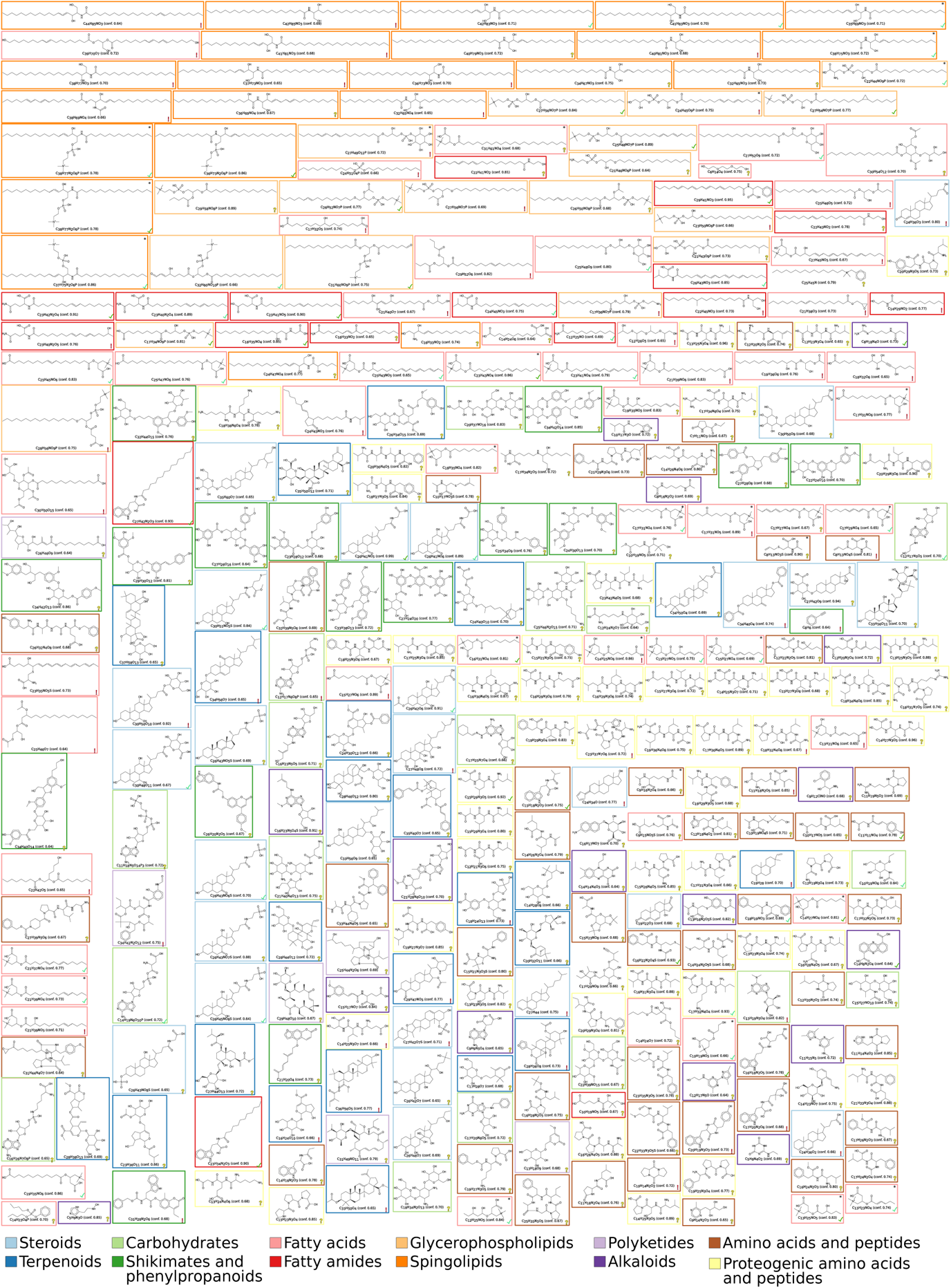
The 315 molecular structure not contained in HMDB annotated with high confidence in the human dataset. Confidence score threshold 0.64 was used. For none of these structures, reference MS/MS data are available. Structures are shown with identification number (ID), molecular formula and COSMIC confidence score. Structures present in the latest version of HMDB (Feb 2021) are marked by an asterisk. Colors indicate compound classes. Notably, 48 compounds were annotated as proteinogenic peptides; these structures were absent from HMDB but are clearly no novel metabolite structures. Lipid structures must be interpreted with some care: It is understood that neither COSMIC nor any other method can deduce, say, the position of the double bond in a carbon chain from MS/MS data alone; rather, this happens to be the candidate present in our biomolecule structure database.

**Supplementary Fig. 28:**
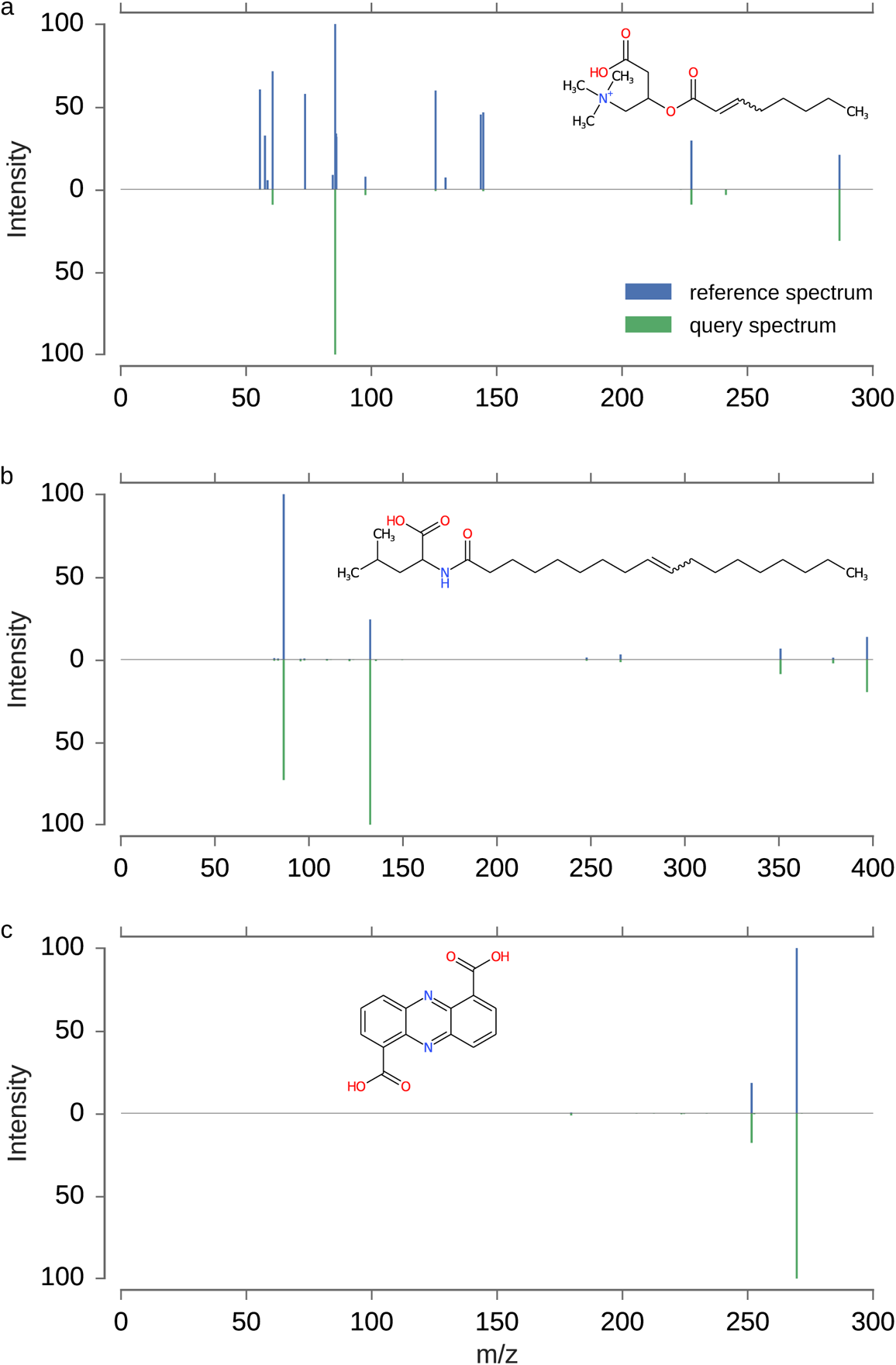
Mirror plots of fragmentation spectra for novel structure annotations in the human dataset. Reference spectra above, query spectra below the x-axis. Reference spectra of 2E-octenoyl-carnitine measured on a Sciex X500R QToF using a collision energy ramp from 20 to 50 eV (a), N-Oleyl-Leucine measured on a Bruker maXis UHR-ToF using a collision energy of 10 eV (b) and phenazine-1,6-dicarboxylic acid measured on a Bruker maXis UHR-ToF using a collision energy of 10 eV (c). All query spectra were measured on Q Exactive Orbitrap instruments with 30 eV (a) and 21 eV (b,c), respectively. Recall that “novel structure” refers to the fact that no training data for CSI:FingerID or COSMIC were available for this structure.

**Supplementary Fig. 29:**
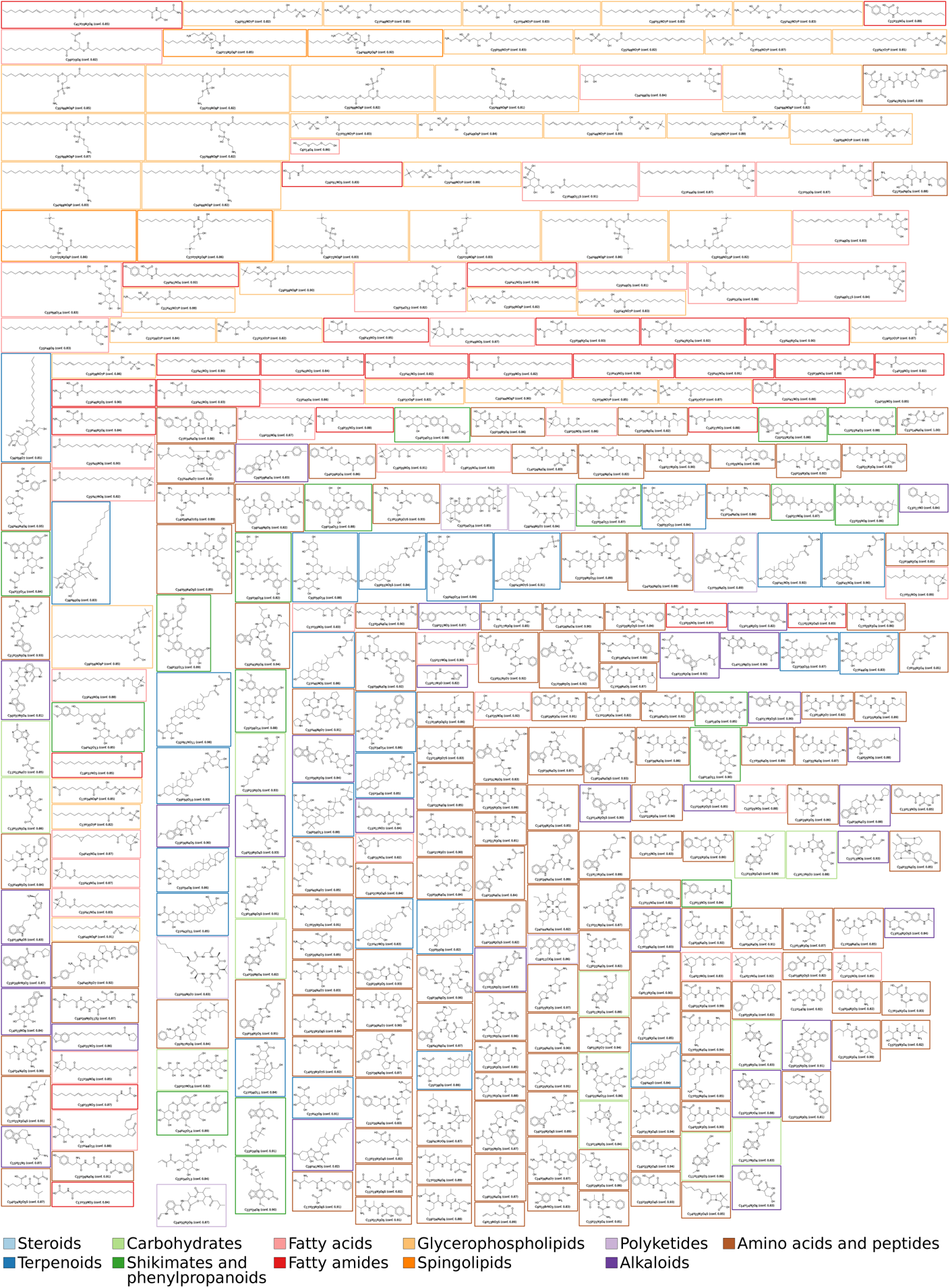
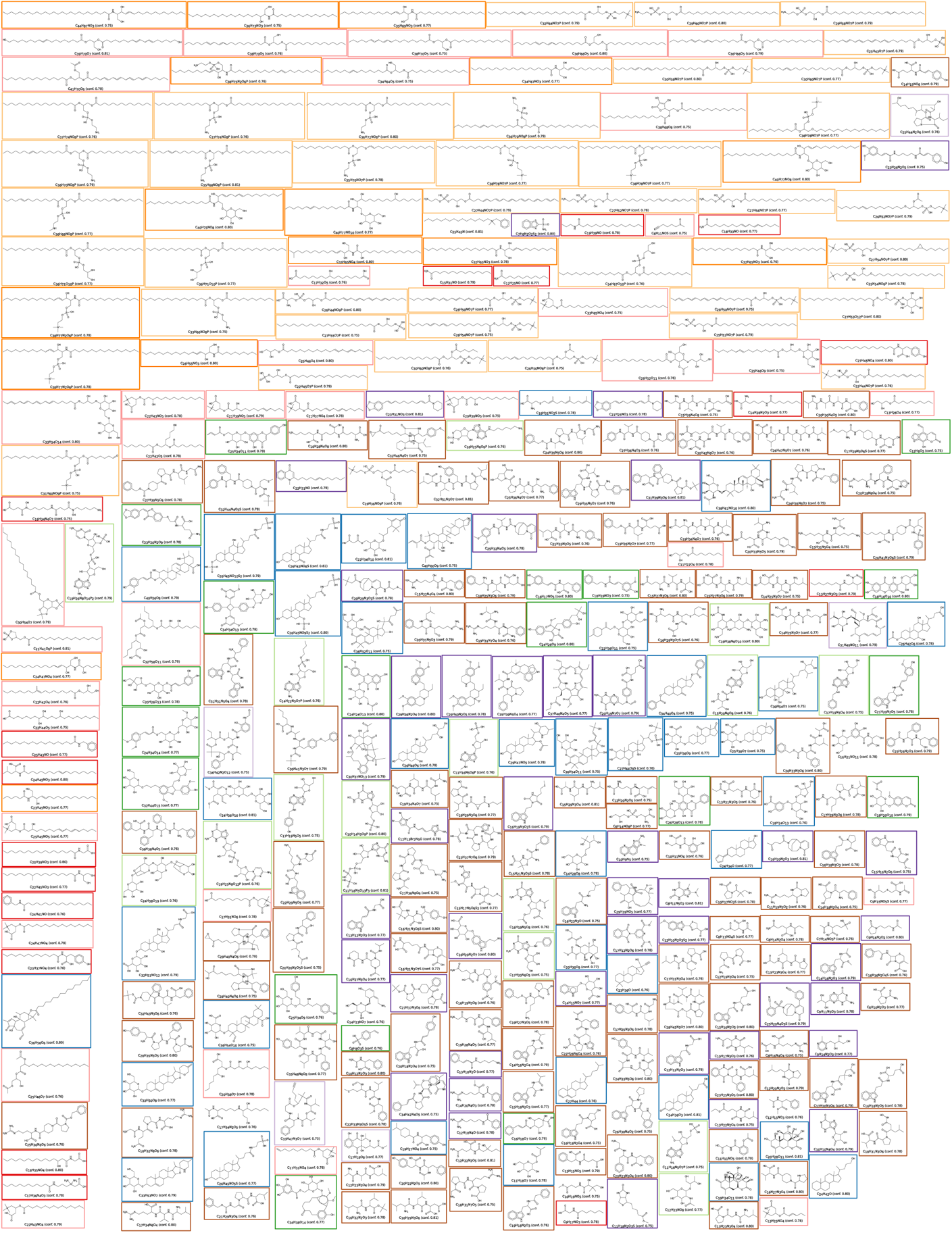
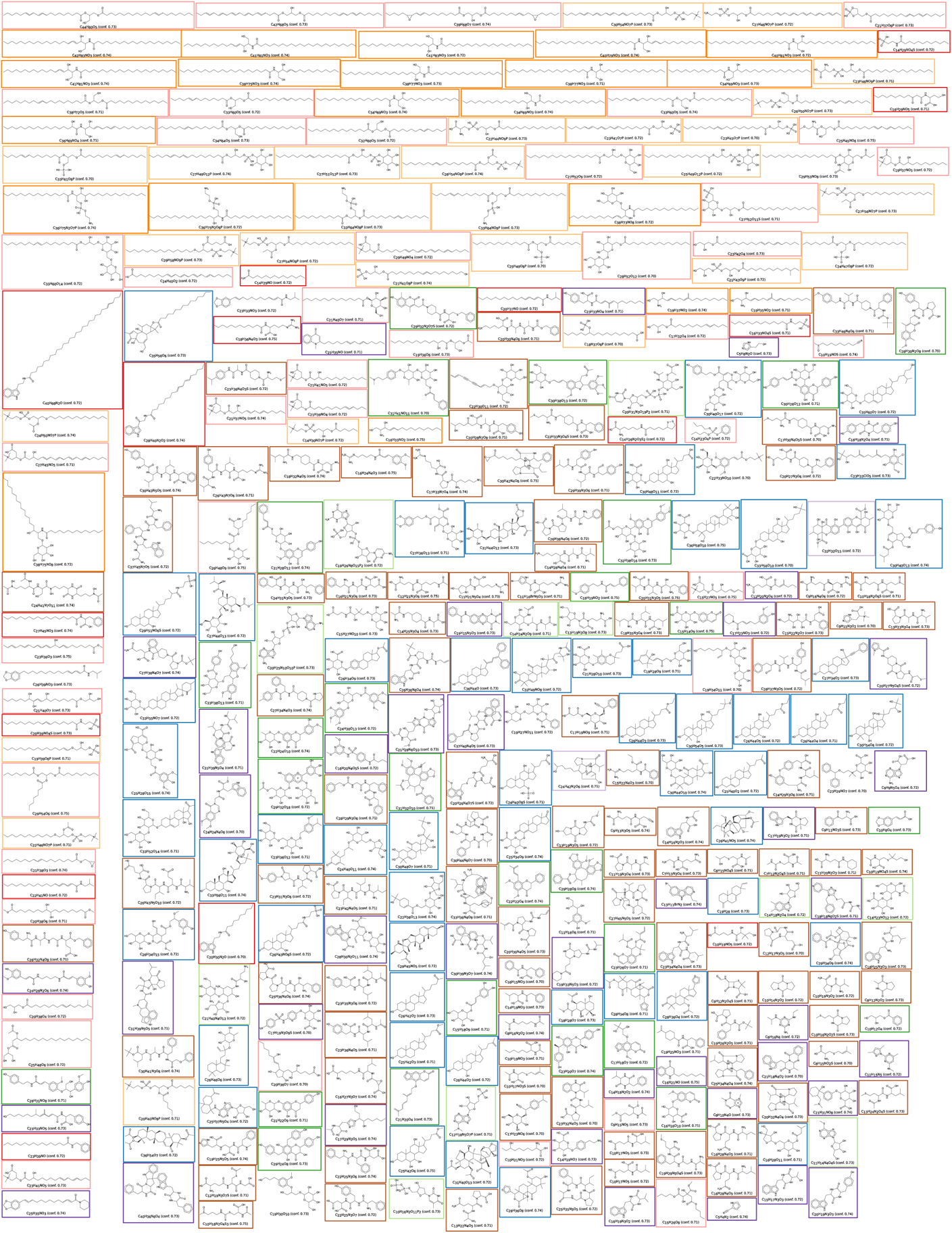
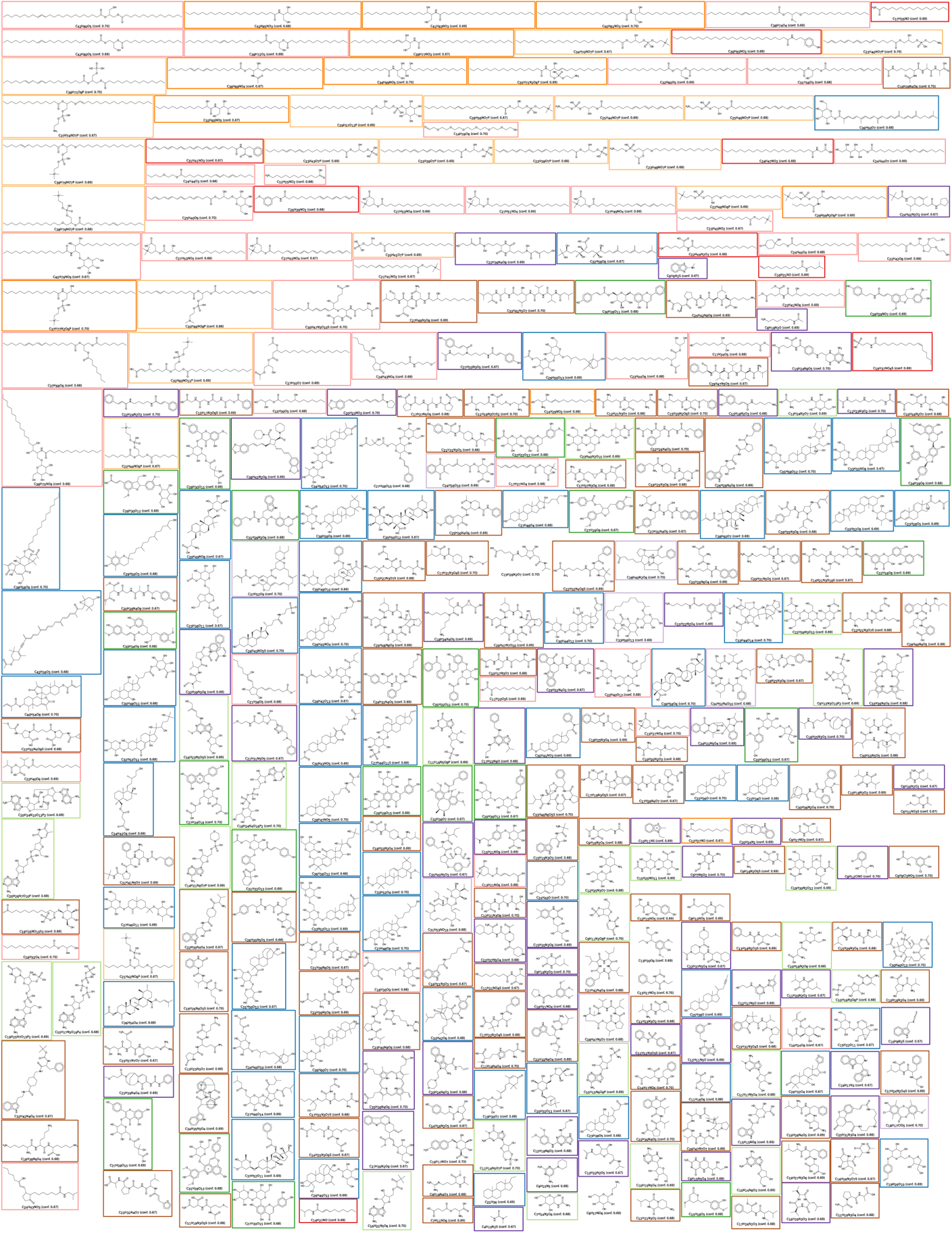
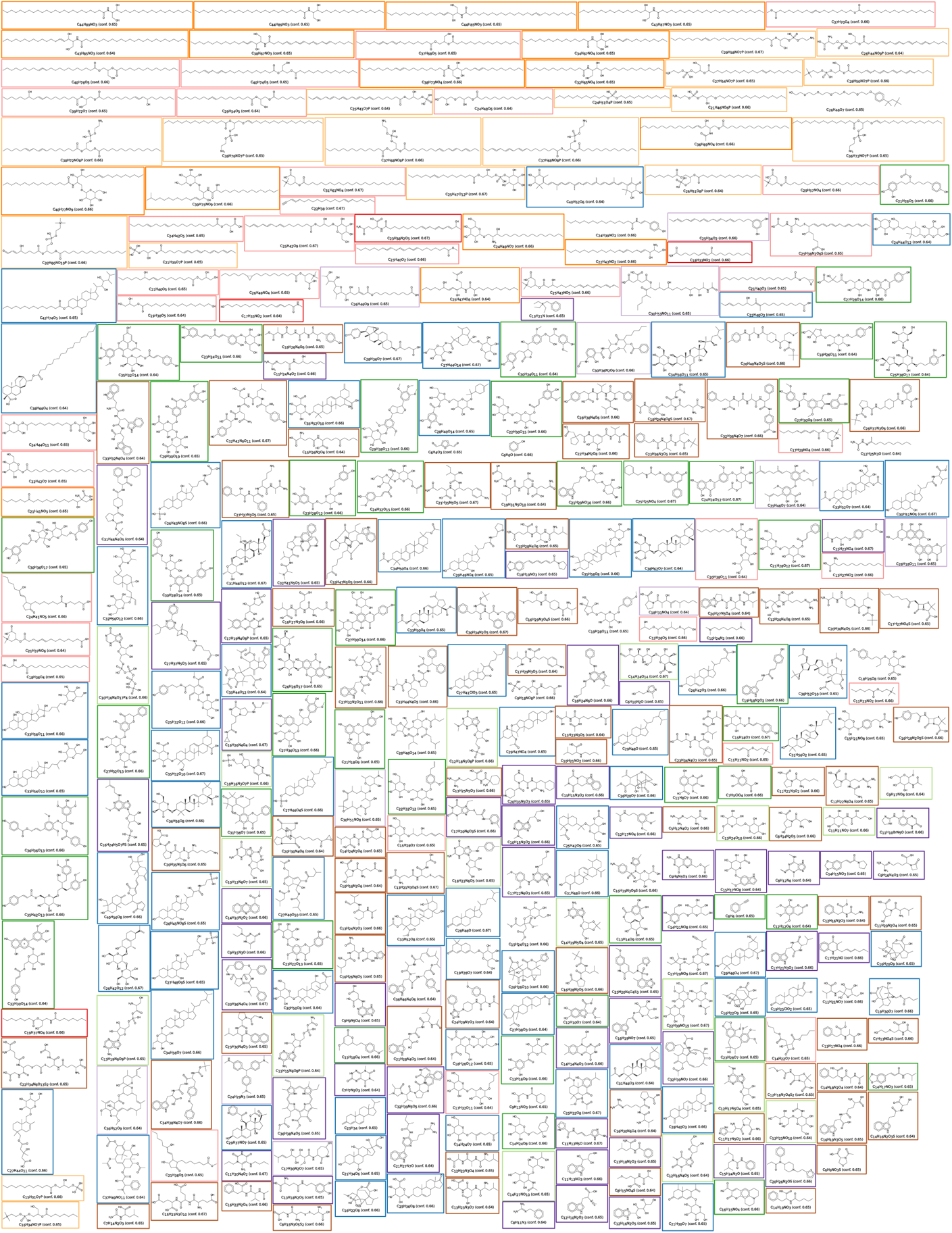
The 1,715 novel molecular structures annotated with high confidence in the Orbitrap dataset. Confidence score threshold 0.64 was used. Structures are shown with identification number (ID), molecular formula and COSMIC confidence score. Colors indicate compound classes. Lipid structures must again be interpreted with some care.

## Notes

https://bio.informatik.uni-jena.de/cosmic

### Competing Interest Statement

S.B., K.D., M.L., M.F., and M.A.H. are co-founders of Bright Giant GmbH. P.C.D. is scientific advisor for Sirenas LLC, Galileo, Cybele and is scientific advisor and co-founder of Enveda and Ometa.

